# Ecology of prophage-like elements in *Bacillus subtilis* at global and local geographical scale

**DOI:** 10.1101/2024.07.03.601884

**Authors:** Polonca Stefanič, Eva Stare, Valentina A. Floccari, Jasna Kovac, Robert Hertel, Ulisses Rocha, Ákos T. Kovács, Ines Mandić-Mulec, Mikael Lenz Strube, Anna Dragoš

## Abstract

Prophages account for a substantial part of most bacterial genomes, but the impacts on hosts remain poorly understood. Here, we combined computational and laboratory experiments to explore the abundance, distribution, and activity of prophage elements in *Bacillus subtilis*. NCBI database genome sequences and isolates from 1 cm^3^ riverbank soil samples were analyzed to provide insights at global and local geographical scales, respectively. Most prophages in wild *B. subtilis* isolates were related to mobile genetic elements previously identified in laboratory strains. Some large groups of prophages were closely related to completely uncharacterized yet functional Bacillus phages, or completely unknown. As certain prophage groups were unique to local isolates, we explored factors influencing prophages within a single genome. Phylogenetic relatedness was a slightly better predictor of host prophage repertoire than geographical origin. We show that cryptic phages can play a major role in acquisition and/or maintenance of other prophage elements both via strong antagonism or by co-dependence. Laboratory experiments showed that most predicted prophages may be cryptic, since they failed to induce under DNA-damaging stress conditions. Interestingly, the magnitude of stress responses remained proportional to the total number of prophage elements predicted, suggesting their importance in host physiology. This study highlights the diversity, integration patterns, and co-occurrence of prophages in *B. subtilis* and their potential impact on host evolution and physiology. Understanding these dynamics provides insight into bacterial genome evolution and prophage-host interactions, laying the groundwork for future experimental studies on the roles of phages in the ecology and evolution of this bacterial species.

## Introduction

Bacteriophages (phages) are the most widespread biological agents on planet Earth ^1^. They not only act as bacterial predators, but can also profoundly impact host bacteria when integrated in their genome. During lysogeny, the host genome integrates phage elements, called prophages, that can influence a wide array of bacterial traits such as virulence ^2,3^, dormancy ^4^, interference with neighbors ^5^, resistance to lytic phages ^6^, metabolic pathways ^7,8^ and cell-cell communication ^9–11^. The effects of prophage elements on their bacterial hosts can further influence higher organisms colonized by lysogenized bacteria, which can translate to global phenomena such as disease outbreaks ^12,13^ or mating patterns in insects ^14^. Given the key role of prophages and their presence in nearly 70% of all bacterial genomes ^5,15^, it is important to understand the effects of particular prophage elements on their bacterial host, their global distribution, potential for transmission, and factors controlling their abundance, transmission, and stability within host genomes.

The abundance of prophage-like elements differs across bacterial phyla, genera, and species ^5,15,16^, as well as within species ^17,18^. Interspecies diversity at the level of mobile genetic elements (MGEs), including prophages, hampers scientific progress in understanding the role of prophages in a broader ecological context because, even for well-established model species of medical, agricultural, or industrial relevance, most experimental studies are performed on a few well-characterized representative strains including *Bacillus subtilis* 168 ^19,20^, *Escherichia coli* K-12 ^21^, and *Pseudomonas aeruginosa* PAO1 ^22^. Consequently, we mostly explore prophages and MGEs that specifically occur in these laboratory strains, while the significance and dynamics of these mobile elements are not recognized in a broader ecological context.

*B. subtilis* is a versatile soil-dwelling bacterium capable of thriving in a wide range of environments, including terrestrial and aquatic environments, as well as humans and other animals. *B. subtilis* is widely used in the industrial production of enzymes and antibiotics^23^, controlling plant diseases and promoting plant growth^24^, improving gut health^25–27^, boosting immunity in mammals ^28^, and promoting overall wellness of humans and animals ^29–31^. *B. subtilis* species form highly resistant spores that can withstand exposure to UV, heat, desiccation, and lack of nutrients ^32^. Furthermore, *B. subtilis* can travel via air over long distances through dust storms ^33,34^. Despite global mixing, a recent study revealed that the relatedness of *B. subtilis* strains is driven by habitat heterogeneity, and sustained at different spatial scales within the environment^35^. Niche-driven diversification seems to be faster than the rate of dispersal, but how habitat-driven diversification affects the acquisition dynamics and evolution of the phage arsenal is unknown.

As a member of the Bacillota (previously Firmicutes) phylum, *B. subtilis* is known to be rich in prophage elements ^16^. Recent experimental studies demonstrated the impacts of these elements on intraspecific competition ^5,36^, dormancy ^4^, surface colonization ^37^, and genome stability ^36,38^. It was also shown that certain prophages can be antagonized by other MGEs, such as conjugative plasmids ^39^, and that the presence of the latter can correlate with prophage diversification ^38^. Furthermore, prophages and defective phage plasmids of this species impair the genetic competence of *B. subtilis* ^40,41^. The above findings suggest that prophage elements play an important role in host adaptation, and that the presence of certain MGEs may influence the stability of prophages within host genomes. However, only a small number of *B. subtilis* temperate phages and prophage-like elements have been described and experimentally characterized, and those examined have been mostly derived from laboratory strains^38,42,43^. As a result, we know very little about the general abundance, phage activity, and ecological relevance of already described temperate phages, or the presence, diversity, and ecology of other, unrelated prophages in non-domesticated isolates. Furthermore, the influence of environmental niche on the phage repertoire of *B. subtilis* strains remains unknown.

This work investigated prophages and prophage-like elements within *B. subtilis* strains at global (publicly available genome sequence databases) and local (our collection of soil isolates) geographical scales. We confirm that *B. subtilis* species are rich in prophage elements, and that these elements integrate into a few specific genomic hotspots. Based on encoded protein similarity, we clustered prophages of *B. subtilis* into dozens of groups, some globally distributed and others restricted to local geographical areas. We also demonstrate clear positive or antagonistic interactions in terms of genomic co-occurrence between certain prophage elements that likely drive their diversification. As isolates from global and local scales showed similar levels of prophage diversity, we experimentally tested the potential of prophages from our local *B. subtilis* strain library for horizontal transmission. Only a minority of prophages could be activated with a common inducing agent. Our work highlights the ecological dynamics of prophage elements within a bacterial species based on public genome sequence database information and a locally collected isolate library, with insights into the genetic interactions (coexistence, exclusion) between MGEs within host genomes.

## Results

### *B. subtilis* species are rich in prophages mainly located in nine integration hotspots

We predicted prophages in a large set of *B. subtilis* genomes, including all 191 complete genomes available on NCBI (now referred to as global-scale isolates) and 40 in a recently sequenced isolate library from a defined 1 cm^3^ soil sample from the Sava riverbank ^44^ (Figure 1A), referred to as local isolates. Collectively, these genomes contained 1243 prophages (Supplementary data 1) and although each genome contained at least one prophage element, most strains carried up to four prophages. Genomes from local scale strains had a slightly different distribution of predicted prophages, with the mode shifted towards a higher average number of prophages (five) per genome, but up to a maximum of eight (Figure 1B). Among global-scale strains, 35 carried 10 or more prophages (Figure 1B). Next, we plotted the predicted prophages according to their integration position in the host chromosome, revealing the presence of at least nine prophage-integration hotspots distributed across the chromosome. The integration hotspots nearly overlapped between local and global-scale isolates (Figure 1C, Supplementary Data 1). The abundance of prophage elements varied among identified hotspots, with the highest abundance of prophages at hotspots near the replication terminus and the lowest abundance close to the origin of replication (Figure 1C). Moreover, the sizes of prophage elements differed depending on their respective integration position. For example, relatively small prophages (averaging 14.6 ± 2.8 kb) were predominantly found at position ∼4 Mb, while two hotspots around position ∼2 Mb exhibited a bimodal distribution of prophage sizes (Figure 1C), with a distinct group of large prophages exceeding 120 kb (Figure 1C, Supplementary data 1).

**Figure 1:**
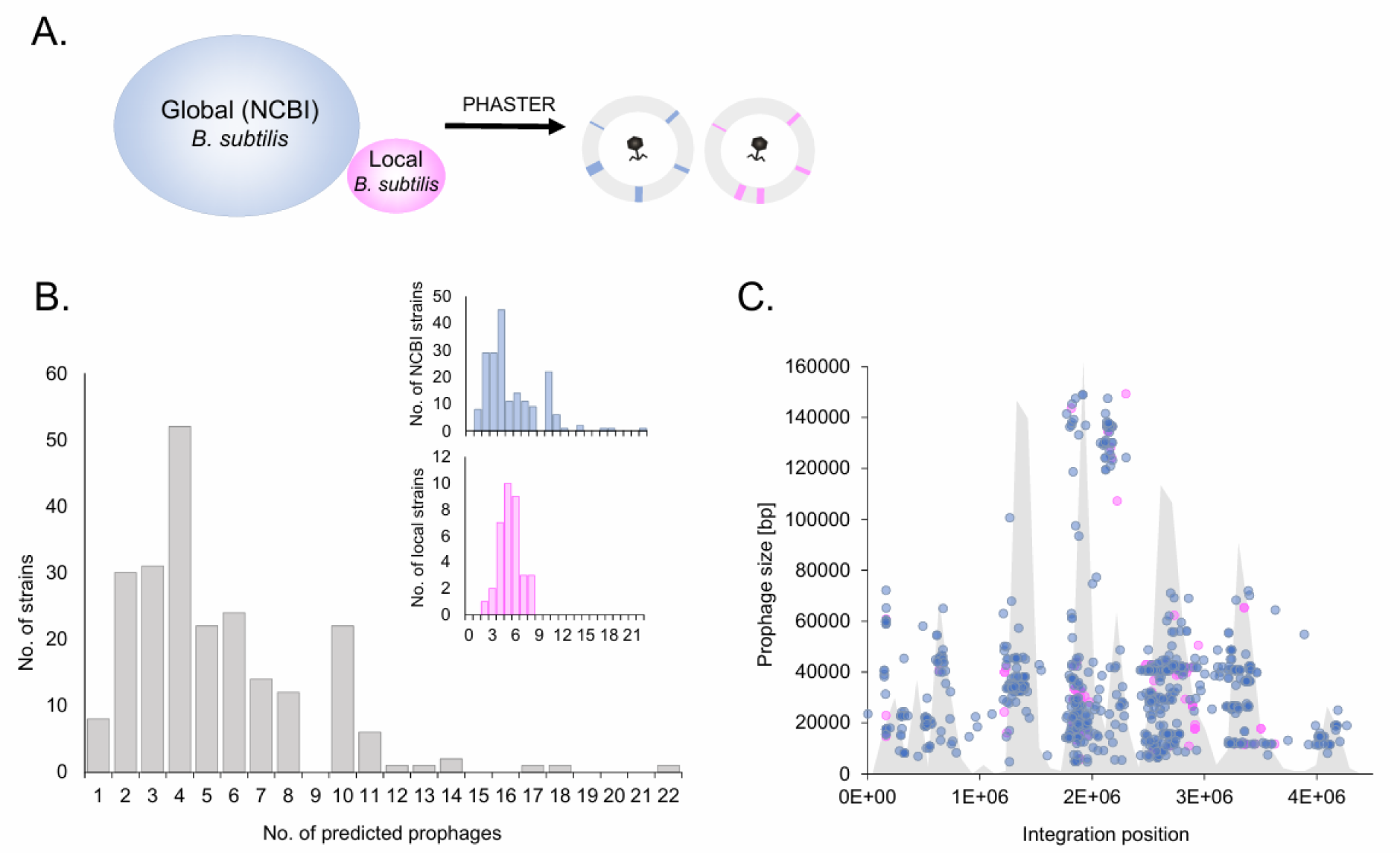
Predicted prophage elements and prophage integration hotspots in *B. subtilis*. A. *B. subtilis* strains from NCBI isolated from a variety of geographical locations and sources, and local scale *B. subtilis* isolates from the Sava riverbank, were analyzed using Phaster software to predict prophage elements. B. Histogram showing the distribution of prophage elements in analyzed *B. subtilis* genomes. In the right corner, global scale NCBI isolates (blue) and local isolates (pink) are represented separately. Chromosomal prophage elements are plotted according to integration position (x axis) and genome size (y axis). Data obtained for NCBI isolates are presented in blue and data for local isolates are presented in pink. The filled histogram in gray depicts the distribution of prophage elements across genomes to indicate the locations of integration hotspots.

### Most prophages are associated with known *B. subtilis* phages or MGEs, but two prophage groups are unique to local geographical scale isolates

Based on total prophage numbers, fragment sizes, and integration sites, broad phylogenetic diversity of prophage elements was expected, which led us to use vContact2 ^45^ to build a local phylogeny for our phage collection. This network derived from vContact2 was subjected to clustering using clusterONE, resulting in 62 vOTUs (Supplementary Data 1−2), which were subsequently used as vContact2-derived operational taxonomic units (vOTUs) in downstream analysis. To determine the taxonomy of the predicted prophages, we used discontinuous mega-BLAST to match them to the INPHARED phage database ^46^ of confirmed active phages, which we supplemented with sequences of previously described MGEs within *B. subtilis* 168 (see Methods). Finally, prophages from vOTU that did not have a match within the aforementioned references, were additionally compared against the genome of *B. subtilis* 168 without MGEs (see Methods) to check for matches to annotated *B. subtilis* chromosomal regions that are not functional prophages. This approach allowed us to classify most of the predicted prophages according to their relatedness to known functional phages or previously described *B. subtilis* MGEs, and according to the predicted phage activity (Figure 2A−C, Supplementary data 2).

**Figure 2.**
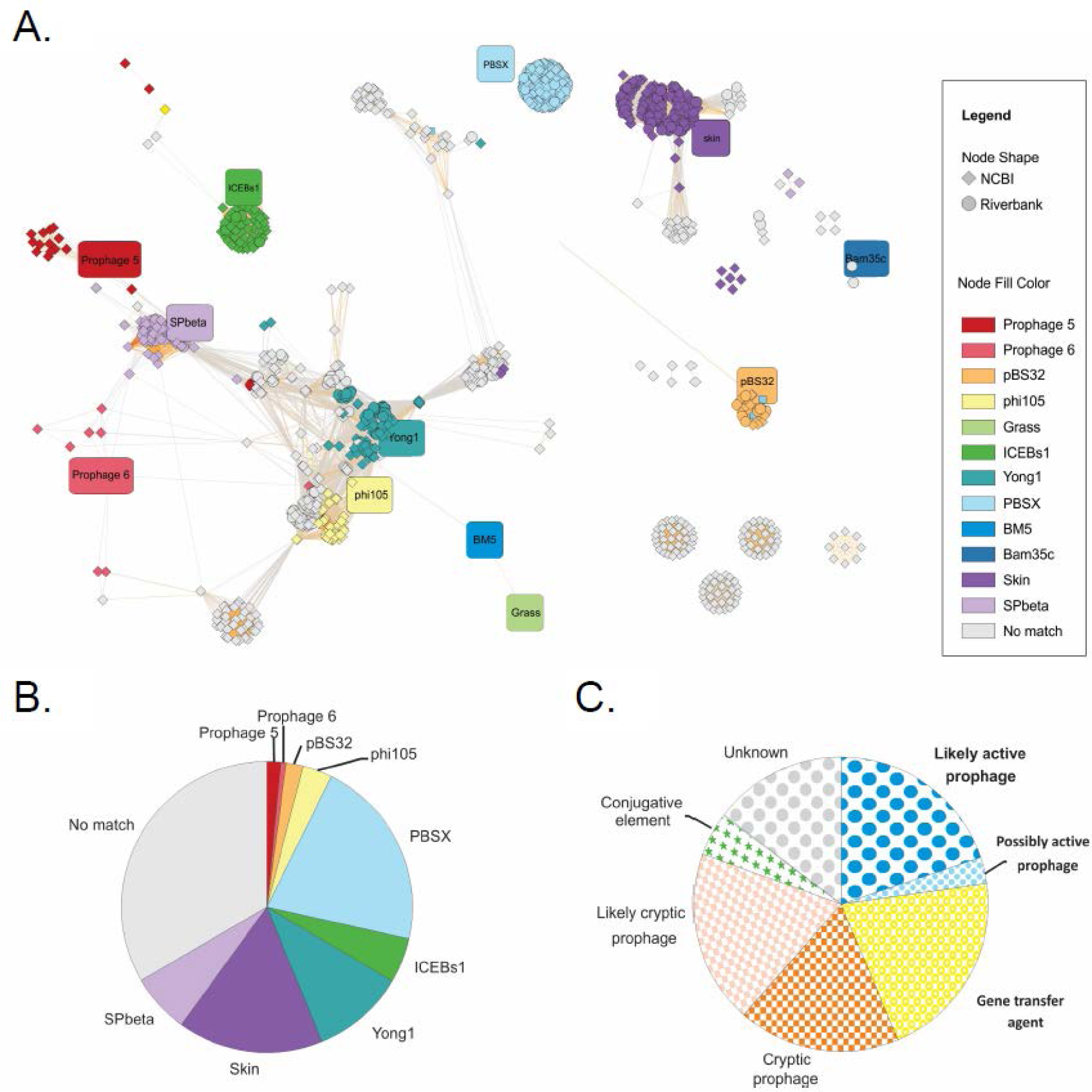
Clustering of predicted prophage elements. A. Prophages colored according to clustering. Reference phages or known mobile genetic elements (MGEs) are depicted as round squares with each color representing one phage cluster. Prophages found within genomes available from the NCBI are shown as diamonds, while prophages from riverbank soil local-scale samples are shown as circles. B. Distribution of predicted prophage elements according to reference matching via nucleotide BLAST. C. Distribution of predicted prophage elements according to their predicted function, based on similarity to active phages, other MGEs, and known *B. subtilis* 168 chromosomal regions. Phages were classified according to the known function of their reference match. In addition, phages matching *B. subtilis* 168 non-MGE chromosomal regions with >30% coverage and >88% identity were classified as likely cryptic. Prophages with lower coverage (<30%) or with no match to references were classified as unknown.

The most prominent group of predicted prophages (n = 238) was phylogenetically related to defective phage PBSX, followed by prophages similar to cryptic prophage *skin* (n = 182), recently isolated phage Yong1 (n = 116), and temperate phage SPꞵ (n = 74; Figure 2A−B, Supplementary Data 2). Other relatively abundant groups were prophages with similarity to conjugative element ICEBs1 (n = 56), previously described temperate phage phi105 (n = 37), a cryptic prophage found on pBS32 plasmid present in *B. subtilis* 3610 (n = 21), or cryptic prophage elements 5 (n = 17) and 6 (n = 7) which are also present in *B. subtilis* 168 (Figure 2A−B, Supplementary data 2). While PBSX, *skin*, ICEBs1, and pBS32 clusters are distant from each other and from other vOTUs, clusters of SPꞵ, element 5 and 6, phi105, and Yong1 are genetically interconnected, which suggests a substantial sharing of proteins between most phages with little phylogenetic conservation (Figure 2A). In addition to taxonomy, the vContact2 network revealed substantial differences in proteome conservation between phage groups, with some groups (e.g. PBSX and ICEBs1) being rather conserved, and others (e.g. *skin*, Yong1, phi105, and SPꞵ) being more diverse, displaying lower similarity within vOTU, and even grouped into several different vOTUs (Supplementary Data 2, Supplementary Figure 1).

We also observed that prophage elements that clustered together tended to reside in similar integration sites within *B. subtilis* genomes (Supplementary Figure 2). However, there were certain exceptions, where phages classified into different phylogenetic groups were found at similar chromosomal locations (e.g. *skin*, Yong1, and phi105-related prophages). Prophages of these two groups were among the less conserved in terms of chromosomal position, as they were found in multiple different chromosomal positions (Supplementary Figure 2). The latter prophage is probably linked to several vOTUs with relatedness to *skin* and phi105, and may be located at different attachment sites. Finally, we gained further insights into larger clusters of completely unknown prophages, such as vOTU 8 that always seem to integrate near Yong1, vOTU 0 that always integrates close to the origin of replication, and vOTU 16 that may replace the *skin* element in certain genomes (Supplementary Figure 2).

A minor portion of *B. subtilis* strains also carried plasmids, and a minority of those also contained prophages, which clustered into a few different vOTUs, entirely unrelated to all other phages (Figure 2A). Finally, certain relatively large clusters were found (e.g. vOTU 8, 16, 0, and 40), which had no matches with known phages or MGEs, or matched poorly with *B. subtilis* 168 chromosomal regions. These were assigned as completely unknown (Figure 2A−C, Supplementary Data 2). Based on similarity to references and chromosomal regions, we classified ∼23% of predicted prophages as likely or possibly active, ∼63% as cryptic, likely cryptic, or other MGEs that are not functional phages, and ∼15% as completely unknown (Figure 2C, Supplementary Data 2).

Finally, we compared prophage clustering networks of database (NCBI) and local-scale bacterial strains, specifically asking whether there are any distinct prophage groups associated solely with strains from local-scale samples. While the vast majority of prophage groups found among riverbank isolates were represented also within global-scale isolates, we found groups that were unique to local-scale isolates, specifically, two plasmid prophages assigned as outliers associated with the Bam35c reference bacteriophage, and a unique cluster of Yong1-related phages (vOTU64) that were unique to local-scale isolates (Figure 2A).

### Specific host phylogenetic clades are defined by distinct sets of prophages

To better understand the connection between host phylogeny and the similarity of prophage elements, a phylogenetic tree based on all analyzed *B. subtilis* strains was constructed using core gene alignment (see Methods). *B. subtilis* strains were clustered into seven phylogenetic clades (PCs), with local strains belonging to only two clades (PC1 and PC2), which were also more closely related to each other than to other clades (Figure 3). Plotting prophage elements to the phylogeny of strains revealed that phylogeny correlated with the presence of specific prophage elements to a certain extent (Figure 3). This was confirmed after we quantified the variability of the prophage repertoire within (0.96−0.915) and between (0.955−0.893) the largest clades (Supplementary Data 3, Figure 3). For all except PC3, prophages were more similar within clades than between clades when strains from all four clades were included (Figure 3). Clade PC3 was rather unique, as it was characterized by the highest diversity of prophage elements (0.915) and by strains carrying a distinct set of prophage elements compared to other strains (Figure 3). Interestingly, PC3 strains were mostly missing the most abundant prophage vOTU 5 (PBSX), and carried vOTU 6 (also PBSX) instead. There was no obvious distinction in phage repertoire between clusters containing fewer strains (clusters PC5−PC7). It was difficult to estimate the effect of local environment on the diversity of prophage repertoire among isolates, mostly because local strains clustered into PC1 and PC2, which also exhibited rather high similarity of prophages between clades (Figure 3). Nevertheless, within local strains, prophage similarity score (0.942) was lower than prophage similarity scores within PC1 (0.96) and PC2 (0.943), suggesting that phylogeny was a better predictor of prophage repertoire than strain isolation site. This was also in line with a rather random distribution of countries and isolation sites across the phylogenetic tree (Figure 3). Together, these results indicate that closely related strains tend to carry similar prophages.

**Figure 3.**
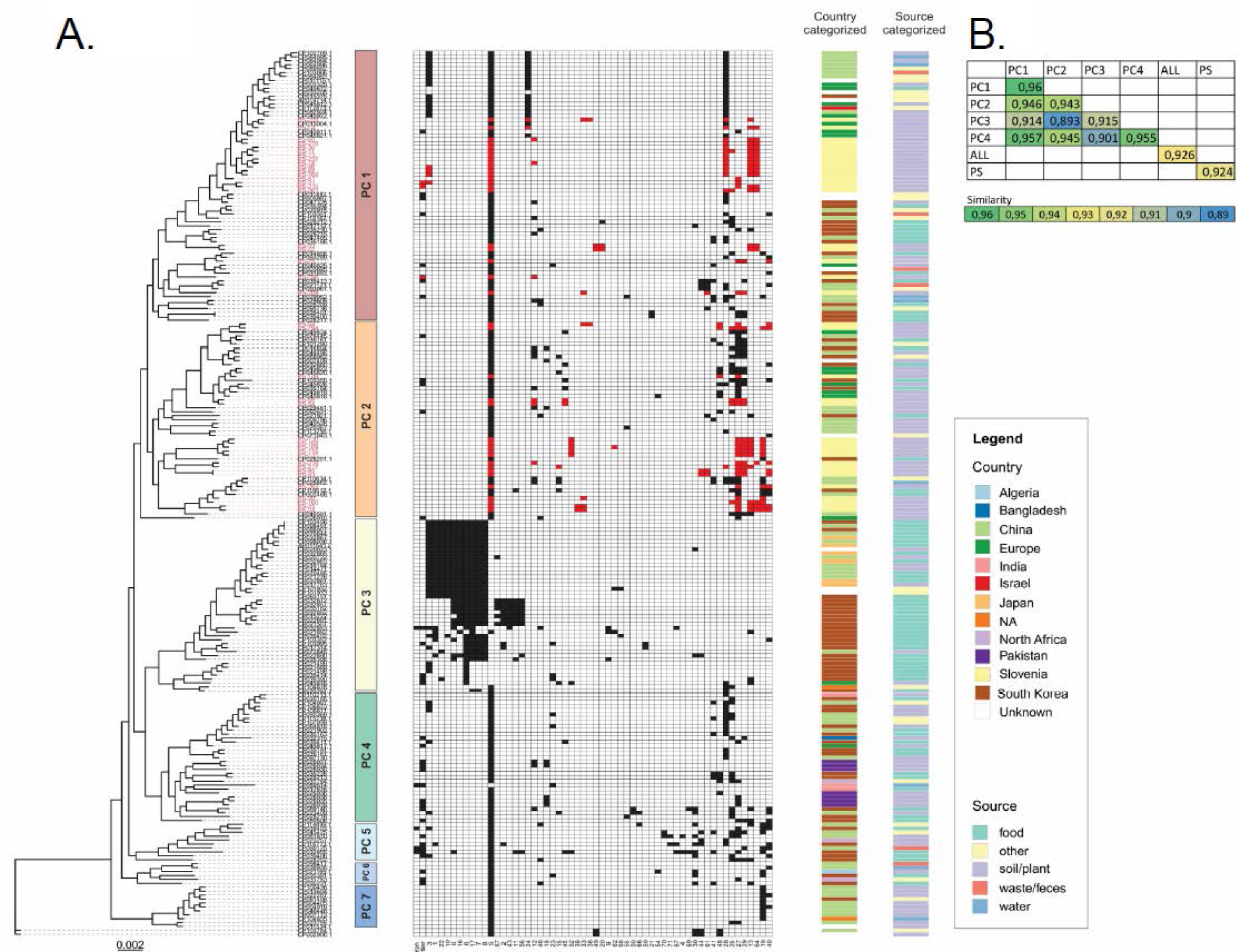
Relationship between phylogeny, prophage cargo, and the country/isolation source of *B. subtilis* strains. A. From the left to right: phylogenetic tree illustrating the phylogeny (core gene alignment) among all analyzed *B. subtilis* strains; phylogenetic cluster demarcation, with each color representing a different phylogenetic clade; matrix of presence/absence of prophage elements (red indicating the presence of local strains, black indicating the presence global strains, and an empty space indicating absence) associated with specific clusters, country of origin, and the source of strain isolation. B. Table showing prophage presence/absence similarity scores calculated within or between phylogenetic clusters or within local-scale isolates (see Methods). Briefly, similarity score within strains for each phage vOTU was calculated, and scores of all vOTUs were averaged to obtain similarity at the level of the entire prophage repertoire (see also Supplementary Data 3).

### Both mutual inclusion and exclusion of prophages is common within bacterial genomes

Next, we tested the hypothesis that prophage elements shape the prophage repertoire of the host, for instance, by either preventing or promoting acquisition of prophages. In particular, this was suggested by several unique prophage elements within PC3 (Figure 3), that seemed to co-occur within genomes belonging to this phylogenetic clade. Indeed, our analysis revealed both negative and positive co-occurrence between predicted prophage elements (Figure 4, Supplementary Data 4, Supplementary Figure 4).

**Figure 4.**
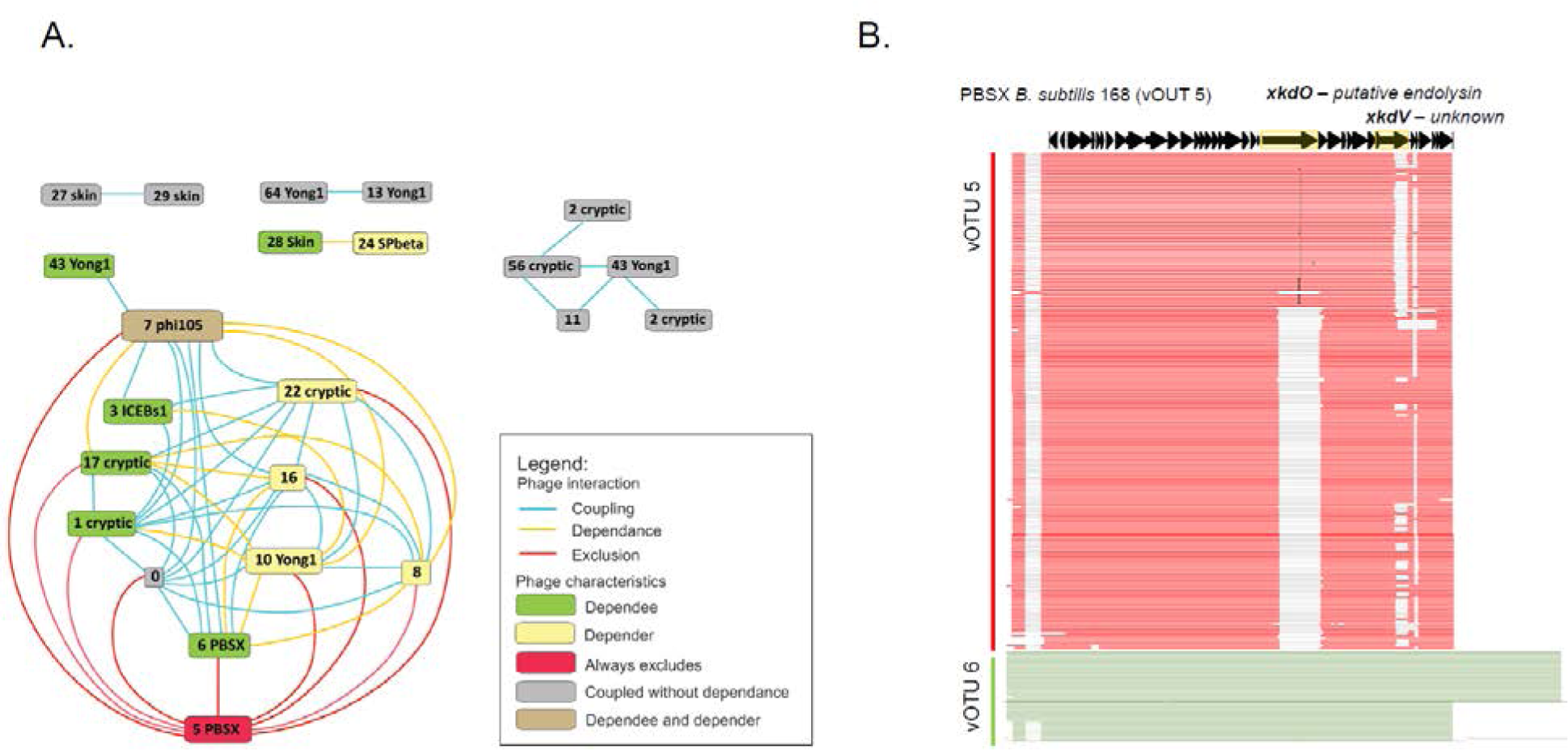
Map of prophage interactions and characteristics considering their co-occurrence within a single host chromosome. A. Prophage vOTU labels colored according to their relationships with other vOTUs within a single chromosome: dependee - prophages of other vOTUs tend to occur only when this vOTU is present; depender - this vOTU occurs only in the presence of another vOTU; always excludes - if this vOTU is present, certain other vOTUs never occur; coupled without dependence - tend to co-occur with another vOTU; depender and dependee - its presence depends on a certain vOTU, but also determines the presence of another vOTU. Lines connecting squares are colored according to interactions: coupling (blue), exclusion (red), dependence (yellow). B. Graphic summary of alignments of all PBSX genomes from vOTU 5 (colored red) and vOTU 6 (colored green) obtain by Blast. Map of PBSX from *B. subtilis* 168 [NC_000964.3] was shown above the alignment. The two regions that are commonly distinct between vOTU 5 and vOTU 6 overlap with open reading frames of *xkdO* and *xkdV* of PBSX *B. subtilis* 168.

Among all vOTUs, roughly a third (21 of all 62 vOTUs) were involved either in positive or negative co-occurrences with other prophage elements within the host chromosome. From all types of interdependencies detected, most vOTUs showed coupling (statistically significant co-occurrence of both vOTUs within the host chromosome), followed by dependence (where the presence of one vOTU depends on the presence of another, which is not dependent on the first), and exclusion (where a certain vOTU is never found in the presence of another vOTU), as shown in Supplementary Figure 4. Prophage elements that seemed to have the largest effect on the overall prophage arsenal were PBSX vOTU 5, which showed exclusion towards nine other vOTUs; PBSX vOTU 6, which showed coupling with five vOTUs, and determined the presence of another three vOTUs; and phi105 vOTU 7, vOTU 0, and Yong1 vOTU 10, which also showed a significant number of positive interactions (Figure 4, Supplementary Figure 4, Supplementary Data 4).

To delve deeper into the factors contributing to these correlations, we focused on two groups of PBSX elements, vOTU 5 and vOTU 6, which never co-occurred within the host chromosome, and which exerted a profound impact on the presence/absence of other prophages (Figure 4, Supplementary Figure 4). PBSX of vOTU 5, encompassing ∼83% of all identified PBSX prophages, exhibited a strong antagonistic relationship with PBSX in cluster 6 (constituting ∼17%) and with phage phi105. Consequently, if an isolate carried PBSX of cluster 5, it was highly unlikely to be lysogenic for phi105 vOTU 7, Yong1 vOTU 10, and a few other unknown or likely cryptic prophages. Conversely, PBSX of vOTU 6 displayed a strong positive correlation with these prophage elements (Figure 4, Supplementary Figure 4). Furthermore, we noticed coupling between certain types of SPbeta prophage and the *skin* element (Figure 4, Supplementary Figure 4).

Given that PBSX of vOTU 5 is the most widespread prophage, we hypothesized that protection from attack and lysogenization by multiple other phages may select for its maintenance within the species. An interesting observation in this prophage was the existence of an alternative PBSX variant differing in the cell wall hydrolase *xkdO* gene region and the end fragment of the hypothetical protein-encoding gene *xkdV* (Figure 4B). These results clearly show that the likelihood of acquisition or loss of certain prophage elements can be predicted based on the presence or absence of other specific prophages.

### Strains with more predicted prophages show stronger DNA damage responses, but only some prophages appear to be active

Previous experimental data suggest that the majority of computationally predicted prophage elements are non-functional temperate phages^47^, meaning they are not capable of forming phage particles and propagating via a lytic cycle. To challenge bioinformatic prophage prediction with experimental data, we tested 40 local isolates from the soil microscale for spontaneous or stress-induced release of active phage particles using mitomycin C. We found that prophages could be induced in 6 of 40 *B. subtilis* strains (PS-11, PS-24, PS-25, PS-233, PS-237, and PS-261) producing titers in the range of 10^2^−10^7^/ml using *B. subtilis* Δ6 (ΔSPβ Δ*skin* ΔPBSX Δprophage 1 pks::cat Δprophage 3) as an indicator strain, and in 2 of these 6 strains the induction was spontaneous (without mitomycin C treatment), as indicated by the presence of plaques on the lawn of the indicator *B. subtilis* strain Δ6 (Figure 5A).

**Figure 5.**
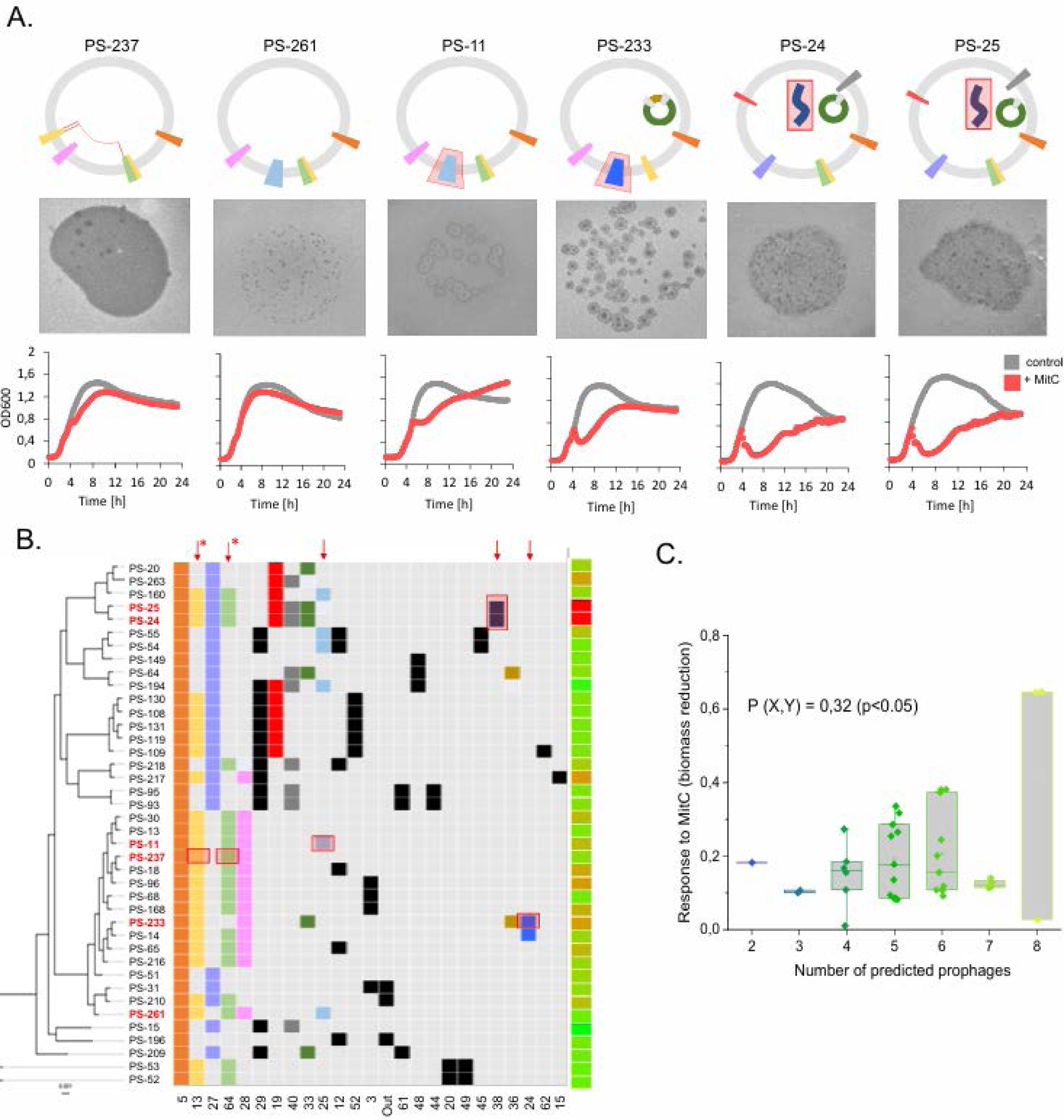
Screening for active prophages within local-scale isolates. A. Of 40 strains treated with mitomycin C, six showed the presence of active phages in their supernatants (PS-237, PS- 261, PS-11, PS-233, PS-24, and PS-25). Prophage elements predicted within their genomes are schematically labeled on circular graphs. Elements that were identified as active by phage DNA sequencing as labelled in red. Graphs below plaque images represent growth curves with and without addition of mitomycin C (0.1 μg/ml). Results are averages of six biological replicates and error bars represent standard error. B. Phylogenetic tree of local strains, with the presence of prophage elements that belong to certain similarity clusters represented by squares of different colors. Supernatants of strains labeled in red produced plaques on the indicator strain when collected post prophage induction with mitomycin C. The heatmap on the right represents the strength of response to mitomycin C, measured as biomass reduction. Active prophage elements as identified by phage DNA sequencing were labelled by red rectangles and entire vOTU within which active prophages were identified, was labelled by red arrow. Arrows with asterisk signify partial coverage of phage DNA reads. C. Number of predicted prophage elements versus strength of response to mitomycin C measured as biomass reduction compared to non-treated controls. P stands for Pearson correlation coefficient. Boxes represent the first and third quartiles. Lines represent the median and bars span maximum to minimum values. All data points are overlaid on boxes.

Spontaneous induction rates were too low to determine active prophages from sequencing coverage (Supplementary Figure 5). Therefore, phage DNA was extracted from post-induction lysates and subjected to Illumina sequencing (see methods). The obtained sequencing reads clearly mapped onto predicted prophage sequences within the genomes of PS strains: in case of PS-24 and PS-25, reads mapped onto linear phage plasmids (vOTU 38), for PS-11 and PS-233 onto SPβ-related elements (vOTU 25 and vOTU 24, respectively) and for PS-233 they distributed among Yong-related elements vOTU 13 and vOTU 64 (Figure 5A, Supplementary Figure 6). As our previous analysis suggested positive association between vOTU 13 and vOTU 64 (Figure 4A), this data strongly suggest co-dependence of these two prophage elements and assembly of composite phage.

Regardless of plaquing on the indicator strain, we also tested how all 40 strains responded to the optimized concentration of mitomycin C (0.1 μg/ml), which triggered strong biomass reduction mainly in strains carrying active prophages (Supplementary Figure 7). However, not all strains that carried predicted active phages also showed strong biomass reduction in the presence of mitomycin C (Figure 5B) and *vice versa*. For example, strains PS-18 and PS-96 showed strong biomass reduction but did not produce active phages (Supplementary Figure 8). In theory, strains PS-18 and PS-96 (and other strains which showed strong responses to mitomycin C but did not plaque) could release phages that are not specific to the indicator strain. Nevertheless, there was no relationship between host phylogeny and detection of plaques on the indicator strain (Figure 5B), suggesting that a growth response to mitomycin C does not always reflect the presence of active prophages in the host genome.

Next, we compared the total number of predicted prophages based on the strength of the response to mitomycin C (calculated as the reduction in biomass compared to untreated controls; Figure 5C). Strains with more predicted prophages within their genome were found to respond more strongly to mitomycin C (Figure 5C). These results indicate that although most of the predicted prophage elements were no longer able to transmit and replicate in a new bacterial host, they still triggered cell lysis in response to DNA damage. The results also show that vOTU membership cannot predict activity of prophage element (e.g. majority of strains carrying vOTU 25 did not show prophage activity) and that even elements with high induction rate can be restricted to only minority of strains within microscale (e.g. prophages of vOTU 38). The results also suggest that coupling detected between certain prophages within single host chromosome (Figure 4A) can be due to assembly of composite phages from two closely related and distantly integrated prophages upon lytic cycle induction.

## Discussion

Here, we assessed prophage elements, their diversity, and potential interactions within complete *B. subtilis* genomes available on NCBI, and within a collection of *B. subtilis* strains isolated from 1 cm^3^ of riverbank soil. Recently, prophage elements were assessed in 194 *B. subtilis* genomes available on NCBI ^48^, where diversity analysis was limited to prophage elements categorized as intact ^48^. In our approach, we ignored the functionality categories assigned by the software to remove many relevant MGEs (such us gene transfer agents, or plasmids with phage features) from the analysis. Genomic comparison identified 55 distinct types of prophages with varying genome sizes, integration sites, and gene content, as well as several groups of prophage elements with homology to temperate phi105 ^48^. Our protein-based clustering analysis confirms these results, as well as the presence of two groups of SPbeta-like phages.

We found that strains from local geographical scale samples had a prophage frequency distribution distinct from that of global isolates, and carried certain unique groups of prophage elements. This supports previous experimental studies that demonstrated rapid acquisition and spread of temperate phages from and within local microbiomes ^49^. Our work also clearly demonstrates barriers against phage transmission; even strains that coexist within the local scale, and are closely phylogenetically related, differ profoundly in the prophage repertoire they carry. Moreover, even though some of these prophages showed high potential for lytic cycle induction and horizontal transmission (like plasmid prophages present in strains PS-24 and PS-25), they were restricted to only 2 of 40 isolates within the local isolate library.

In both global and local isolates, integration of prophage elements was restricted to a few specific genomic hotspots, with increasing numbers of integrated prophages close to the replication terminus. This observation is in line with previous work ^17^ and seems to be a general cross-species phenomenon, likely linked to transcriptional regulation and easier maintenance of stable lysogeny in the chromosomal region where gene expression (including lytic cycle genes) is lowest ^50^.

As most of the present knowledge about *B. subtilis* MGEs is derived from laboratory strain *B. subtilis* 168 and its ancestor strain *B. subtilis* NCIB 3610, we attempted to group the predicted prophage elements according to their similarities to MGEs residing in these strains. Our work confirmed the conservation of defective prophage PBSX in all isolates of local and global scale. The second most abundant group included phage elements that clustered with cryptic prophage *skin*, recently characterized *Bacillus* phage Yong1, temperate phage SPβ, and functional temperate phage phi105. Meanwhile, within natural isolates we found PBSX and *skin* that were identical to those present in *B. subtilis* 168, but no strains carrying full-length Yong1 or phi105 in a prophage form. This may suggest a strong selection for the domestication of this temperate phage within the host genome^51^.

How and why PBSX is strictly maintained within *B. subtilis* is unclear. Although it was successfully deleted from the *B. subtilis* genome and classified as dispensable ^52^, its conservation within species implies a strong ecological advantage. It was recently shown that PBSX is involved in extracellular vesicle formation during stress-induced cell lysis, a process that depends on expression of the PBSX endolysin gene ^53^. Previous studies also suggest that PBSX may be constantly induced in a small subpopulation of cells^54^, which could potentially infer toxicity towards PBSX-negative strains and cause a rapid vertical spread of this element. Importantly, we show that PBSX exists in two taxonomic subgroups, which are strictly exclusive towards each other, and where one strictly antagonizes phi105-related prophage elements. This suggests that this dominant PBSX type could protect *B. subtilis* from other MGEs, like phi105, especially since the latter was classified as a ‘naive’ phage, entering the lytic cycle during *B. subtilis* genetic competence development^41^. This observation is also in line with previous reports on competition between phages for intracellular resources during lytic induction^55^, and superinfection immunity encoded within prophage elements^56^.

In addition to the strict antagonism between the two subgroups of PBSX, we found a strict co-occurrence between certain Yong 1 and phi105 ‘relatives’, as well as unknown and cryptic prophage groups. It is likely that phi105 vOTU 7, vOTU 17 (cryptic), and vOTU 22 (cryptic) are remnants of a single prophage because they have neighboring integration sites (*att* at 3.2 Mb). The fact that they always co-occur with distantly integrated elements such as Yong1 vOTU 10 (*att* at 1.7 Mb) may indicate that all three are needed for maintenance of lysogeny and host survival. Another possibility is molecular piracy, in which one defective element cannot spread without the other helper element^57^ – a hypothesis that is strongly supported by phage DNA sequencing results, indicating that sequences from two topologically distant prophage elements showing statistically significant co-occurrence within single chromosome (both related to Yong1) can indeed assemble into composite phage Interestingly, certain prophage elements that were abundant among wild isolates clustered together with well-described non-prophage regions of domesticated *B. subtilis* 168. For example, elements clustered with conjugative element ICEBs1 were found in 54 strains spanning local and global scale isolates. Although this element was previously shown to block lytic cycle induction in SPβ^58^, we did not find any antagonism or statistically significant patterns of co-occurrence between these two elements in *B. subtilis* genomes.

We also found that some prophage groups present in wild strains partially or almost completely matched chromosomal regions of *B. subtilis* 168. This may either suggest a miss-prediction by the software, or the presence of prophage remnants in the lab strain. Some of these regions, such as those encoding biosynthetic gene clusters, deserve special attention. This suggests that, in addition to sublancin encoded by SPβ, other BGCs may also be transferred within prophage genomes. Finally, we found that certain genomic regions of *B. subtilis* 168, proposed to be prophage regions, had very few or no matches to the collection of natural isolates (e.g. prophage element 1, 2, or 3), strongly suggesting they are not part of functional prophages.

While very few *B. subtilis* strains carried plasmids, many of these appear to carry prophage-like elements. We discovered very potent functional phages on small linear plasmids within PS-24 and PS-25 local scale isolates. Further studies are needed to better understand their role in bacteria and/or host range. Our experimental data support recent findings ^47^ showing that despite the abundance of predicted prophage regions within *B. subtilis* chromosomes, only a fraction are induced by mitomycin C. Our work also shows that the most active phages are not necessarily widespread within the local community. Further studies will show whether the presence of plasmid prophages with a high spontaneous induction rate may provide certain benefits to host bacteria.

Although *B. subtilis* is a model organism and an industrially relevant species, knowledge of the natural diversity of its prophages remains limited. Our study uncovers this diversity and novel interactions between MGEs, as well as their level of conservation at the global scale and their level of uniqueness at the local scale. To what extent these elements drive interactions within the species and adaptation to certain growth conditions, such as fermented food, remains to be discovered. Our work opens multiple possible directions for further experimental research on the molecular mechanisms of antagonistic and positive interactions between MGEs, as well as host range, transmission, and conversion studies on active MGEs.

## STAR Methods

### Bacterial genomes

A dataset of *B. subtilis* genomes consisting of complete genome sequences available on NCBI (ftp://ftp.ncbi.nih.gov/genomes/Bacteria/, last accessed March 2023) was used along with complete genomes of local soil microscale isolates (See below; Supplementary dataset 1). Raw sequencing data for microscale isolates were obtained using various sequencing platforms including PacBio, Nanopore, and Illumina (Nextera and Truseq PCR-free libraries) from single isolate DNA extracts. Detailed information about genome sequences, libraries, and platforms used for each strain can be found in the NCBI database under Bioproject accession number PRJNA437002. Unicycler was used to hybrid assemble genomes using short and long reads^59,60^. NCBI genomes were carefully curated based on metadata, which specifically meant that laboratory strains and their derivatives were excluded, while isolates with data for isolation sites and sources were retained. All genomes were oriented relative to *B. subtilis* 168 (AP011541.2), if necessary, using the ‘linearize’ function of SnapGene (www.snapgene.com). The final dataset contained 191 NCBI genomes and 40 genomes of soil microscale isolates.

### Identification of prophages and integration loci

Prophage coordinates in bacterial genomes were determined using the web-based Phaster tool ^61,62^. As genome orientations were corrected prior to prophage extraction, prophage coordinates were treated as integration loci. Prophages were retrieved from corresponding genomes using a custom-made pipeline and used for further analysis. As references, a collection of sequences consisting of prophages and other MGEs was individually retrieved from *B. subtilis* 168 (NC_000964) and contained the following: prophage elements 1−6 ^63^, conjugative element ICEBs1 ^64^, defective phage PBSX ^65^, *skin* element ^66^, SPꞵ ^67^, and plasmid pBS32 (KF365913.1) known to carry a defective prophage ^68^. In addition, phi105 (NC_048631.1) was added as a reference, since this is a known temperate phage targeting *B. subtilis* ^69^. Finally, *Bacillus* phage Yong1 (OP918669.1) was added as a reference after initial analysis of a substantial number of matches.

### Classification of phages into virus operational taxonomic units (vOTUs)

In order to organize predicted prophages into clusters, vContact2 ^45^ was used. In brief, prophage genes were predicted with Prodigal, translated into proteins, and these were used for reference-free phylogenetic clustering with vContact2, which builds a network based on the number of shared proteins between phages. ClusterOne was then used to infer clusters of related phages based on this network. Reference sequences of selected MGEs of *B. subtilis*, as mentioned above, were manually added to this set. These clusters were then used as vOTUs for further analysis. To further examine the taxonomy of these phages, each phage was aligned to a database of known phages. Here, we used discontiguous megablast (dc-megablast) to match them to the INPHARED phage database ^46^ of confirmed active phages, allowing for large gaps, which was necessary due to the high level of phage recombination. We considered a phage to match a reference if the coverage of the phage was at least 50% at ≥70% identity. Prophages from vOTUs that did not have a match within reference sequences were additionally blasted against the *B. subtilis* 168 genome without MGE sequences to check for potential matches to annotated *B. subtilis* chromosomal regions that do not carry functional prophages.

### Host phylogeny and prophage content

All bacterial host genomes were phylogenetically evaluated using pan-genome analysis. Specifically, bacterial genomes were annotated with PROKKA ^70^ using the *B. subtillis* 168 annotation as a reference, and the resulting annotations were used to build a core genome using ROARY ^71^ by considering genes to be core if they had >95% protein similarity in >99% of genomes. The resulting core gene alignment was then used to construct a maximum likelihood phylogenetic tree with FastTree ^72^ using the generalized time reversible (GTR) substitution rate model. Phylogenetic clusters containing more than five *B. subtilis* subspecies *subtilis* strains were determined arbitrarily.

### Ecological distribution

Information on strain geography and habitat was derived from NCBI metadata. Furthermore, isolation sites were limited to include the most abundant geographical locations (Supplementary dataset 1), and isolation habitats were divided into five main categories: soil/plant, food, waste/feces, water, and others. Host prophage cargo and isolation habitats of hosts were mapped onto a phylogenetic tree using R. The locations of strain isolation were mapped onto the world map using Python 3.10.12 and <u>Open cage Geocoding API</u> (Supplementary Figure 9). Briefly, global strains were isolated from North America, Africa, Europe, and Asia. Local strains were isolated from two 1 cm^3^ soil samples from the riverbank of Sava river in Slovenia ^73^ and have been previously described and phenotypically analyzed^35,44,74,75^.

### Correlations and anticorrelations between the presence of prophage elements within a single host genome

To investigate if some phages are mutually exclusive/inclusive within the same genome, we noted the presence of each vOTU, as defined above, within each host genome, resulting in a binary presence/absence matrix. Logistical regression was then performed to statistically test whether or not vOTUs co-occurred within genomes (i.e. which vOTUs were mutually inclusive or conversely, mutually exclusive).

### Prophage induction and plaque assay

Bacterial strains were routinely cultivated in LB broth (LB-Lennox, Carl Roth; 10 g/l tryptone, 5 g/l yeast extract, 5 g/l NaCl). To assess phage activity in culture spent media, overnight cultures were initiated from -80°C glycerol stocks and cultivated overnight at 37°C with shaking at 200 rpm for ∼16 h. Cultures were transferred to fresh LB medium at 1% inoculum and further cultivated at 37°C with shaking at 200 rpm. In mid-exponential phase (OD_600_ ∼0.5), mitomycin C was added (1.5 μg/ml) to trigger prophage induction, followed by cultivation for another 4 h. Cells were pelleted by centrifugation (8000 g, 10 min), supernatants were filtered-sterilized, and diluted in order to obtain well-separated single plaques. Plaque assays were performed using *B. subtilis* Δ6 (ΔSPβ Δ*skin* ΔPBSX Δprophage 1 pks::cat Δprophage 3) ^52^ as an indicator strain. A similar experiment, but without addition of mitomycin C, was performed to assess spontaneous prophage induction (SPI). The effect of mitomycin C on growth dynamics was also analyzed in a separate assay on a microtiter plate using two different concentrations (1.5 μg/ml and 0.1 μg/ml). The mitomycin C effect was quantified by comparing areas under the growth curves between controls (without addition of mitomycin C) and treated samples, and expressed as % of biomass reduction.

### Phage DNA extraction and sequencing

Phages were propagated on soft agar to obtain at least 20ml of lysates with titer of 10^6^ pfu/ml. Lysates were filter-sterilized and mixed at a 1:4 rate with PEG-8000 solution (PEG-8000 20%, 116 g/l NaCl). After overnight incubation at 4 °C, the solutions were centrifuged for 60 min at 11000 rpm to obtain phage precipitates. The pellets were resuspended in 500µl of SM buffer (5.8 g/l NaCl, 0.96 g/l MgSO4, 6 g/l Tris-HCl, pH 7.5), followed by phage DNA extraction using Genomic Mini AX Phage kit (A&A Biotechnology, Poland), using modified protocol with nuclease incubation step extended to 60 minutes. Phage sequencing was performed by Illumina MiSeq instrument and a 2 × 250 nt paired-end chemistry (MiSeq Reagent Kit v2 (500-cycles). Primary data analysis (base-calling) was carried out with Bbcl2fastq^ software (v2.17.1.14, Illumina). In vitro fragment libraries were prepared using the NEBNext®Ultra™ II DNA Library Prep Kit for Illumina.

### Analysis of predicted prophage activity from Illumina reads coverage

For the sequenced genomes (*B. subtilis* PS-11, PS-24, PS-25, PS-233, PS-237, PS-261), predicted prophage activity was assessed by visually analyzing coverage discrepancies between prophage sequences and the rest of the genome. This involved mapping Illumina reads to their corresponding reference genome in Geneious Prime (2024.0.2) using BBMap (v38.84) with a normal sensitivity setting, and comparing the average prophage sequence coverage with the average coverage of the rest of the genome. Enrichment of prophage regions indicates more phage DNA and lytic phages.

## Supplemental information

Supplementary Figure 1-9, Supplementary Data 1-5

## Supporting information

All Supplementary figures combined

## Acknowledgements

The authors thank to students Agnieszka Surowka, Mercedes Polajnar, Ema Florjančič and Mia Babič for help with prophage induction, enrichment and DNA extraction experiments.

AD and VF were funded by the European Union (ERC, PHAGECONTROL, 101041421). Views and opinions expressed are however those of the author(s) only and do not necessarily reflect those of the European Union or the European Research Council Executive Agency. Neither the European Union nor the granting authority can be held responsible for them. IMM, PS, ES and AD were supported by Slovenian Agency for Research and Innovation [Grant number P4-0116]. IMM, PS and ES were also supported by Slovenian Agency for Research and Innovation J4-4550. PS was additionally supported by J1-4411 and BI-US/22-24-161. ATK, MLS and AD were supported by Danish National Research Foundation (DNRF137) for the Centre for Microbial Secondary Metabolites. JK was supported by the USDA NIFA Hatch Appropriations under Project PEN04853 and Accession 7005519, and the Multistate project 4666. Sequencing of some genomes was funded by the CDC (BAA 75D301-21-C-12111). UR was supported by German Research Foundation (DFG) project 460129525.

## Author contributions

A.D., P.S., I.M.M., and R.H. conceived the project; V.A.F. performed experiments. P.S., E.S., J.K., and U.R. performed genome sequencing and assembly analysis; A.D., M.L.S., E.S., V.A.F., and P.S. contributed to methods; A.D., and P.S. produced figures; A.D., P.S., M.L.Z., and I.M.M. wrote the manuscript; all authors contributed to the final version.

## Supplementary Data legends

**Supplementary Data 1. All predicted prophage elements with metadata.** Each row contains metadata for each predicted prophage element from a particular chromosome or plasmid. The metadata includes information on clustering, phage coordinates, and bacterial hosts. This dataset was used to create other graphs and figures in this manuscript.

**Supplementary Data 2. List of prophage clusters (vOTUs) with prophage abundance, references, and function prediction.** All vOTUs (column A) and the number of sequences within each vOTU (column B) are listed. Subsequent columns are reference matches for most sequences within vOTUs (column C); reference matches for the few sequences within vOTUs (column D); matches to non-MGE chromosomal regions of *B. subtilis* 168 [NC_000964.3] (column E) with indicated coverage and identity ranges for all sequences within vOTUs; and predicted functions based on matches to functional reference phages, known MGEs, or non-MGE chromosomal regions (column F). Function classification was performed as follows: matching the majority to functional reference phages - likely active prophages; matching the minority to functional reference phages, or matching the majority but >50% coverage with non-MGE chromosomal regions - possibly active prophages; matching the majority to known MGEs - functions of particular MGEs; >30% coverage and >88% identity to non-MGE chromosomal regions of *B. subtilis* 168 - likely cryptic prophages; no reference matching and <30% match to non-MGE chromosomal regions of *B. subtilis* 168 - unknown. Outliers and singletons were not included in the aforementioned analysis. Data used to create pie charts in Figure 2B and Figure 2C are also included on the right.

**Supplementary Data 3. Similarity scores of prophage elements within and between phylogenetic clades.** Presence or absence of prophages from each vOTU (excluding singletons and outliers) within members of single or multiple phylogenetic clades was calculated. Groups of strains were scored according to the presence or absence of a given prophage considering the majority rule (e.g. if a prophage was present in the majority, the number of lysogens was counted within the analyzed group, and if a prophage was present in the minority, the number of non-lysogens was counted). Number of specific lysogens and non-lysogens was divided by the total number of strains analyzed to obtain % of lysogens or non-lysogens. Results from each vOTU were averaged to give prophage arsenal similarity score.

**Supplementary Data 4. Statistics for co-occurrence or exclusion of prophages from given vOTUs within single host chromosomes.** A binary presence and absence matrix was constructed for each prophage vOTU in combination with another vOTU. Statistically significant combinations are listed. (left) Graphical representation of the co-occurrence or exclusion of prophage elements. Each dot within rectangles represents a single genome where a given prophage element is present. Red lines represent correlations between elements. P-values were transformed to -log_10_(p) to manage the wide range of values, with higher values corresponding to greater statistical significance. (right) The same prophages were plotted according to their integration site in the host chromosome.

**Supplementary Data 5. Raw optical density measurements**. These data were used to plot growth curves in the presence and absence of mitomycin C, as well as to calculate the effect of mitomycin C on biomass reduction.

