## Supplementary material for "Ecology of prophage-like elements in *Bacillus subtilis* at global and local geographical scale": All Supplementary figures combined

**Supplementary figures 1-9**

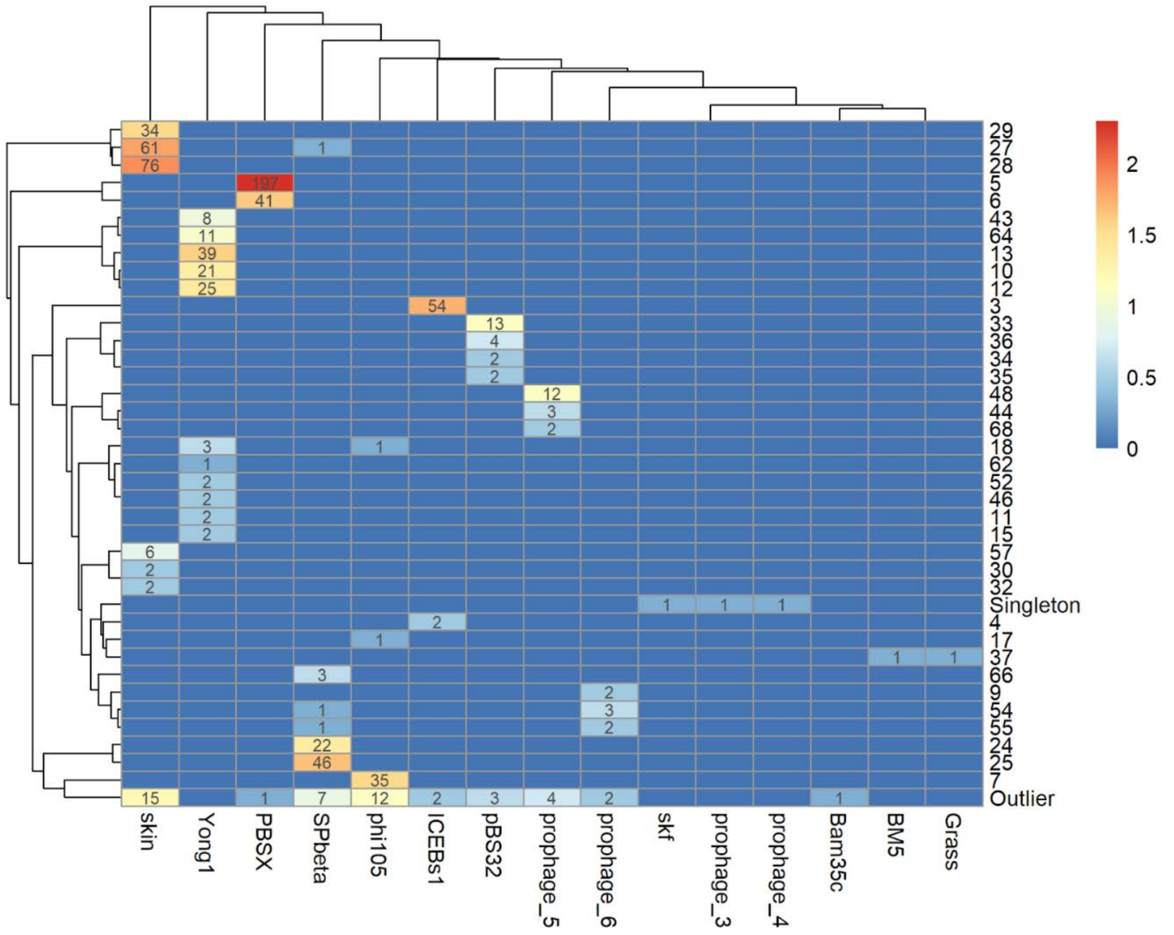


**Supplementary Figure 1. Genetic similarity of clustered prophage elements to known mobile genetic elements of *B. subtilis*.** Cluster vOTU (viral Operational Taxonomic Units) numbers are listed vertically on the right and known MGE (Mobile Genetic Elements) are listed horizontally at the bottom. Numbers in cells indicate number of prophage sequences within vOTU, which show genetic similarity with known mobile genetic element (e.g., 197 sequences within vOTU 5 shows sequence similarity to PBSX). Outliers and singletons were grouped together into artificial clusters. Outliers carry certain amount of shared proteins with different vOTU, but below the 20% threshold. Singletons have no similarity to other clusters. Heat map corresponds to numbers in cells. Dendrogram on the left corresponds to vOTU clustering, and dendrogram on top corresponds to clustering of values (number of matches) per MGE.

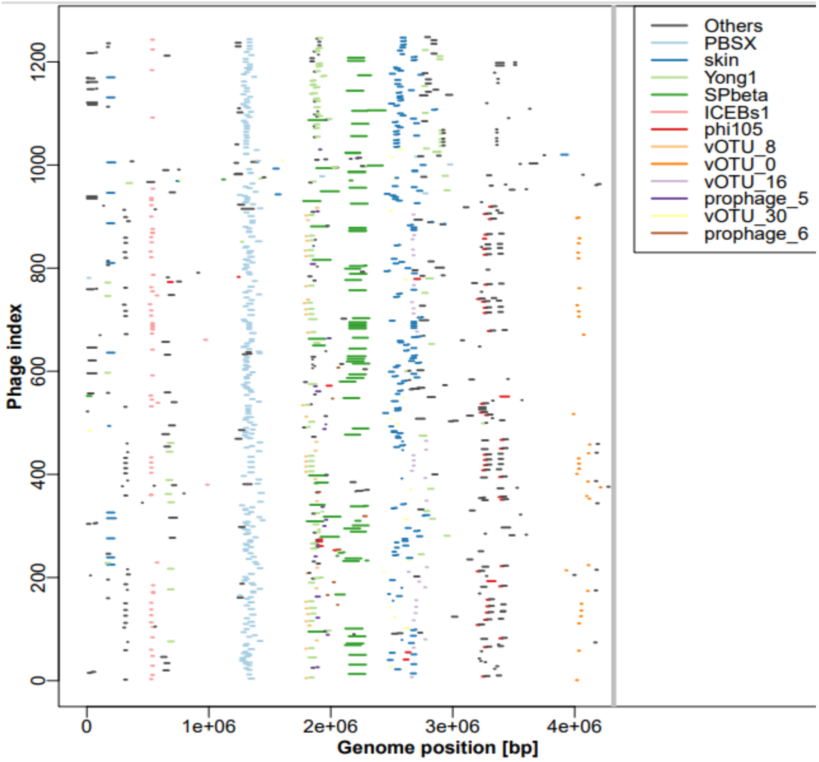


**Supplementary Figure 2: The positions of integration sites of prophage elements from selected vOTUs.** Different color represents different vOTU (viral Operational Taxonomic Units), and length of the line corresponds to prophage size.

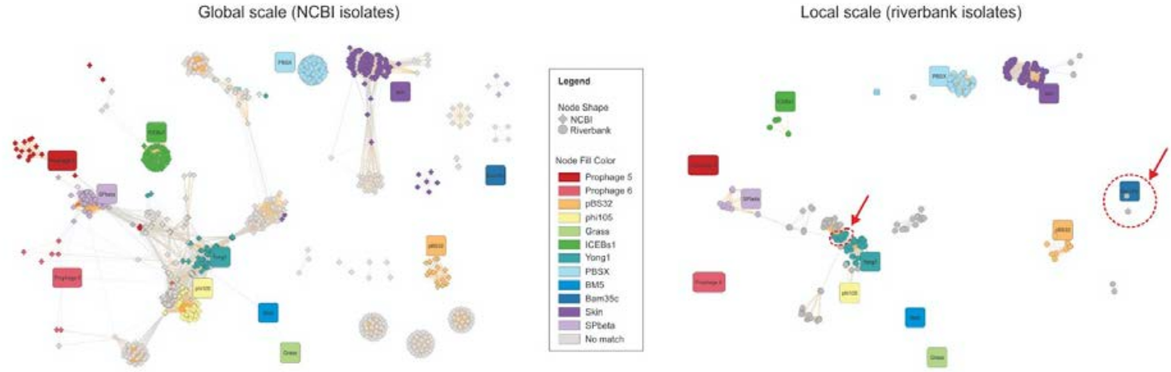


**Supplementary Figure 3. Clustering of predicted prophage elements within global and local scale isolates.** Prophages colored according to clustering. Reference phages or known mobile genetic are depicted as round squares with each color representing one phage cluster. Prophage vOTUs that are unique to local scale isolates were labeled with dashed red line.

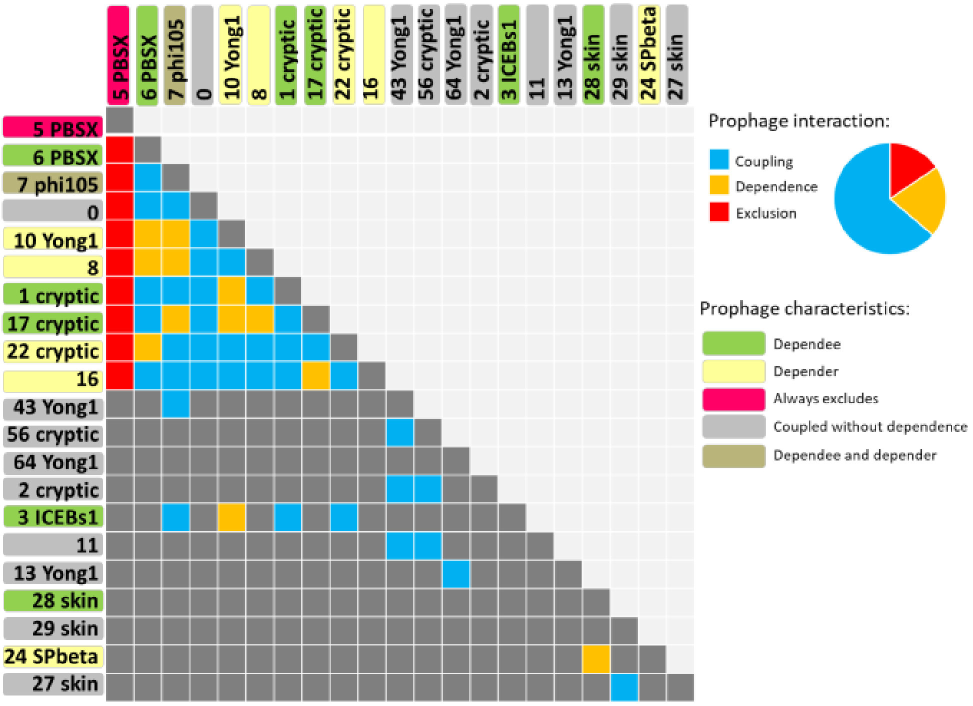


**Supplementary figure 4. Summary of prophage interactions and characteristics of prophages within viral Operational Taxonomic Units (vOTUs).** All vOTUs with significant co-occurrence or exclusion interactions were indicated. Prophage vOTUs labels were colored according to their relationships with other vOTUs within single chromosome, specifically: dependee - prophage of other vOTUs tends to occur only when this vOTU is present; depender - this vOTU occurs only in presence of another vOTU; always excludes - if this vOTU is present, certain other vOTUs never occur; coupled without dependence - tend to co-occur with another vOTU; depender and dependee - its presence depends on certain vOTU, but also determines the presence of certain vOTU. Squares in the chart are colored according to interactions, specifically: coupling, exclusion, and dependence.

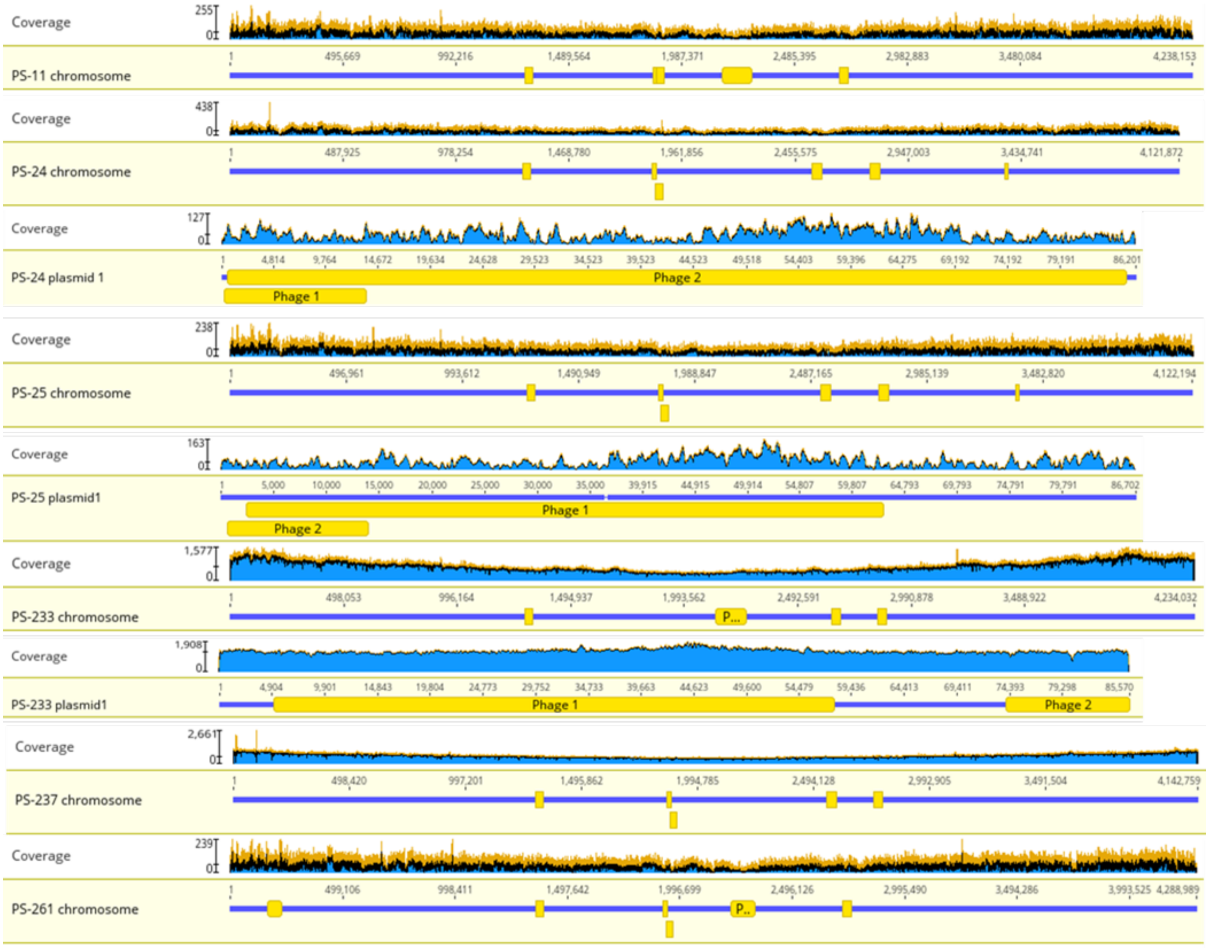


**Supplementary Figure 5**: **Coverage analysis of prophage regions in chromosomes and plasmids**. This figure illustrates the coverage profile of chromosomes and plasmids of PS-11, PS-24, PS-25, PS-233, PS-237 and PS-261. For each chromosome, the coverage plot is displayed against the chromosome length. Blue, black, and orange color represent minimum, mean, and maximum coverage, respectively. The height of the graph at each position represents the number of sequences which have a non-gap character at that position. Images were generated by Geneious Prime 2024.0.2

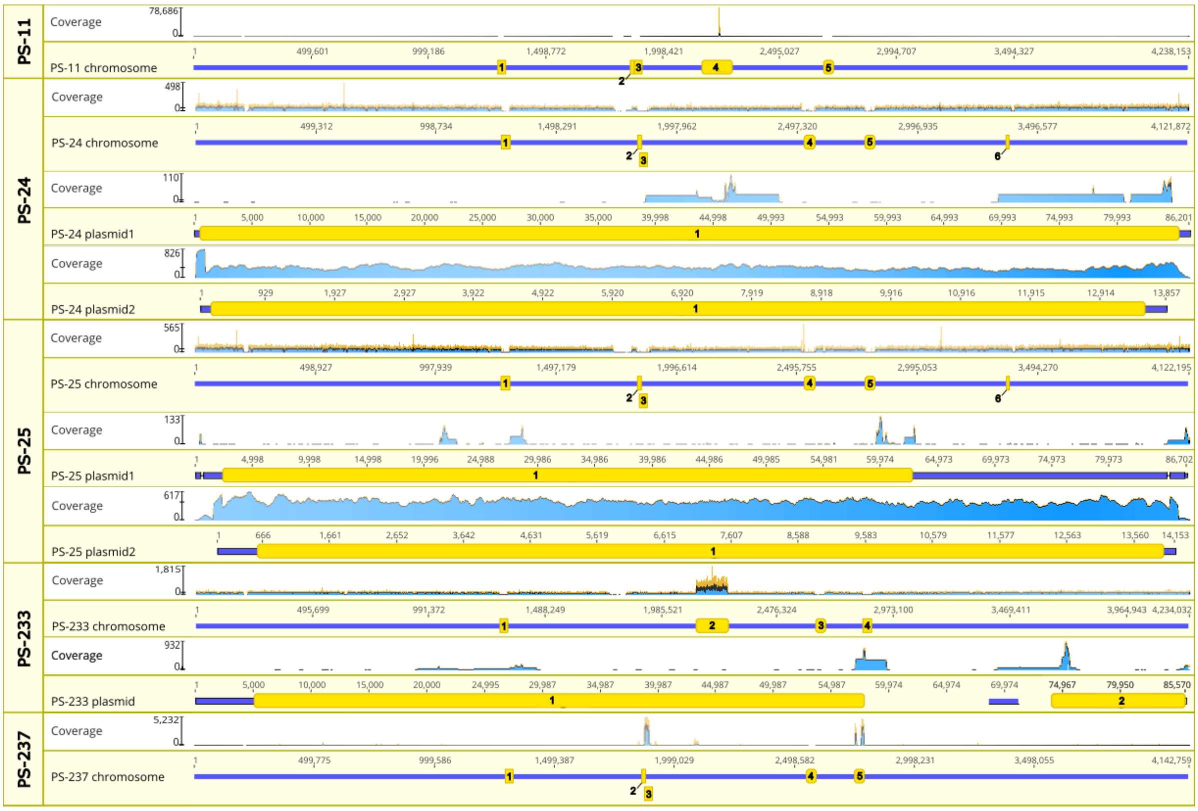


**Supplementary Figure 6**: **Coverage Analysis of prophage regions in chromosomes and plasmids**. This figure illustrates the coverage profile of chromosomes and plasmids of PS-11, PS-24, PS-25, PS-233 and PS-237. Raw sequencing reads were trimmed and filtered using fastp (0.23.4) and mapped against an assembled genome using Geneious mapper with default settings. For each chromosome and plasmid the coverage plot is displayed against the chromosome length. Blue, black and orange color represent minimum, mean and maximum coverage, respectively. The height of the graph at each position represents the number of sequences which have a non-gap character at that position. Phage locations are marked along the sequence by an orange rectangle. Mapping as well as images were generated by Geneious Prime 2024.0.5.

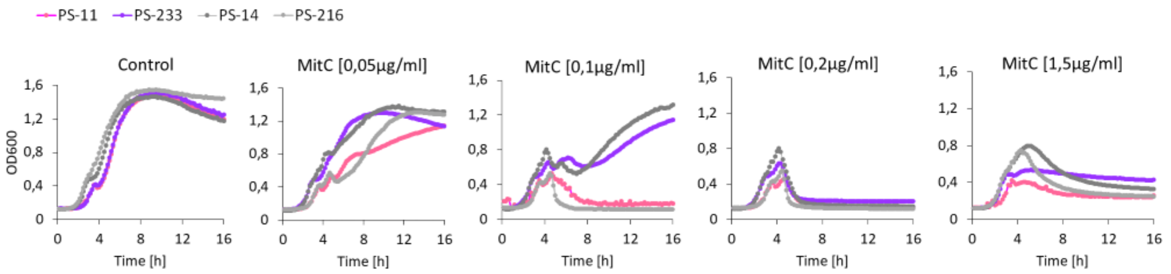


**Supplementary figure 7. Mitomycin C concentration vs growth response.** Effect of different Mitomycin C concentrations on strains that produce active phages (PS-11 and PS-233; pink and purple, respectively) vs strains that do not produce active phages (PS-14 and PS-216; light and dark grey, respectively) was assessed. Data represent average values from 6 biological replicates, error bars represent standard error.

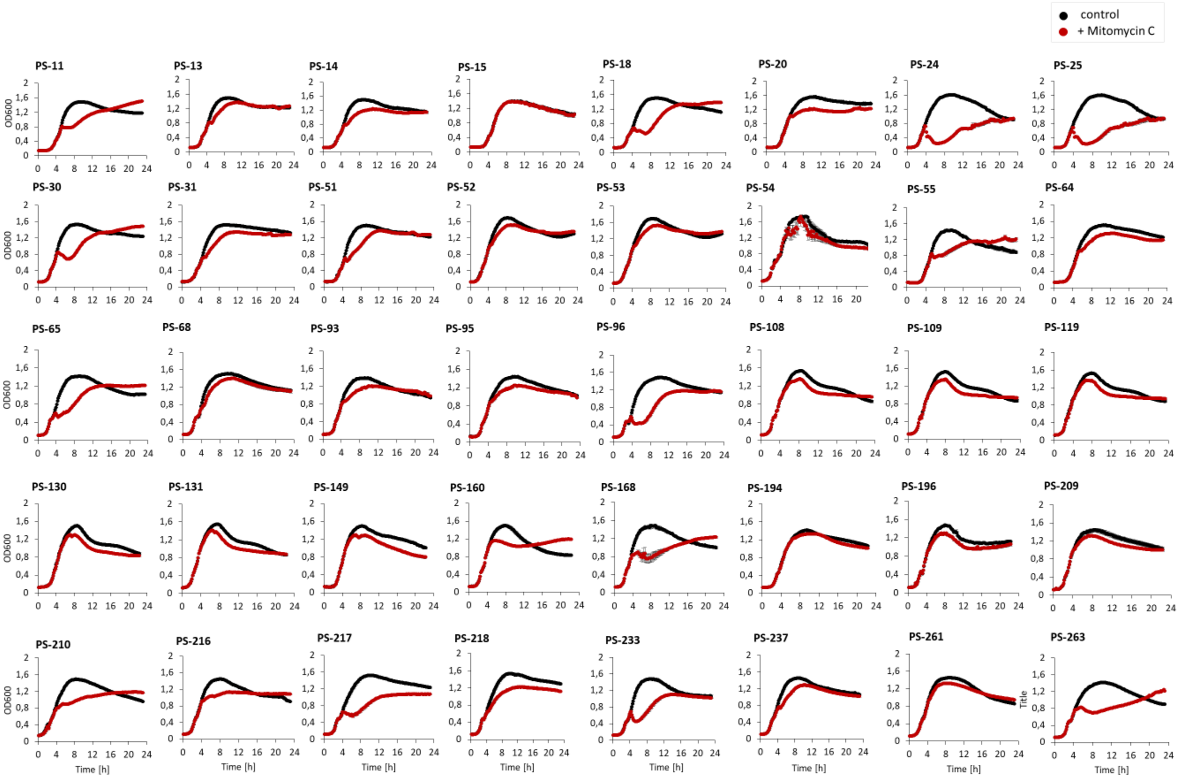


**Supplementary figure 8. Growth response to Mitomycin C treatment.** Growth curves of local scale isolates with (red) and without (black) addition of Mitomycin C (0.1ug/ml). Data represent average values from 6 biological replicates, error bars represent standard error.

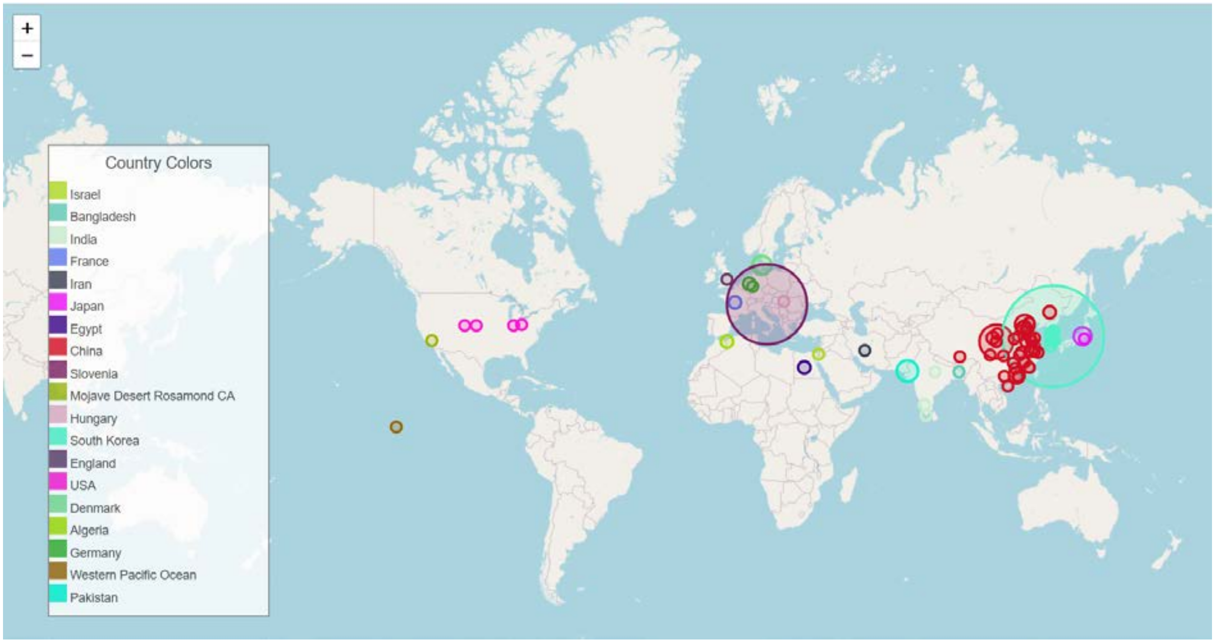


**Supplementary figure 9. Geographical distribution of isolates.** The data of strain isolation sites were extracted from genomic metadata and mapped onto the world map using Python 3.10.12 and Open cage Geocoding API. The colors and sizes of the circles correspond to the isolation source (site/country) and the number of strains isolated from the same isolation site/country, respectively.
