## Supplementary data 4 for "Ecology of prophage-like elements in *Bacillus subtilis* at global and local geographical scale"

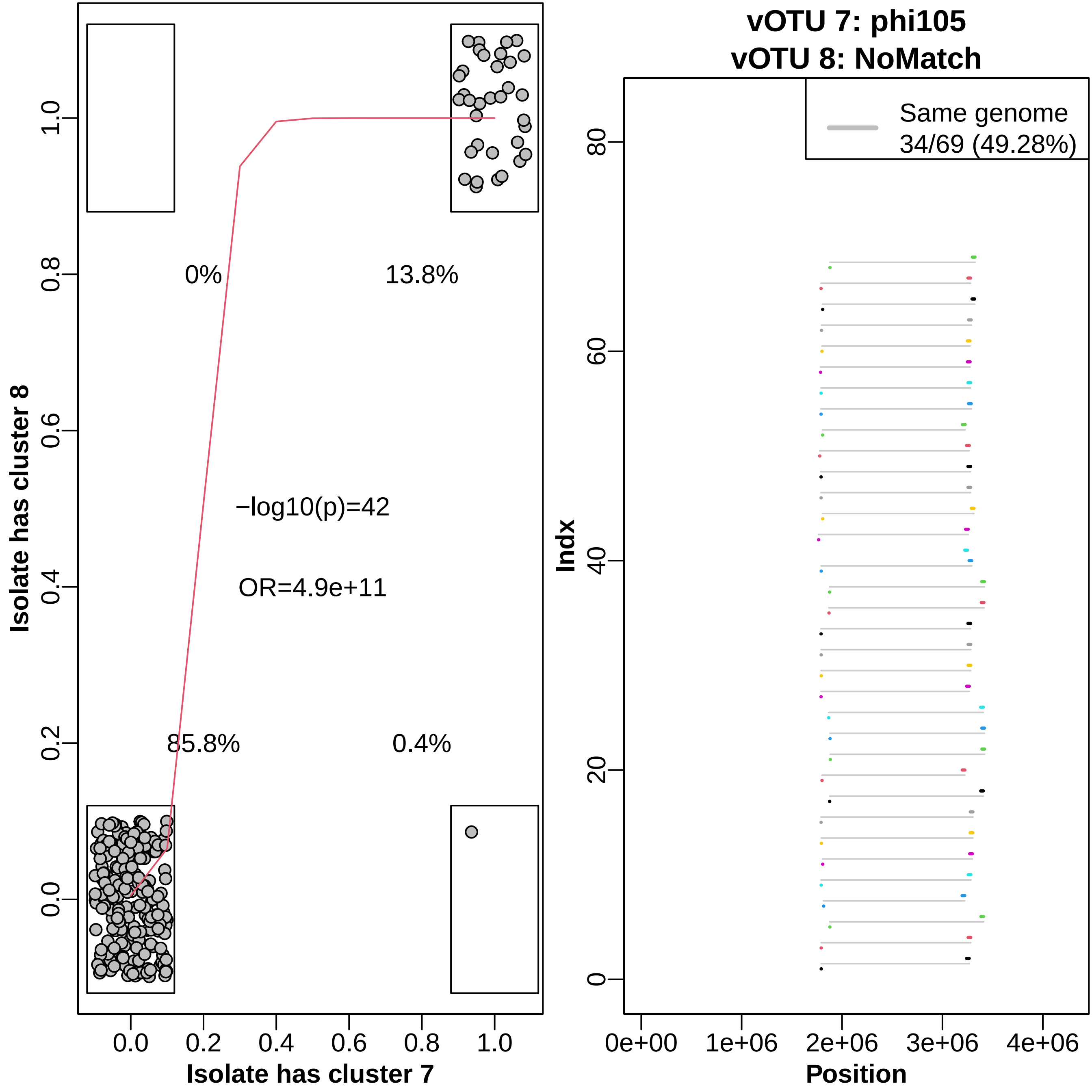

0%
13.8%
−log10(p)=42
OR=4.9e+11
85.8%
0.4%
0.0
0.2
0.4
0.6
0.8
1.0
Isolate has cluster 7
Isolate has cluster 8
vOTU 7: phi105
vOTU 8: NoMatch
Same genome
34/69 (49.28%)
Indx
0
20
40
60
80
0e+00
1e+06
2e+06
3e+06
4e+06
Position

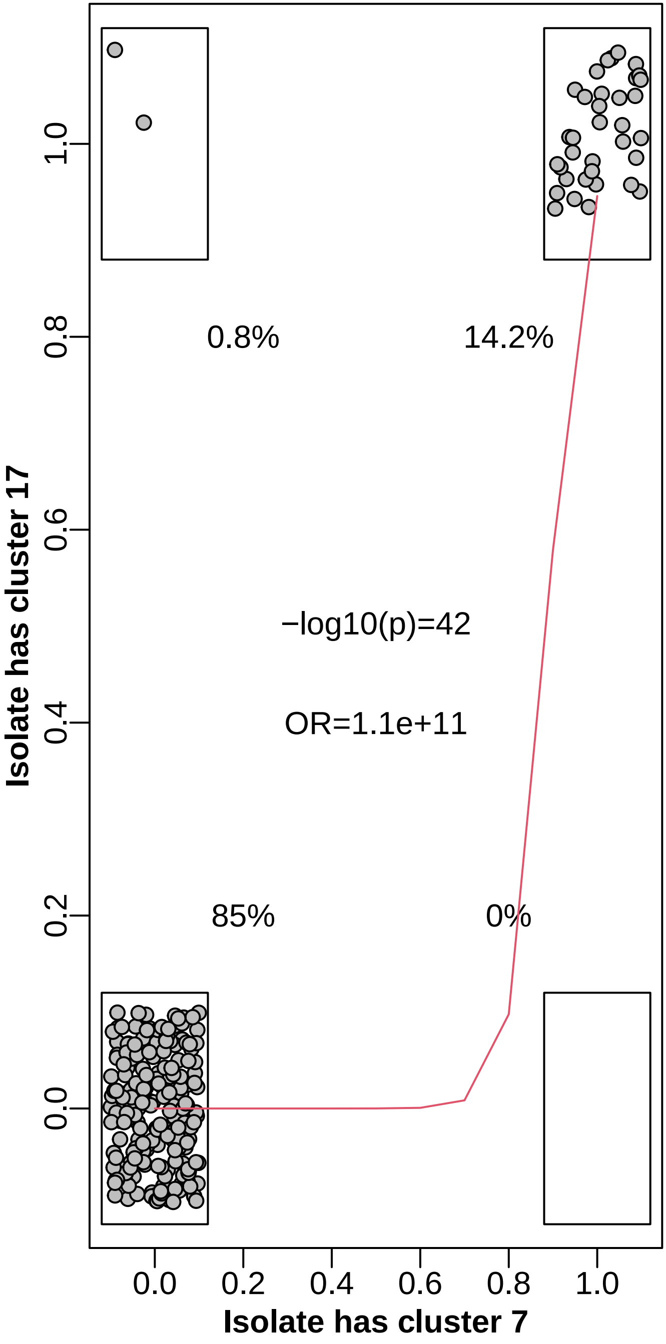

0.8%
14.2%
−log10(p)=42
OR=1.1e+11
85%
0%
Isolate has cluster 17
Isolate has cluster 7
0.0
0.2
0.4
0.6
0.8
1.0


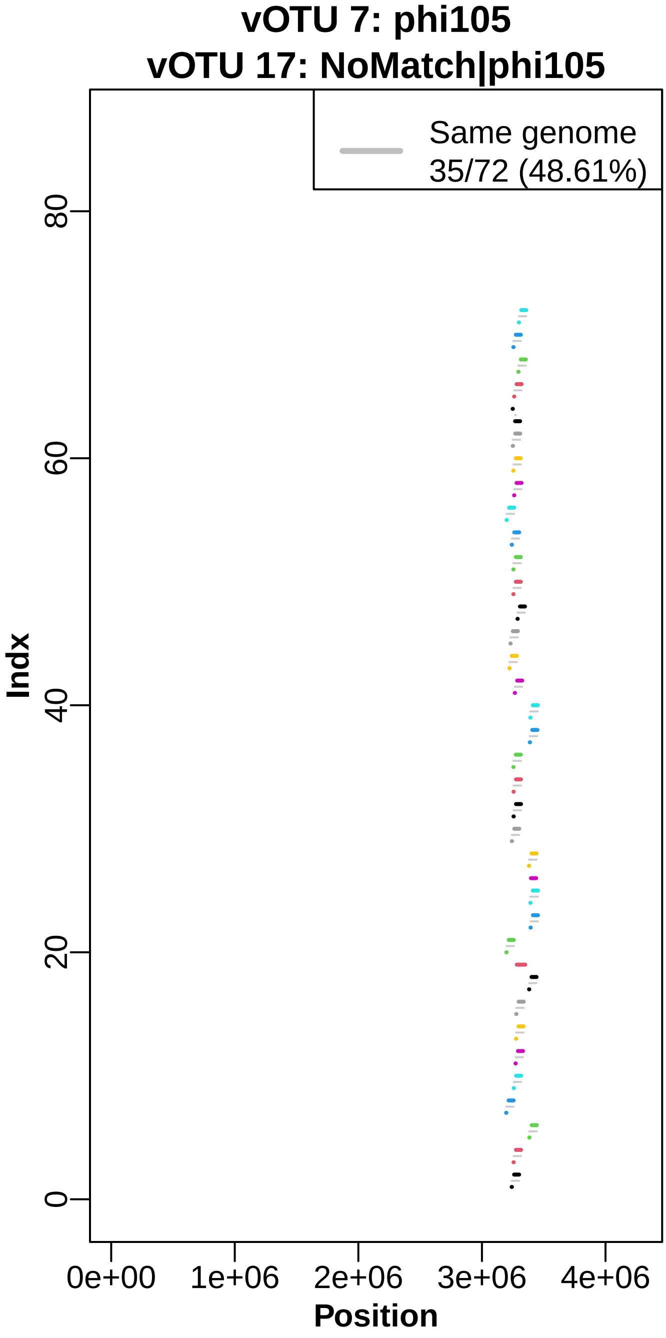

vOTU 7: phi105
vOTU 17: NoMatch|phi105
Same genome
35/72 (48.61%)
Indx
Position
0e+00
1e+06
2e+06
3e+06
4e+06
0
20
40
60
80

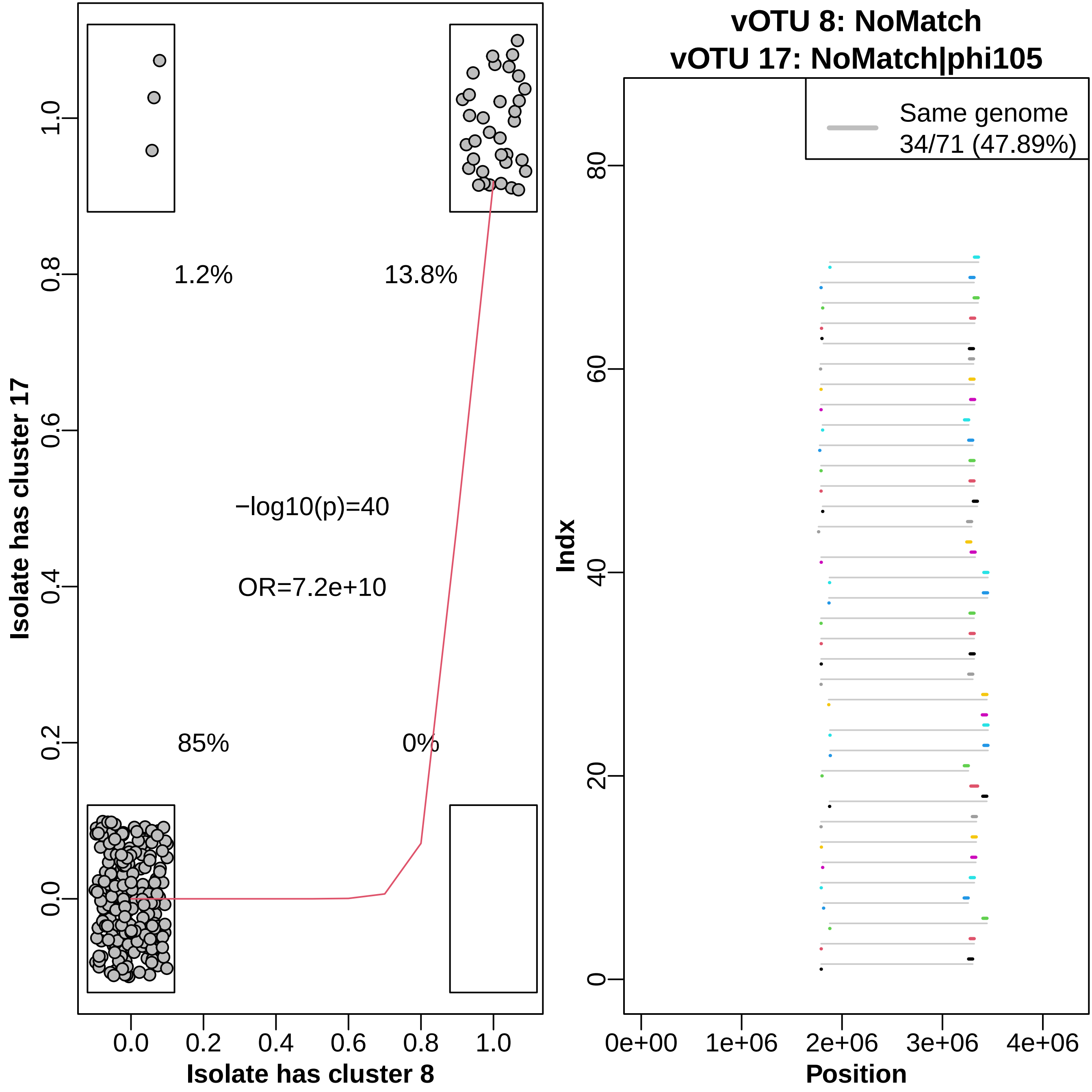

1.2%
13.8%
−log10(p)=40
OR=7.2e+10
85%
0%
Isolate has cluster 17
Isolate has cluster 8
0.0
0.2
0.4
0.6
0.8
1.0
vOTU 8: NoMatch
vOTU 17: NoMatch|phi105
Same genome
34/71 (47.89%)
Indx
Position
0
20
40
60
80
0e+00
1e+06
2e+06
3e+06
4e+06

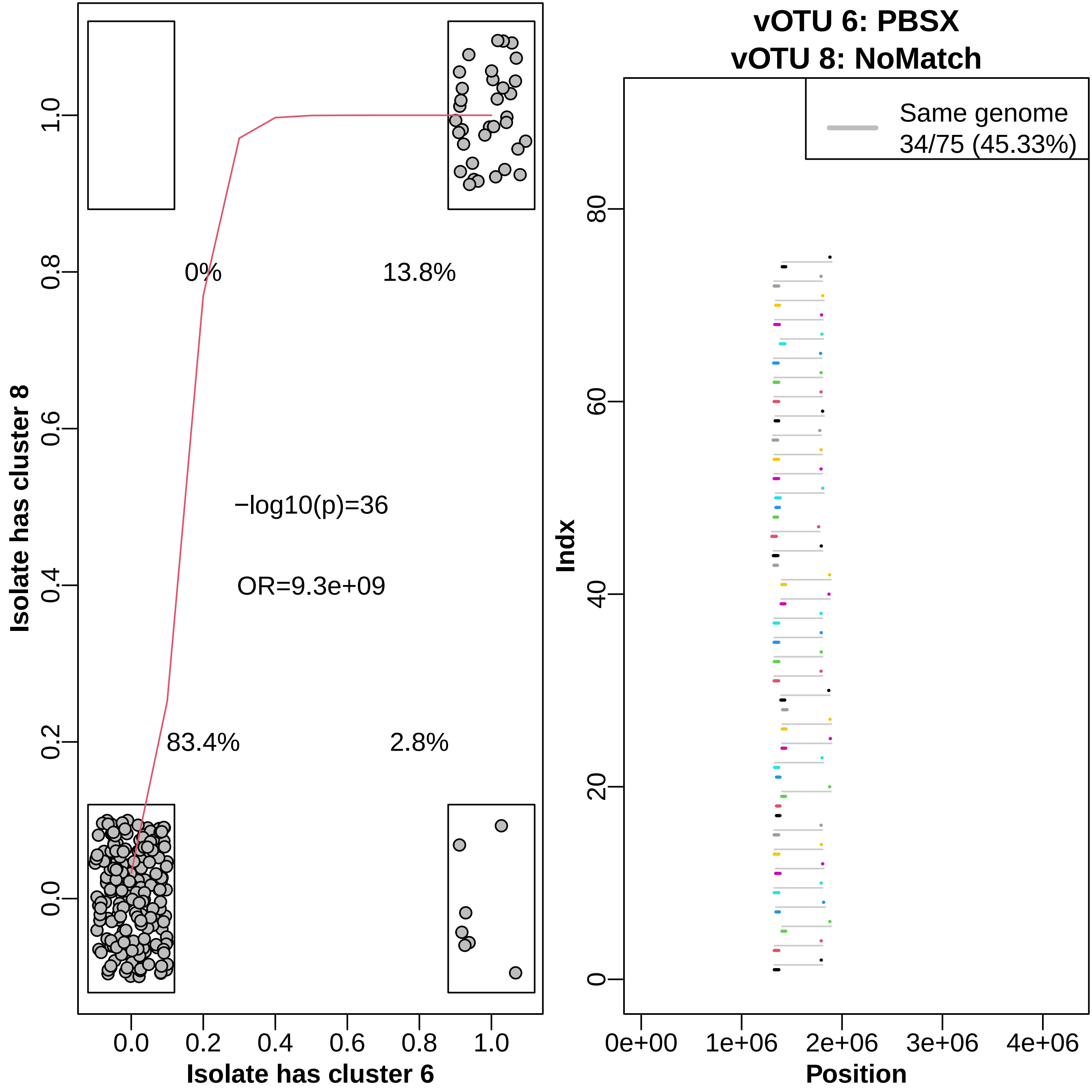

0%
13.8%
−log10(p)=36
OR=9.3e+09
83.4%
2.8%
Isolate has cluster 8
Isolate has cluster 6
0.0
0.2
0.4
0.6
0.8
1.0
vOTU 6: PBSX
vOTU 8: NoMatch
Same genome
34/75 (45.33%)
Indx
0
20
40
60
80
0e+00
1e+06
2e+06
3e+06
4e+06
Position

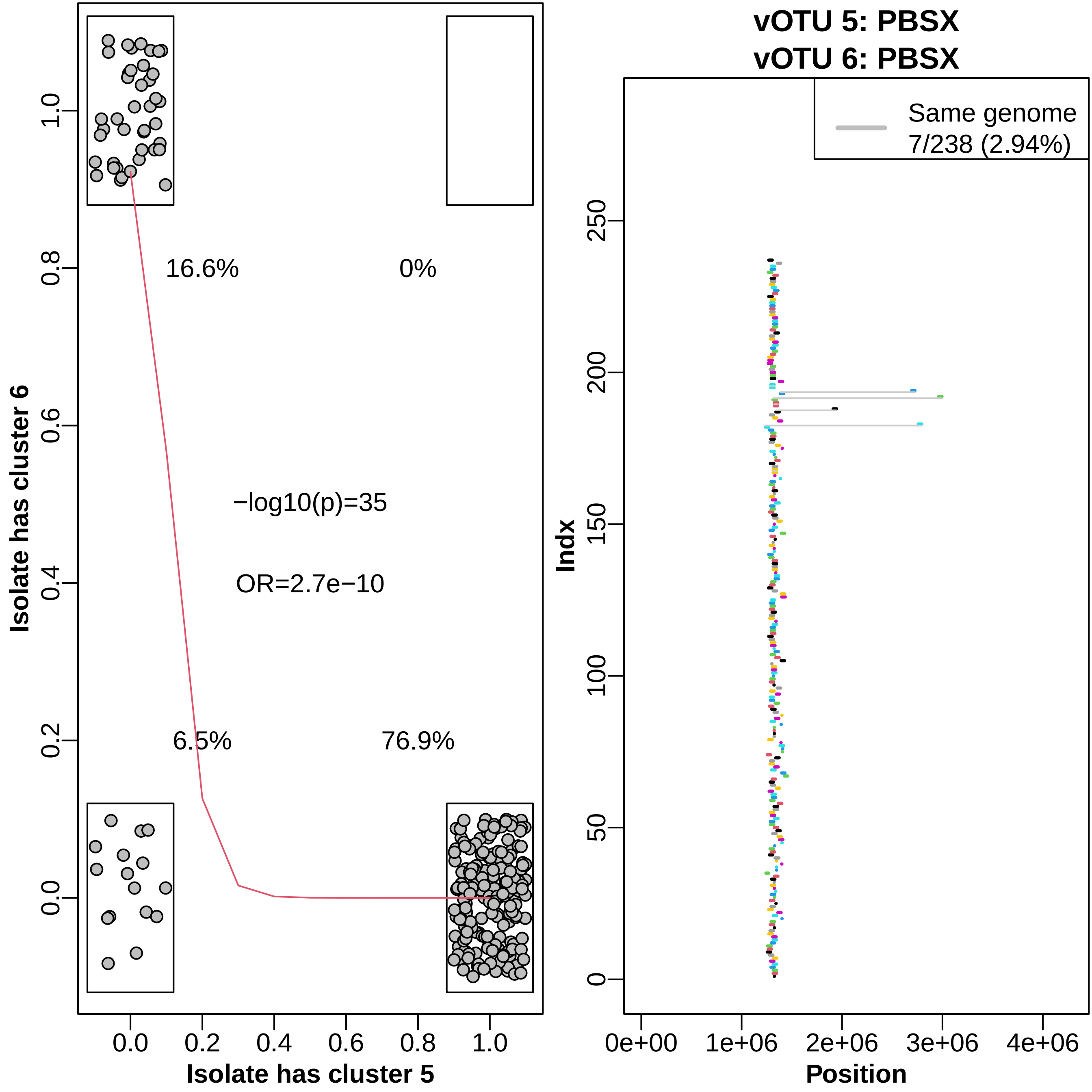

vOTU 5: PBSX
vOTU 6: PBSX
Same genome
7/238 (2.94%)
16.6%
0%
−log10(p)=35
OR=2.7e−10
6.5%
76.9%
Isolate has cluster 6
Isolate has cluster 5
Indx
Position
0.0
0.2
0.4
0.6
0.8
1.0
0
50
100
150
200
250
0e+00
1e+06
2e+06
3e+06
4e+06

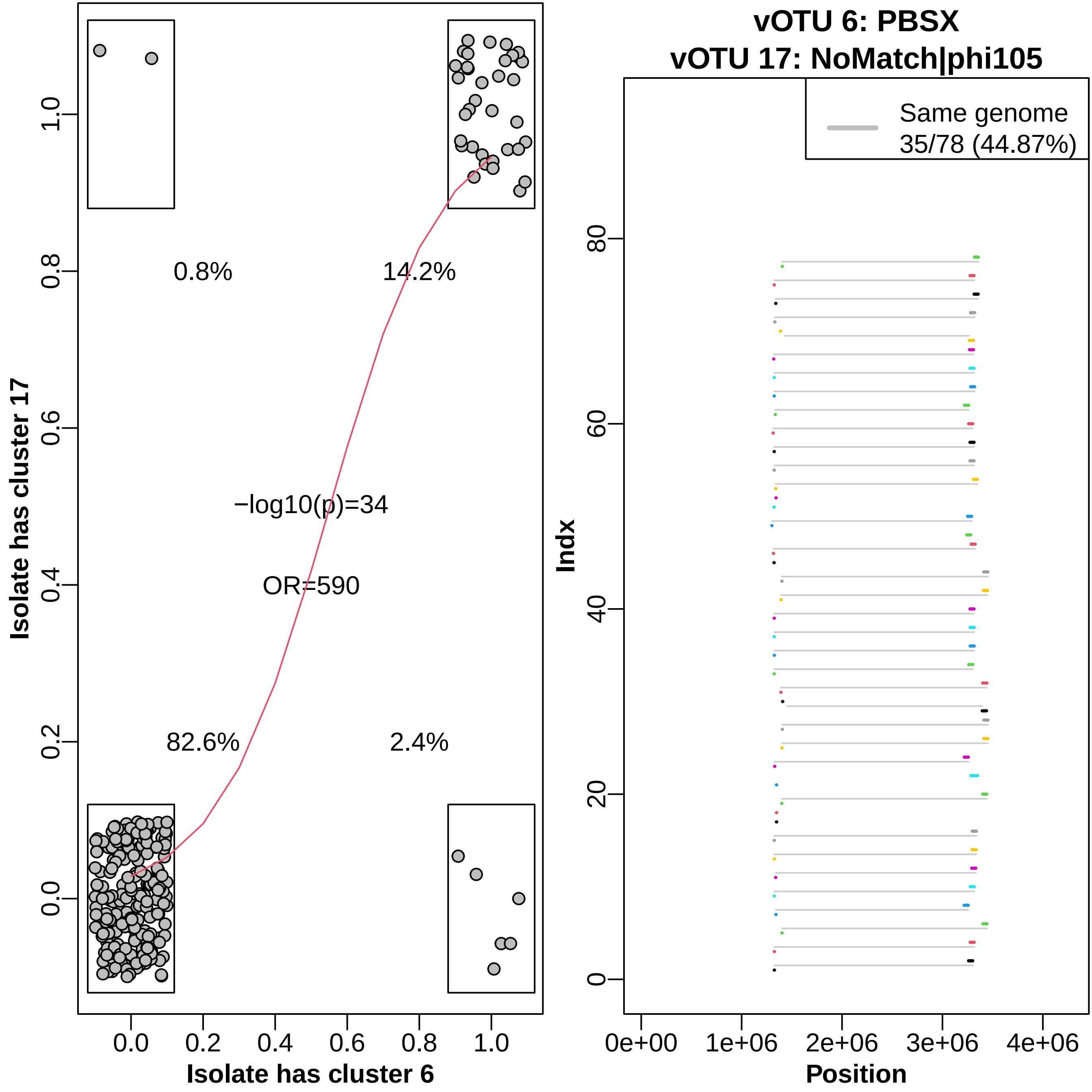

Isolate has cluster 17
Isolate has cluster 6
0.8%
14.2%
−log10(p)=34
OR=590
82.6%
2.4%
0.0
0.2
0.4
0.6
0.8
1.0
vOTU 6: PBSX
vOTU 17: NoMatch|phi105
Same genome
35/78 (44.87%)
Indx
Position
0
20
40
60
80
0e+00
1e+06
2e+06
3e+06
4e+06

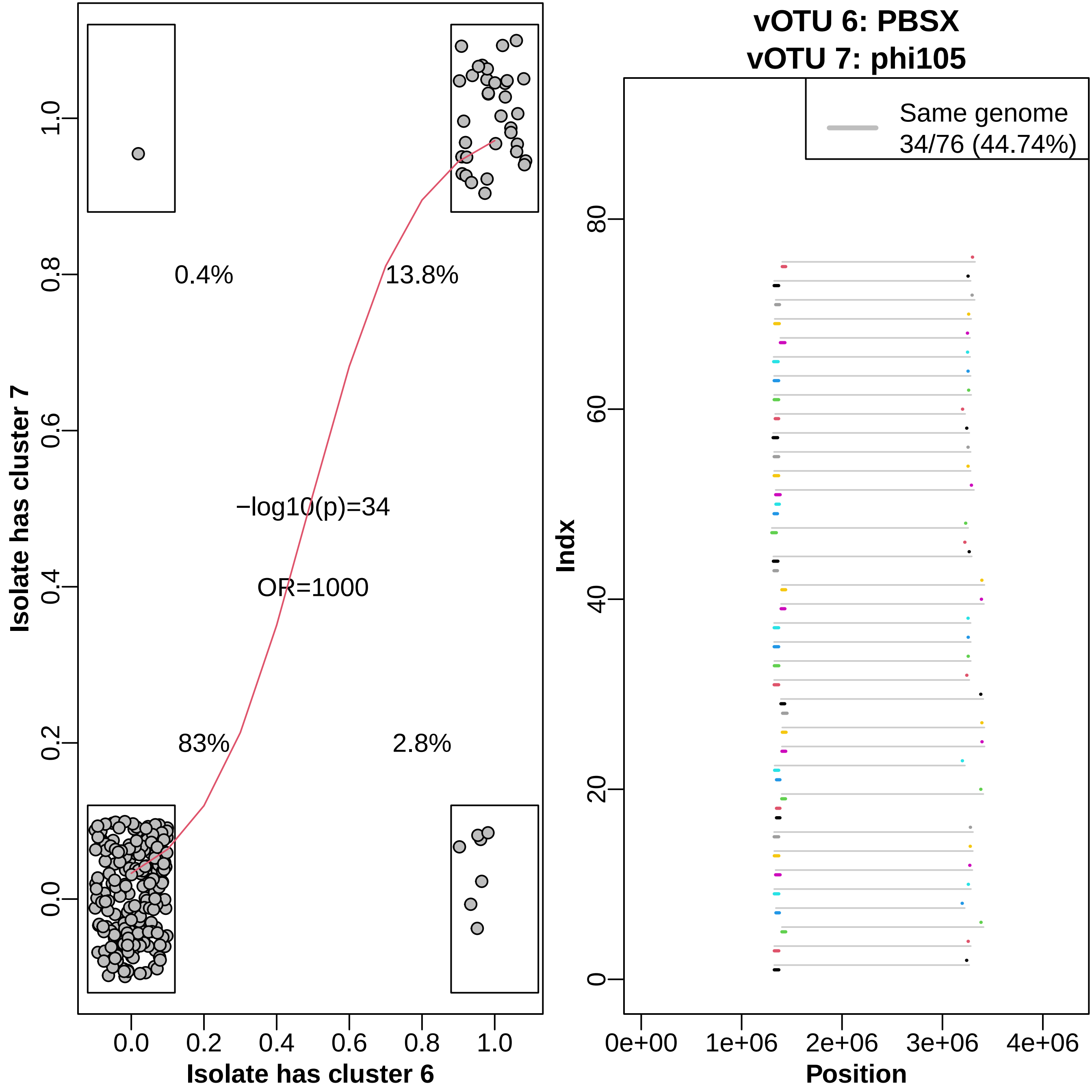

Isolate has cluster 7
Isolate has cluster 6
0.0
0.2
0.4
0.6
0.8
1.0
0.4%
13.8%
−log10(p)=34
OR=1000
83%
2.8%
vOTU 6: PBSX
vOTU 7: phi105
Same genome
34/76 (44.74%)
Indx
0
20
40
60
80
Position
0e+00
1e+06
2e+06
3e+06
4e+06

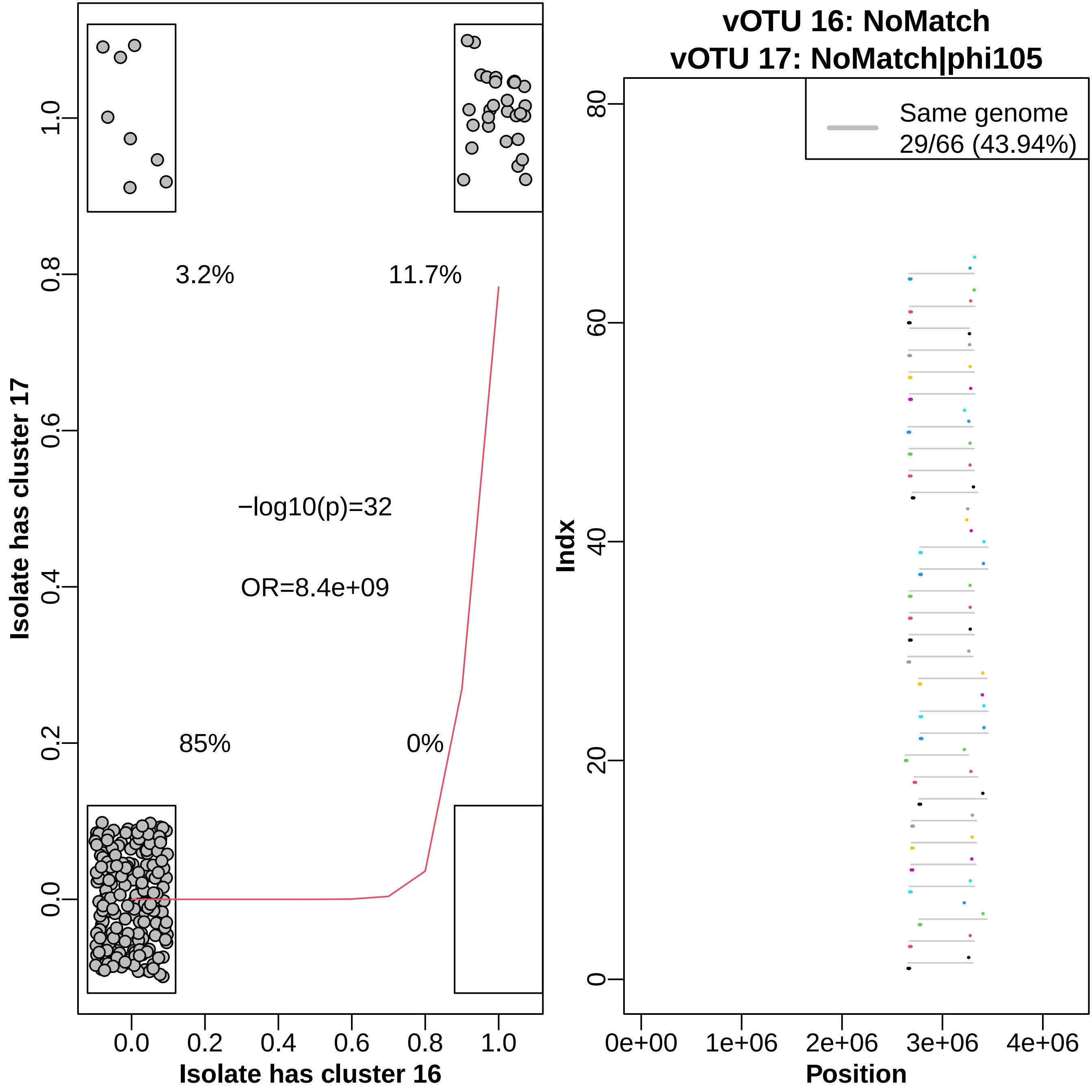

vOTU 16: NoMatch
vOTU 17: NoMatch|phi105
3.2%
11.7%
85%
0%
−log10(p)=32
OR=8.4e+09
Isolate has cluster 17
Isolate has cluster 16
0.0
0.2
0.4
0.6
0.8
1.0
Same genome
29/66 (43.94%)
Indx
Position
0e+00
1e+06
2e+06
3e+06
4e+06
0
20
40
60
80

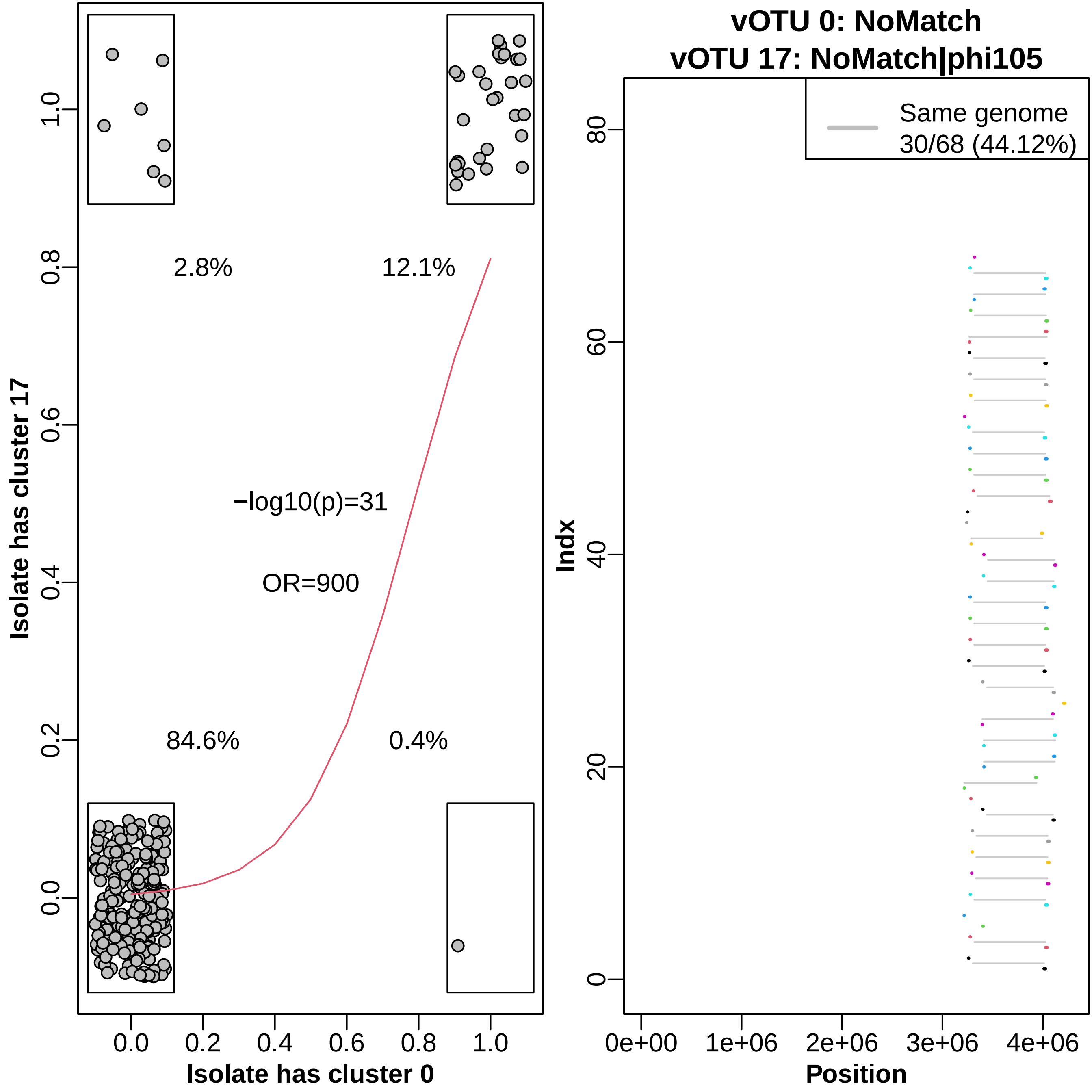

vOTU 0: NoMatch
vOTU 17: NoMatch|phi105
2.8%
12.1%
84.6%
0.4%
−log10(p)=31
OR=900
Isolate has cluster 17
Isolate has cluster 0
0.0
0.2
0.4
0.6
0.8
1.0
Same genome
30/68 (44.12%)
Indx
Position
0e+00
1e+06
2e+06
3e+06
4e+06
0
20
40
60
80

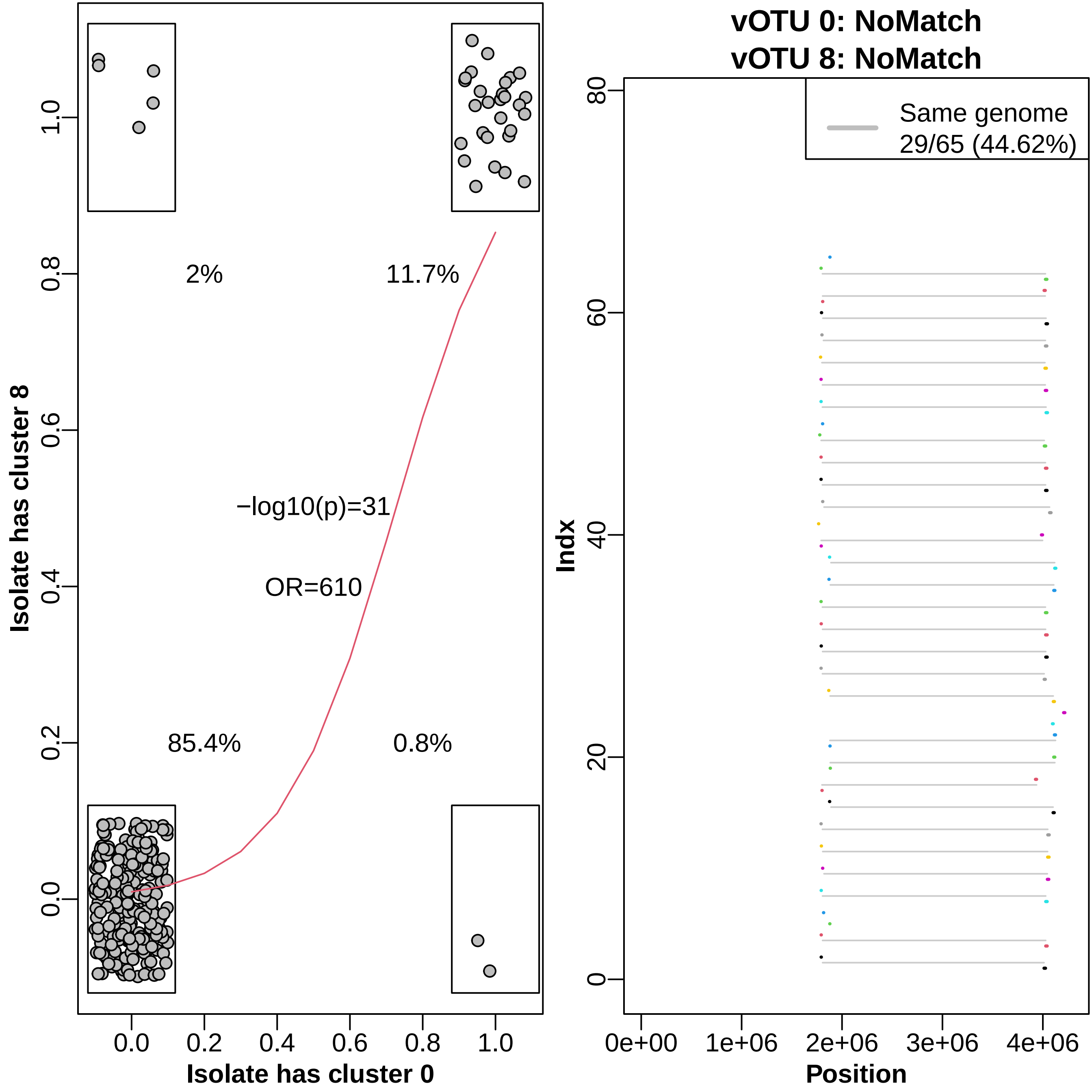

Isolate has cluster 8
Isolate has cluster 0
2%
11.7%
85.4%
0.8%
−log10(p)=31
OR=610
vOTU 0: NoMatch
vOTU 8: NoMatch
Same genome
29/65 (44.62%)
Indx
Position

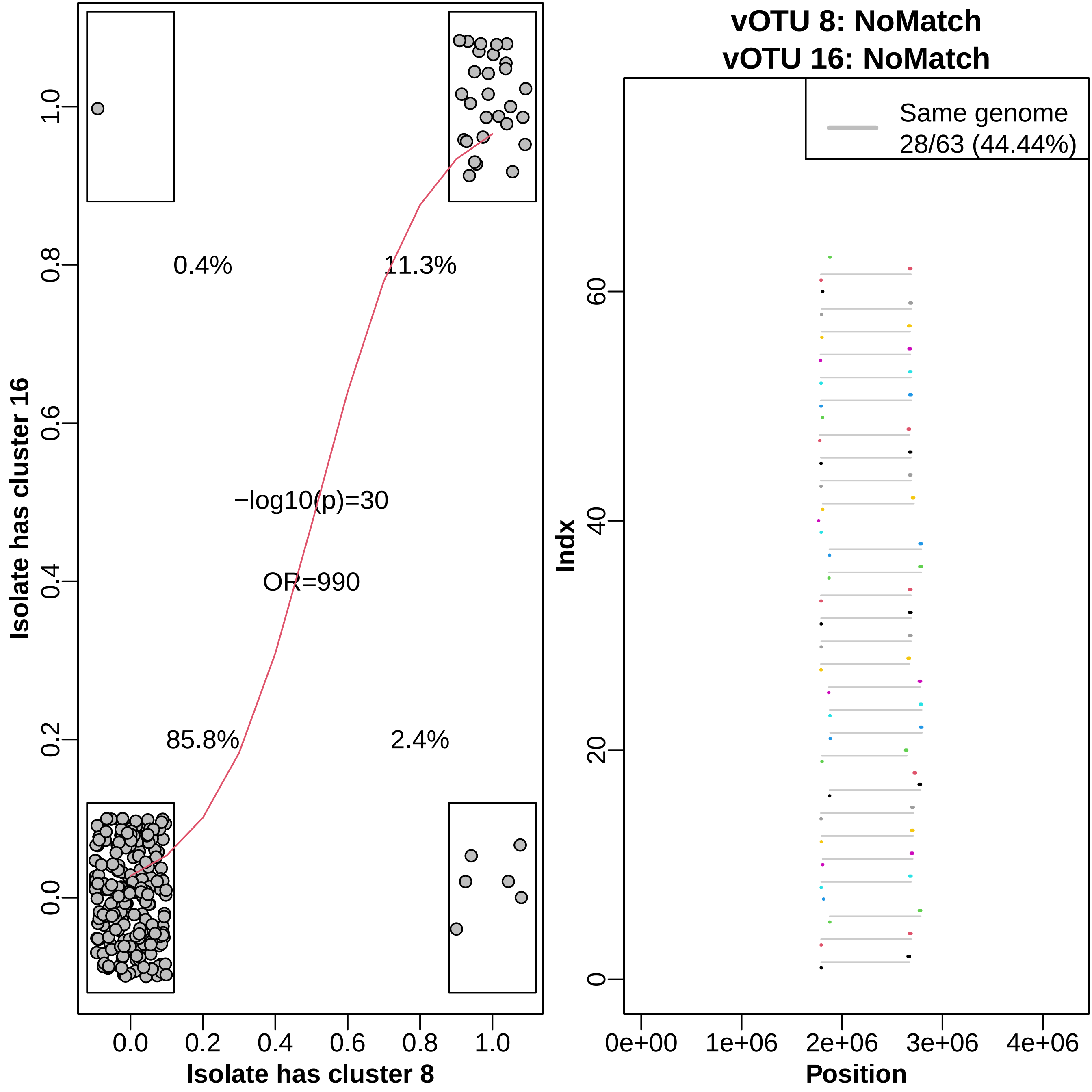

0.4%
11.3%
−log10(p)=30
OR=990
85.8%
2.4%
Isolate has cluster 16
Isolate has cluster 8
vOTU 8: NoMatch
vOTU 16: NoMatch
Same genome
28/63 (44.44%)
Indx
Position

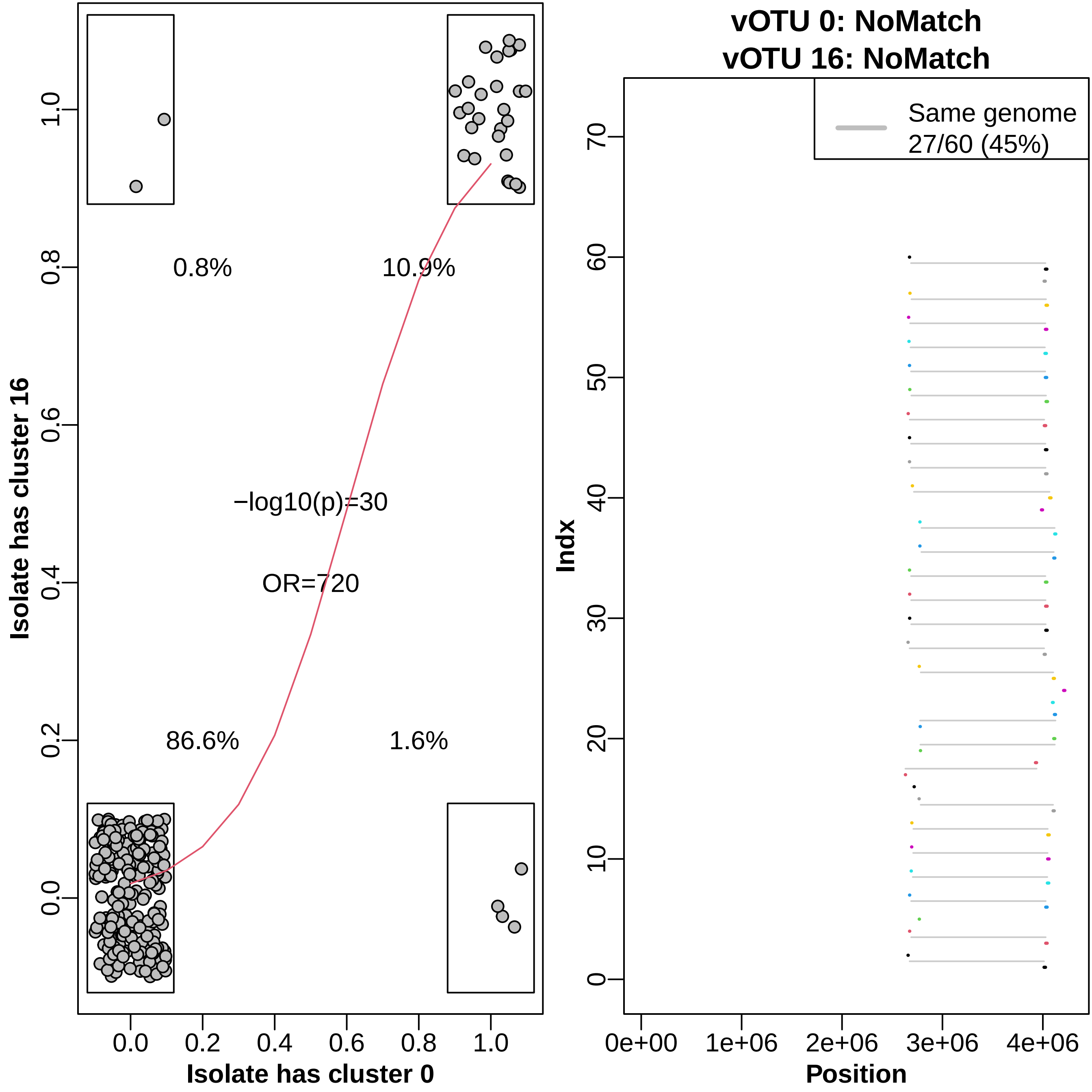

vOTU 0: NoMatch
vOTU 16: NoMatch
Isolate has cluster 16
Isolate has cluster 0
0.8%
10.9%
86.6%
1.6%
−log10(p)=30
OR=720
0.0
0.2
0.4
0.6
0.8
1.0
Same genome
27/60 (45%)
Indx
Position
0e+00
1e+06
2e+06
3e+06
4e+06
0
10
20
30
40
50
60
70

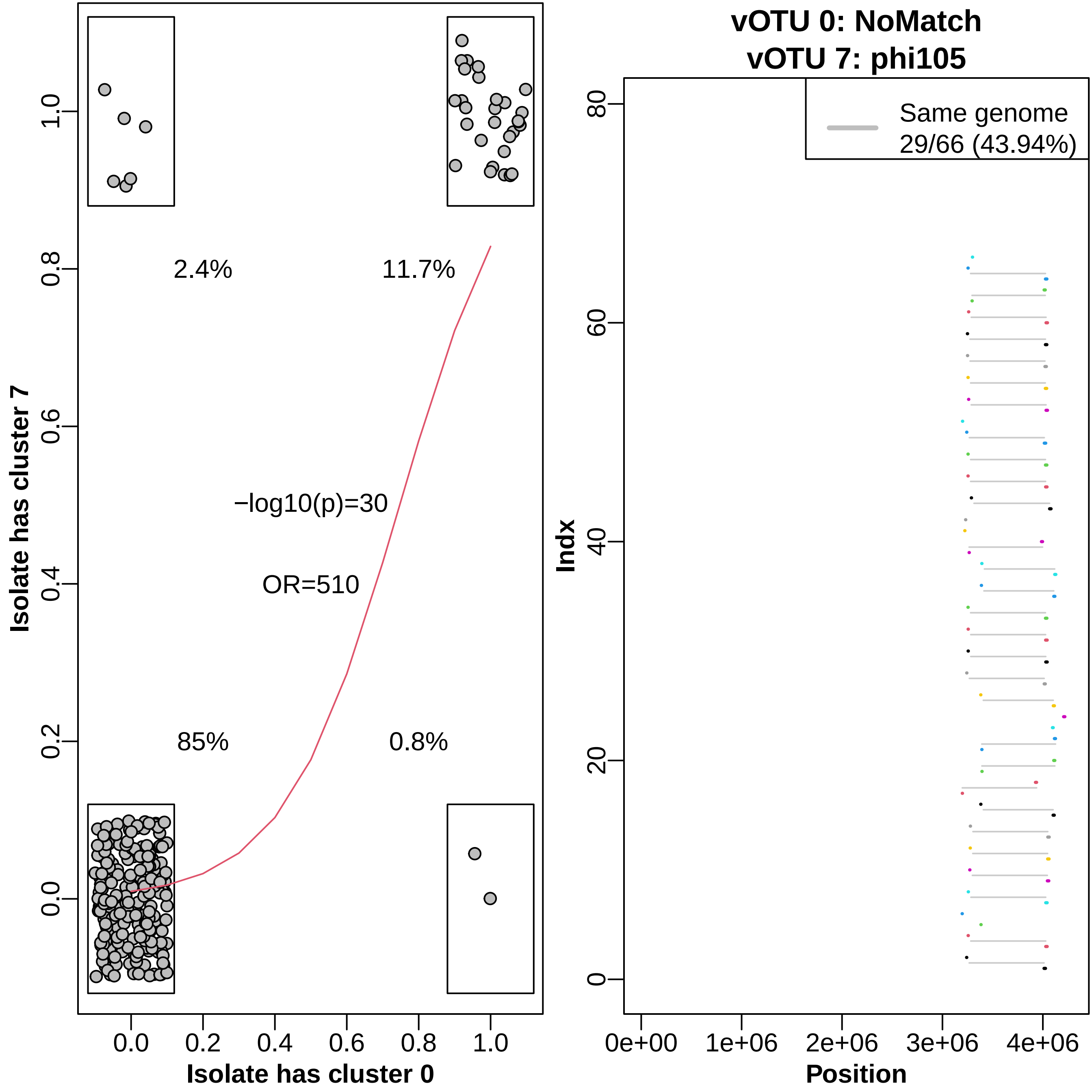

vOTU 0: NoMatch
vOTU 7: phi105
2.4%
11.7%
−log10(p)=30
OR=510
85%
0.8%
Isolate has cluster 7
Isolate has cluster 0
0.0
0.2
0.4
0.6
0.8
1.0
Same genome
29/66 (43.94%)
Indx
0
20
40
60
80
0e+00
1e+06
2e+06
3e+06
4e+06
Position

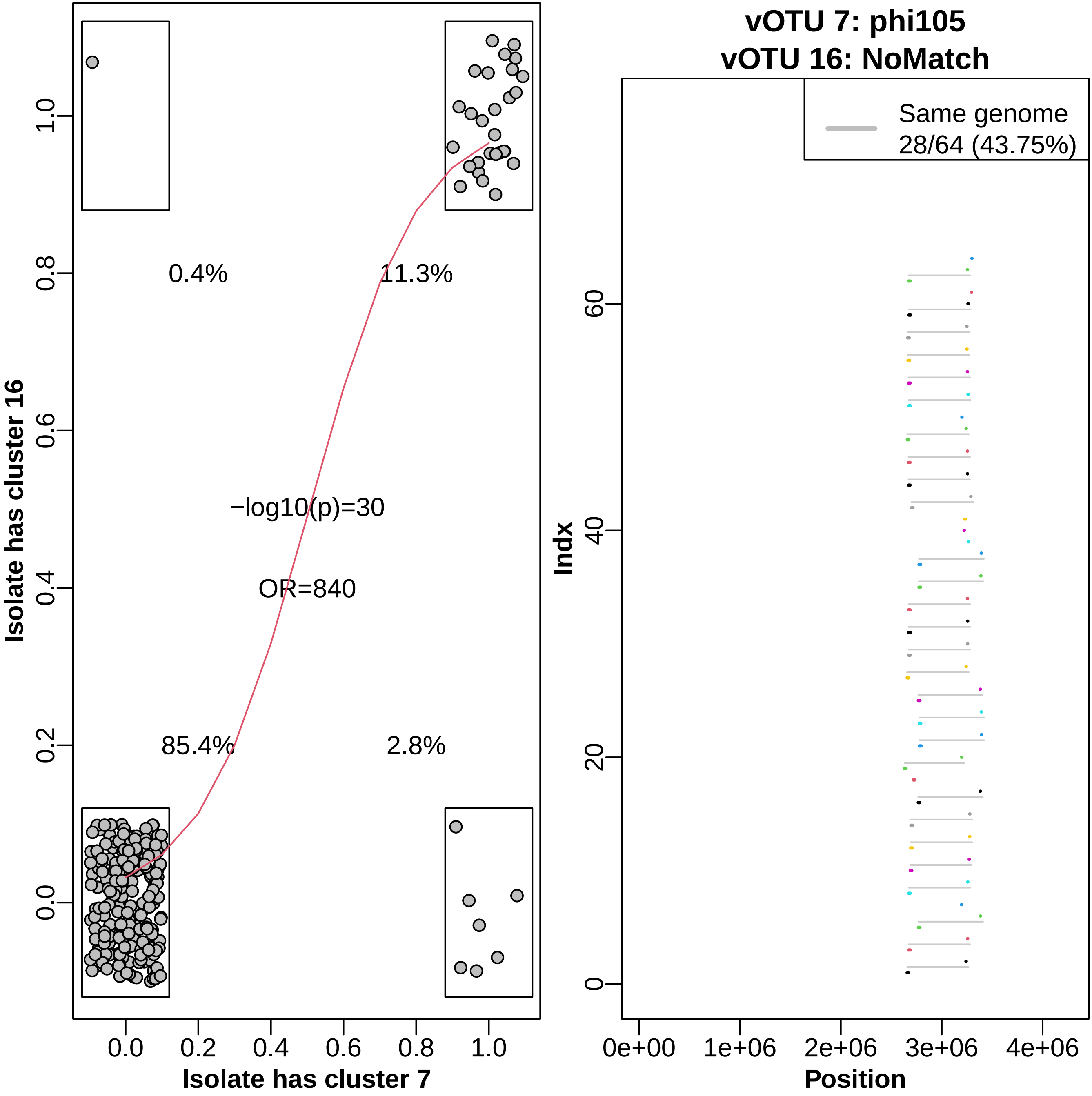

vOTU 7: phi105
vOTU 16: NoMatch
Same genome
28/64 (43.75%)
Indx
Position
0e+00
1e+06
2e+06
3e+06
4e+06
0
20
40
60
Isolate has cluster 16
Isolate has cluster 7
0.0
0.2
0.4
0.6
0.8
1.0
0.4%
11.3%
85.4%
2.8%
−log10(p)=30
OR=840

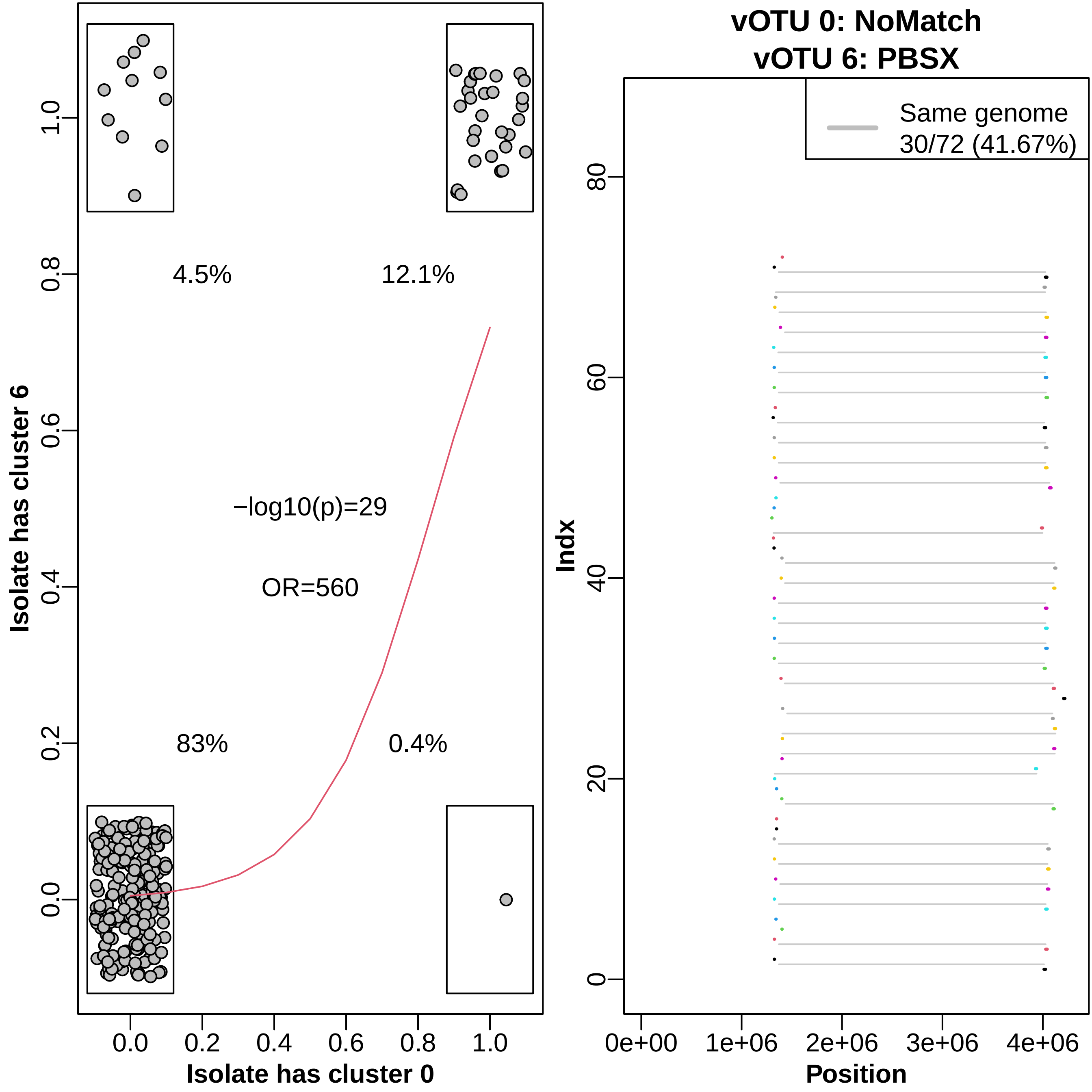

Isolate has cluster 6
Isolate has cluster 0
4.5%
12.1%
−log10(p)=29
OR=560
83%
0.4%
0.0
0.2
0.4
0.6
0.8
1.0
vOTU 0: NoMatch
vOTU 6: PBSX
Same genome
30/72 (41.67%)
Indx
Position
0
20
40
60
80
0e+00
1e+06
2e+06
3e+06
4e+06

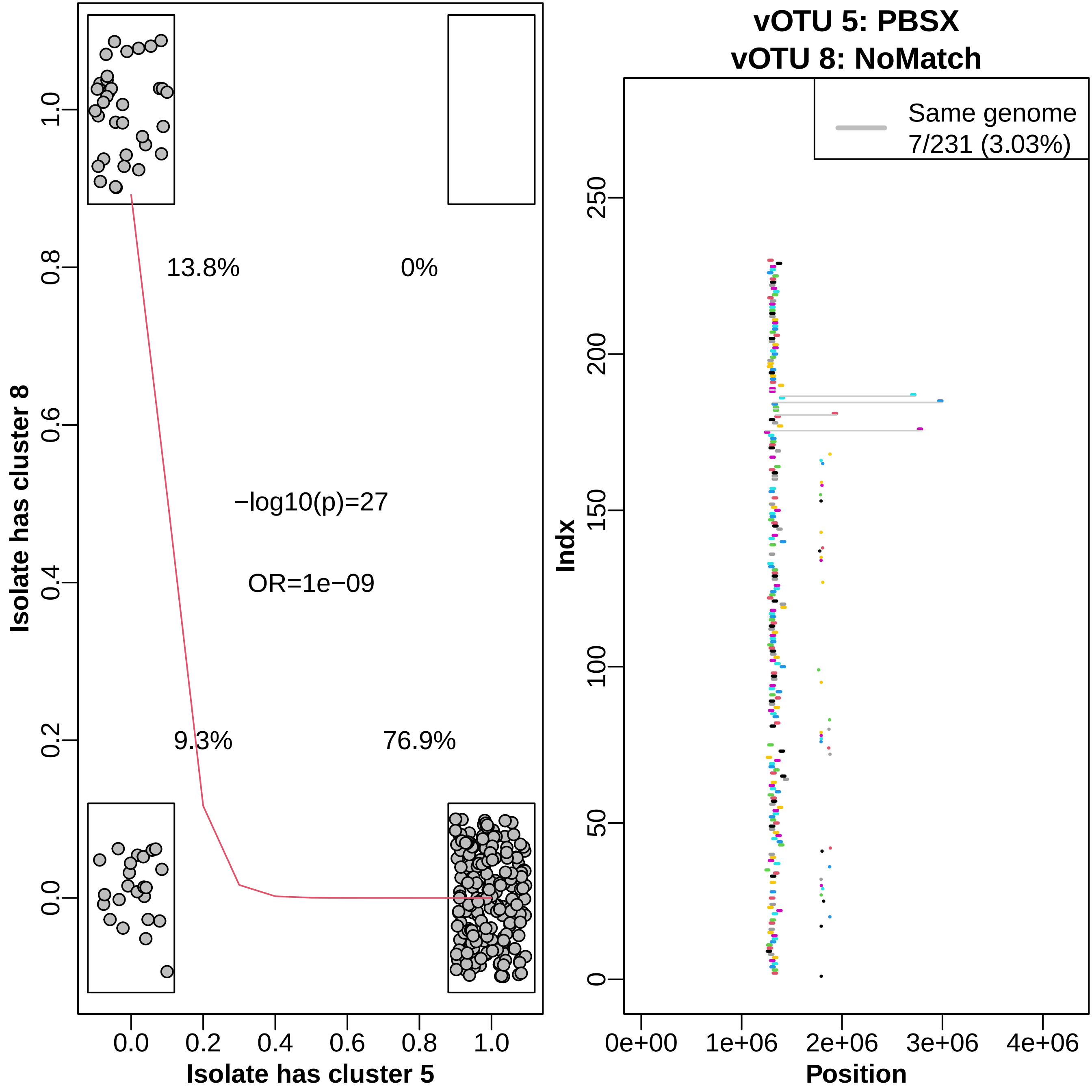

13.8%
0%
−log10(p)=27
OR=1e−09
9.3%
76.9%
Isolate has cluster 8
Isolate has cluster 5
0.0
0.2
0.4
0.6
0.8
1.0
vOTU 5: PBSX
vOTU 8: NoMatch
Same genome
7/231 (3.03%)
Indx
Position
0e+00
1e+06
2e+06
3e+06
4e+06
0
50
100
150
200
250

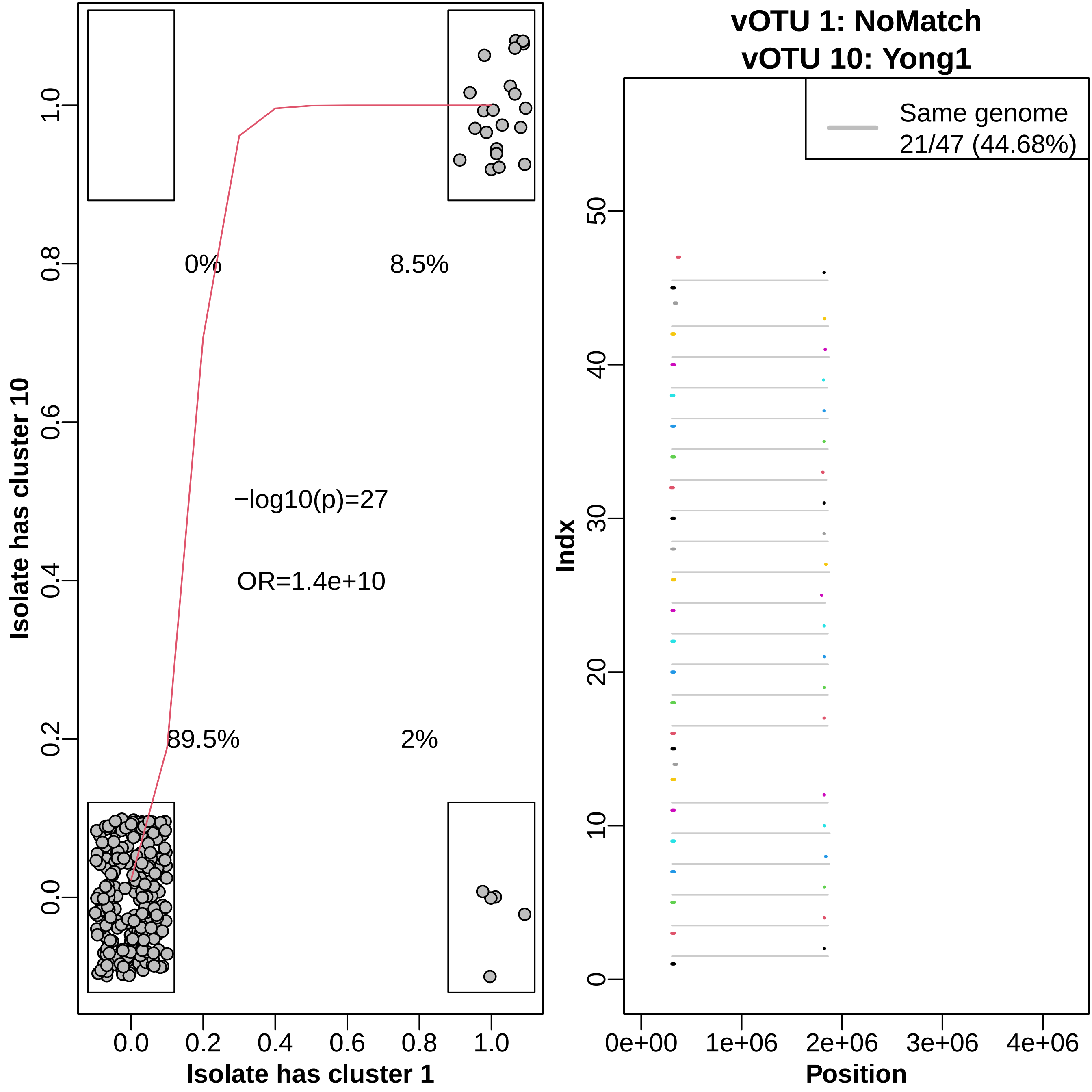

vOTU 1: NoMatch
vOTU 10: Yong1
0%
8.5%
−log10(p)=27
OR=1.4e+10
89.5%
2%
Isolate has cluster 10
Isolate has cluster 1
0.0
0.2
0.4
0.6
0.8
1.0
Same genome
21/47 (44.68%)
Indx
Position
0e+00
1e+06
2e+06
3e+06
4e+06
0
10
20
30
40
50

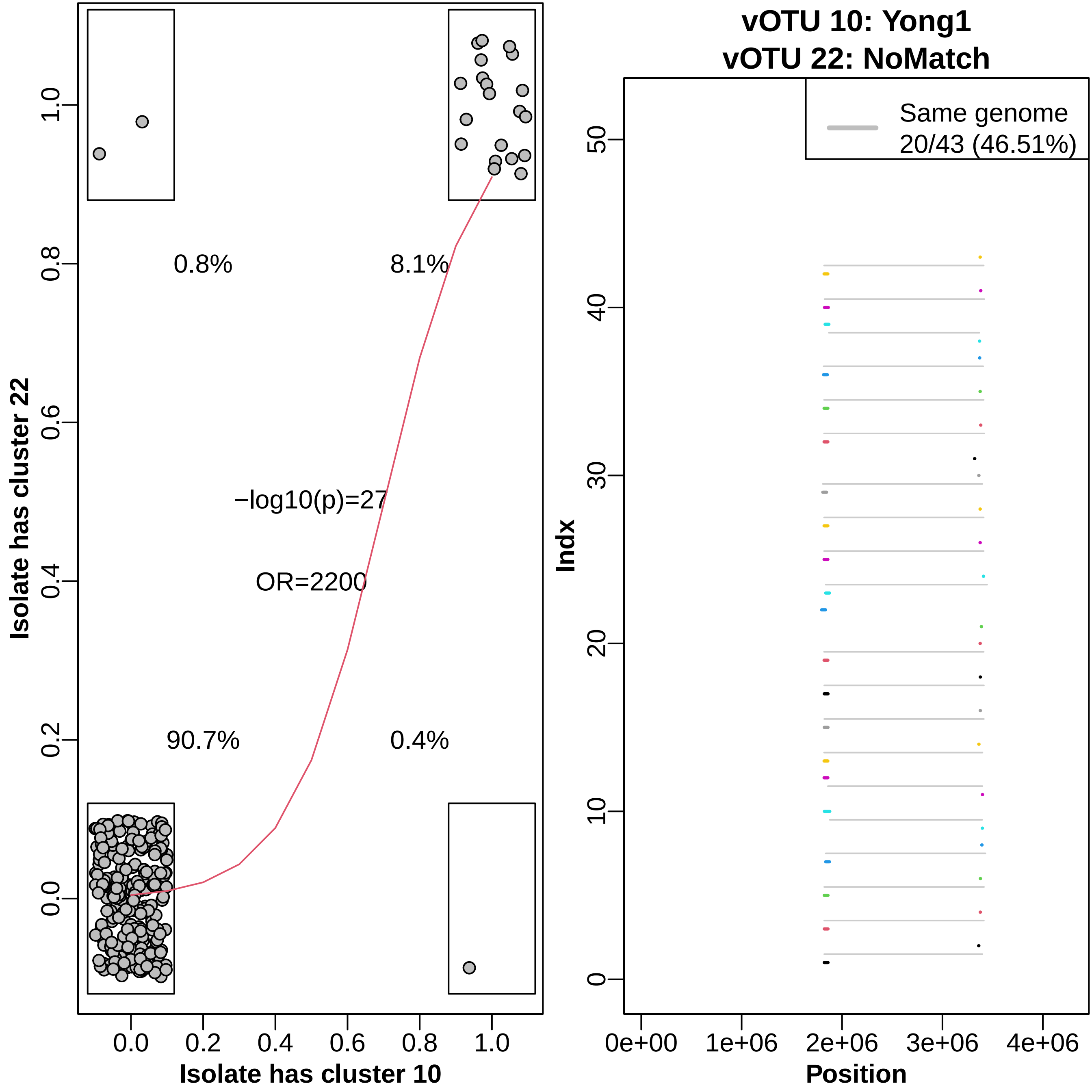

Isolate has cluster 22
Isolate has cluster 10
0.8%
8.1%
−log10(p)=27
OR=2200
90.7%
0.4%
0.0
0.2
0.4
0.6
0.8
1.0
vOTU 10: Yong1
vOTU 22: NoMatch
Same genome
20/43 (46.51%)
Indx
Position
0
10
20
30
40
50
0e+00
1e+06
2e+06
3e+06
4e+06

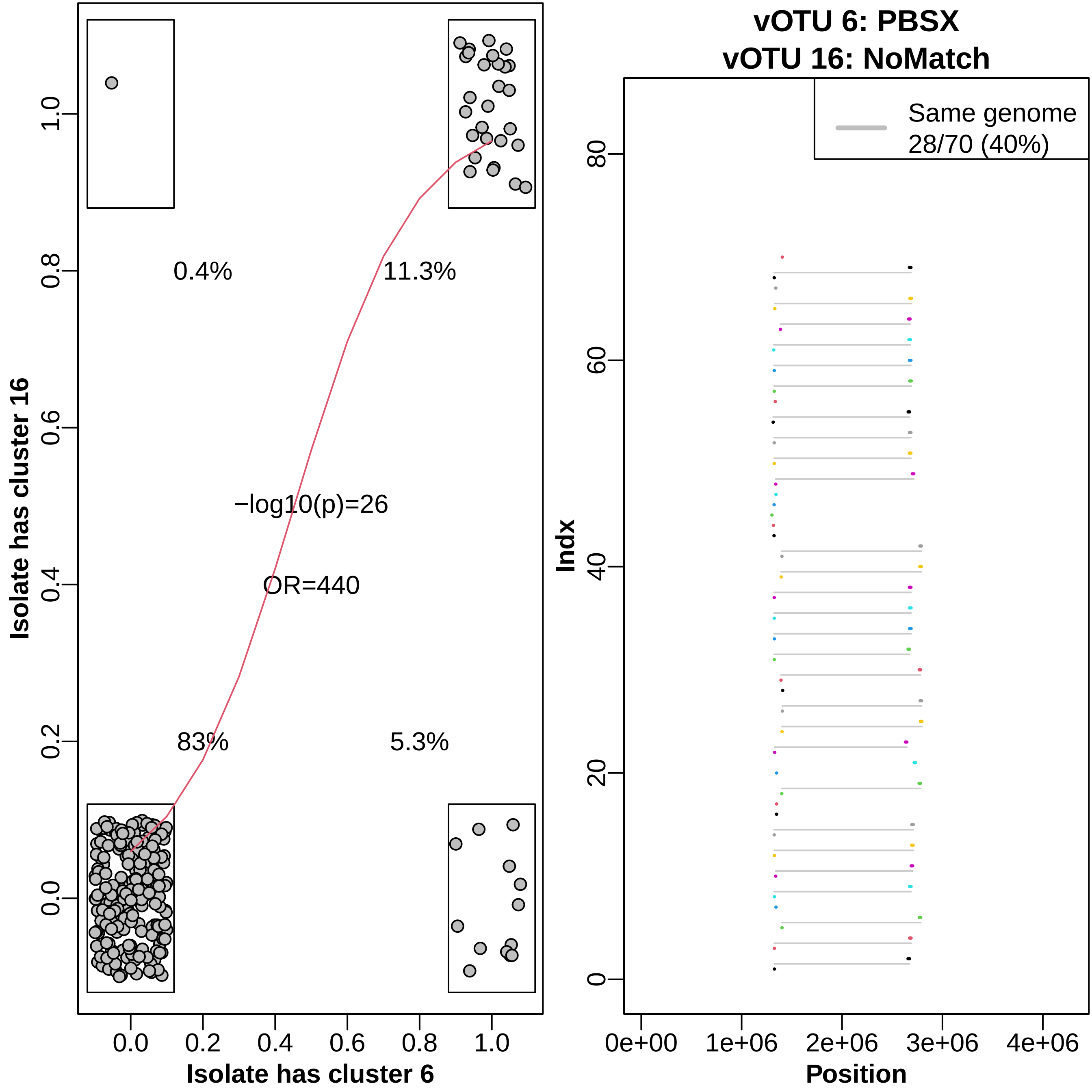

vOTU 6: PBSX
vOTU 16: NoMatch
Isolate has cluster 16
Isolate has cluster 6
0.4%
11.3%
83%
5.3%
−log10(p)=26
OR=440
0.0
0.2
0.4
0.6
0.8
1.0
Same genome
28/70 (40%)
Indx
Position
0
20
40
60
80
0e+00
1e+06
2e+06
3e+06
4e+06

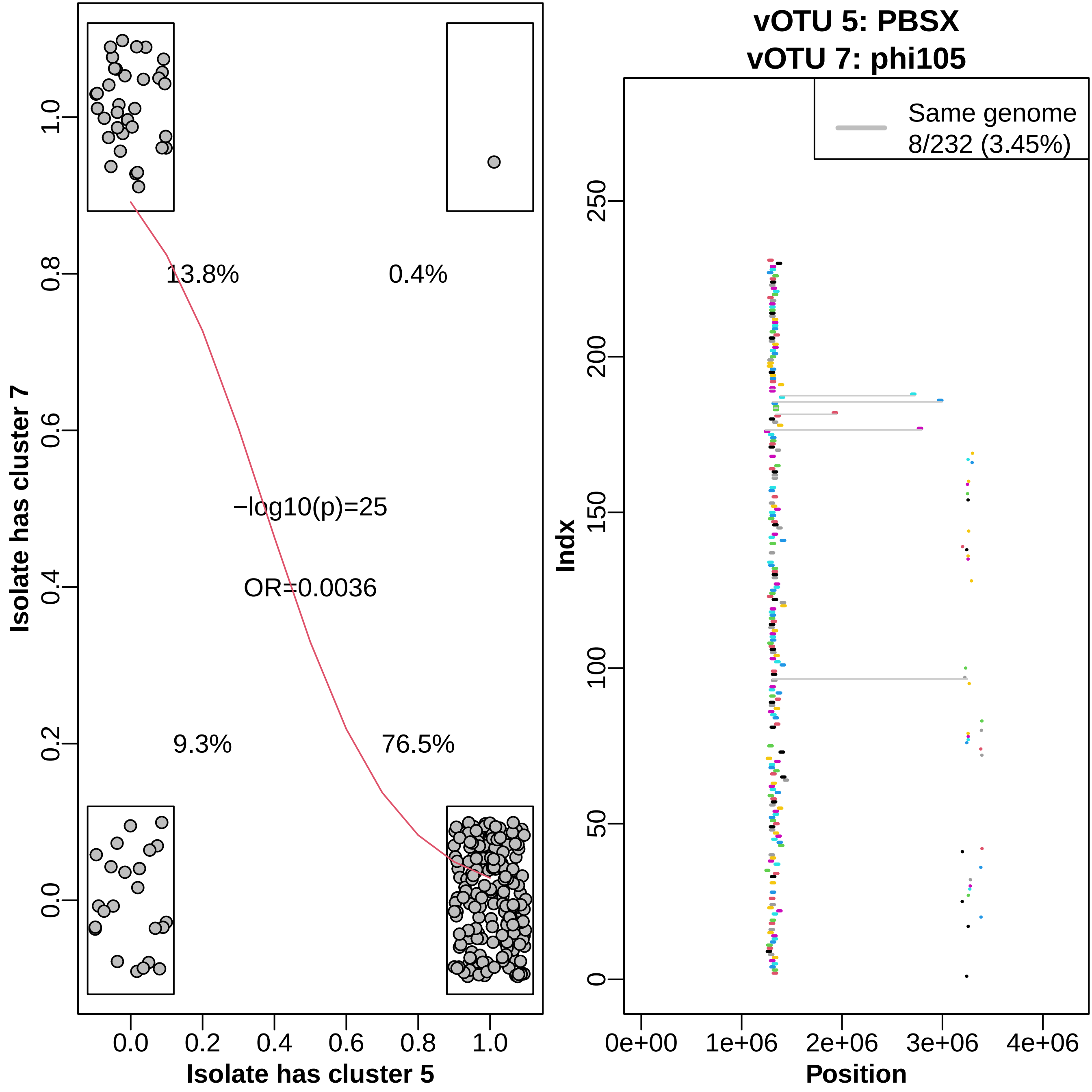

13.8%
0.4%
−log10(p)=25
OR=0.0036
9.3%
76.5%
Isolate has cluster 7
Isolate has cluster 5
0.0
0.2
0.4
0.6
0.8
1.0
vOTU 5: PBSX
vOTU 7: phi105
Same genome
8/232 (3.45%)
Indx
Position
0
50
100
150
200
250
0e+00
1e+06
2e+06
3e+06
4e+06

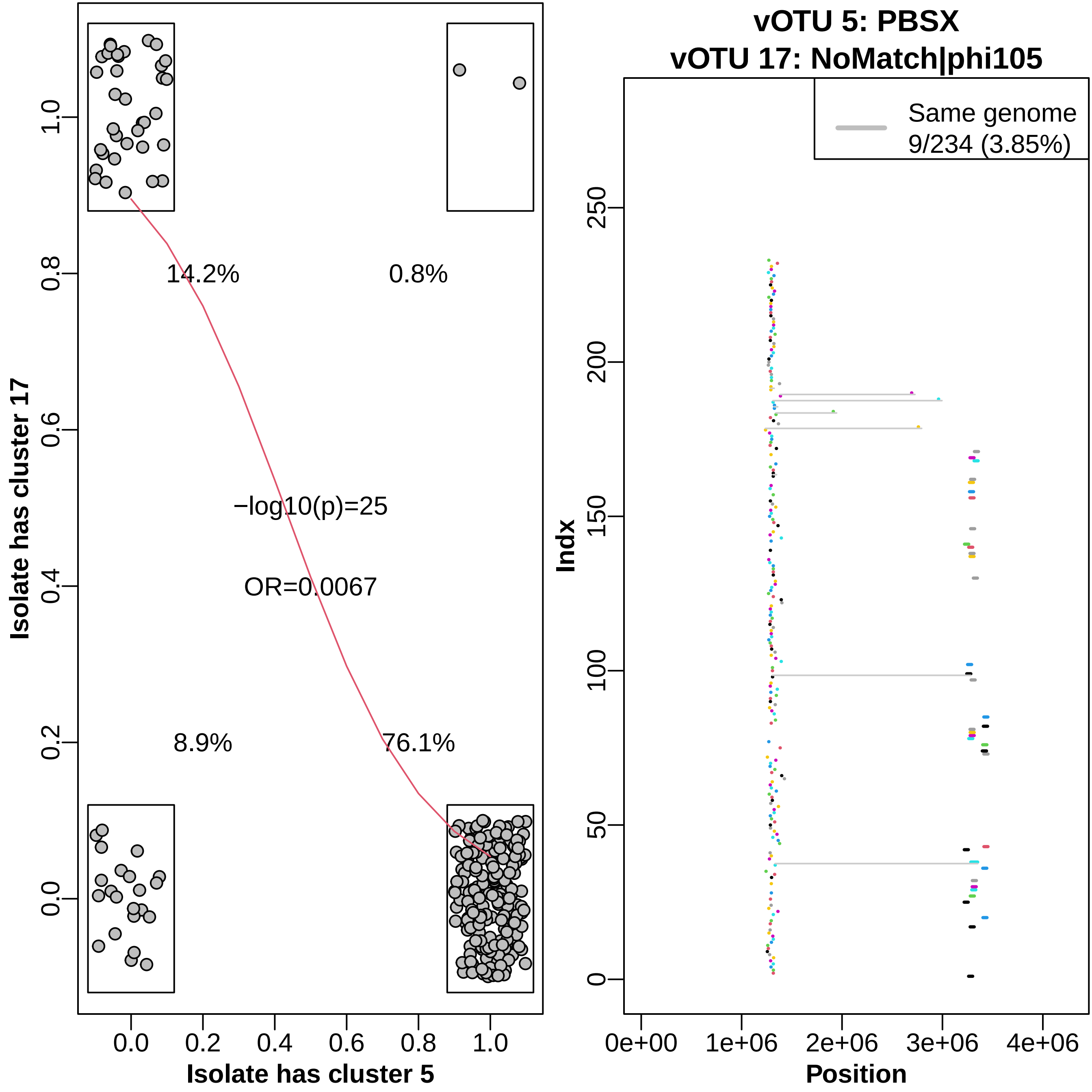

vOTU 5: PBSX
vOTU 17: NoMatch|phi105
14.2%
0.8%
−log10(p)=25
OR=0.0067
8.9%
76.1%
Isolate has cluster 17
Isolate has cluster 5
Same genome
9/234 (3.85%)
Indx
Position

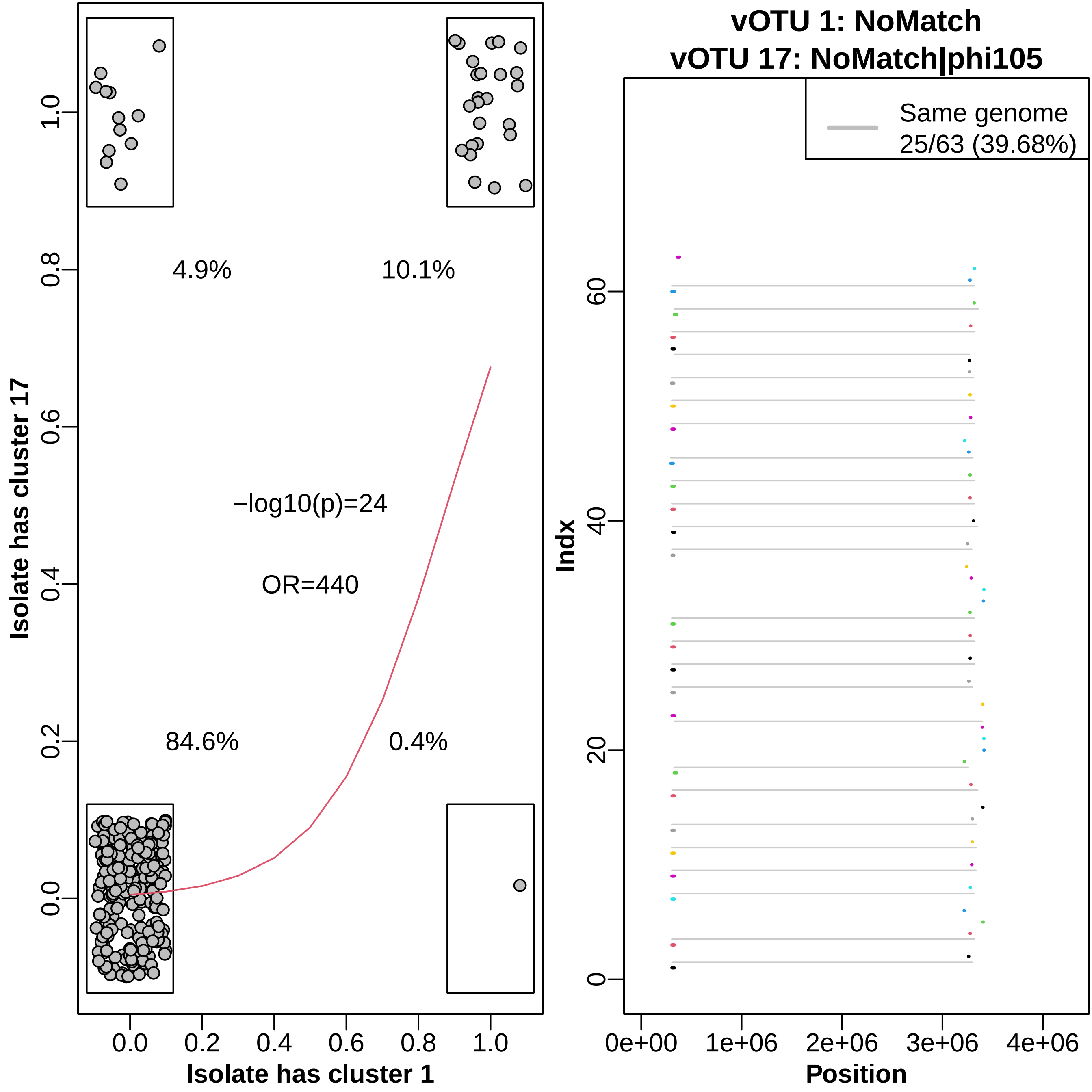

vOTU 1: NoMatch
vOTU 17: NoMatch|phi105
Same genome
25/63 (39.68%)
Indx
Position
0e+00
1e+06
2e+06
3e+06
4e+06
0
20
40
60
Isolate has cluster 17
Isolate has cluster 1
0.0
0.2
0.4
0.6
0.8
1.0
4.9%
10.1%
−log10(p)=24
OR=440
84.6%
0.4%

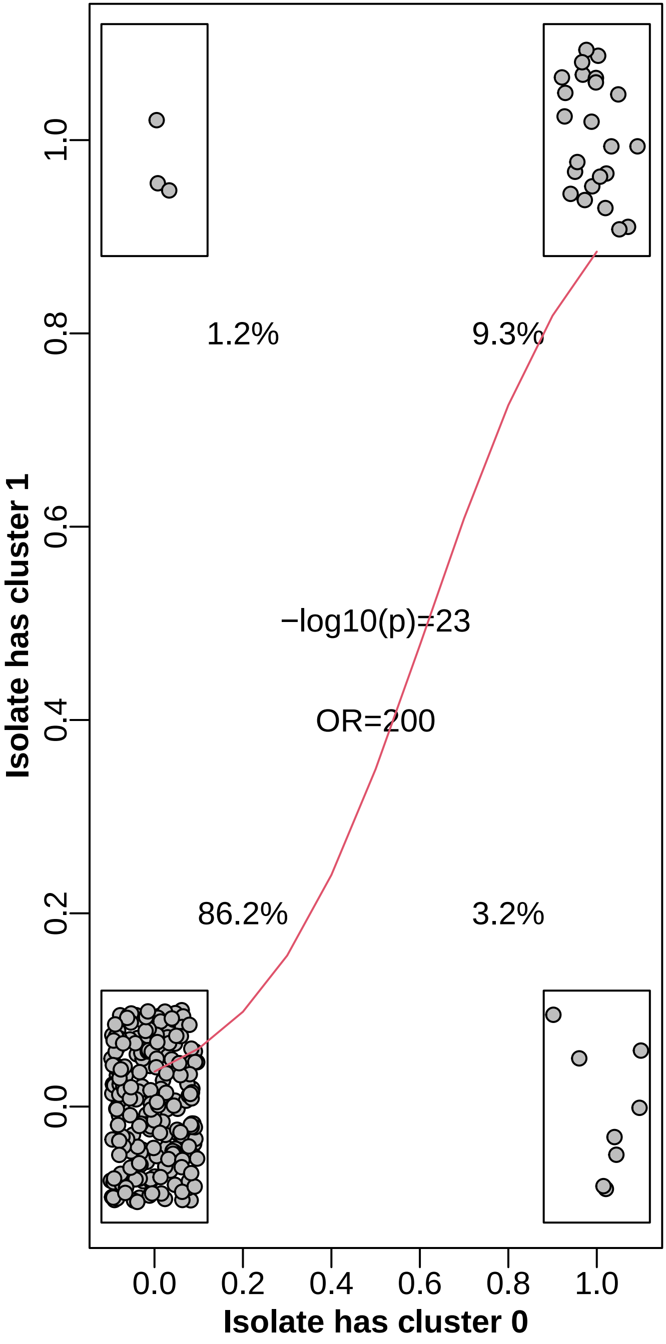

1.2%
9.3%
−log10(p)=23
OR=200
86.2%
3.2%
Isolate has cluster 1
Isolate has cluster 0
0.0
0.2
0.4
0.6
0.8
1.0


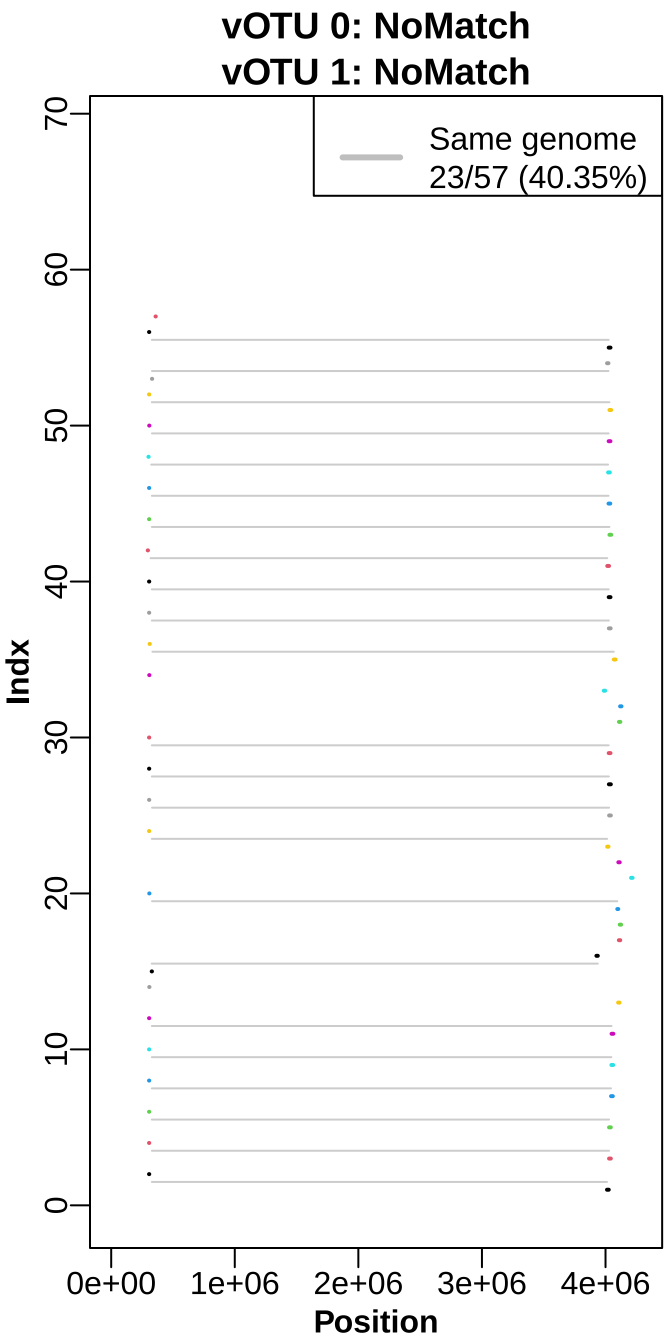

vOTU 0: NoMatch
vOTU 1: NoMatch
Same genome
23/57 (40.35%)
Indx
Position
0e+00
1e+06
2e+06
3e+06
4e+06
0
10
20
30
40
50
60
70

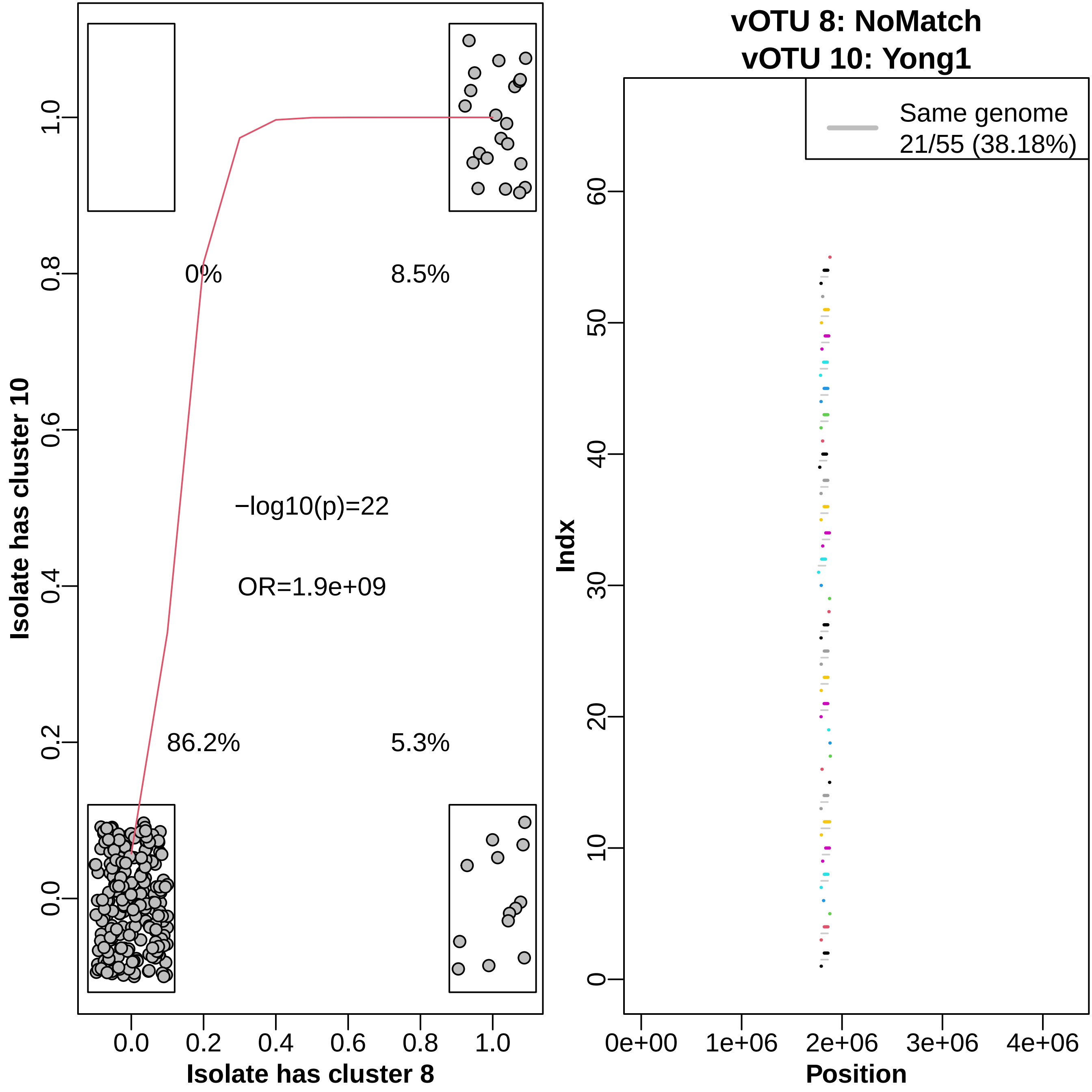

vOTU 8: NoMatch
vOTU 10: Yong1
Isolate has cluster 10
Isolate has cluster 8
0%
8.5%
86.2%
5.3%
−log10(p)=22
OR=1.9e+09
0.0
0.2
0.4
0.6
0.8
1.0
Same genome
21/55 (38.18%)
Indx
Position
0e+00
1e+06
2e+06
3e+06
4e+06
0
10
20
30
40
50
60

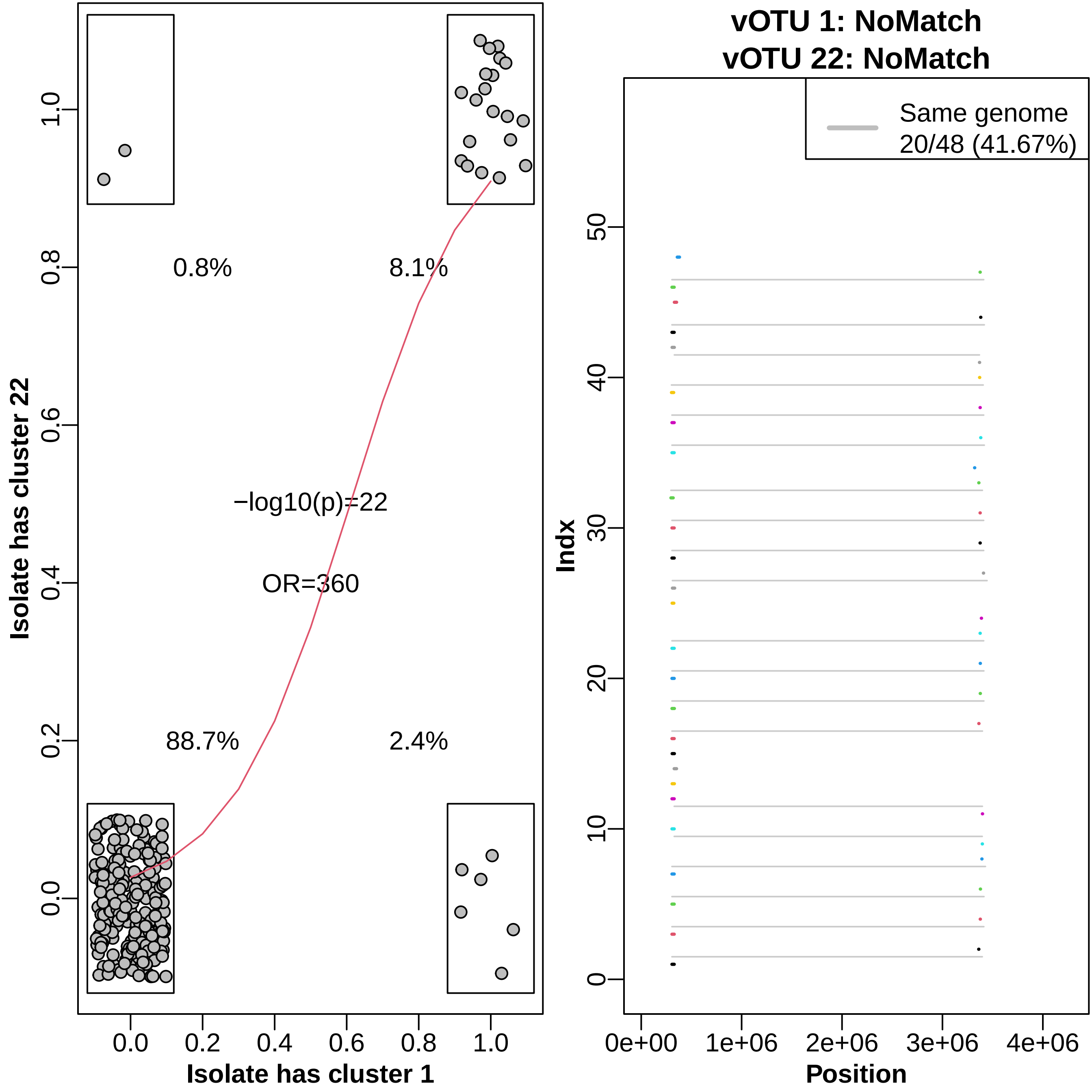

0.8%
8.1%
−log10(p)=22
OR=360
88.7%
2.4%
Isolate has cluster 22
Isolate has cluster 1
0.0
0.2
0.4
0.6
0.8
1.0
vOTU 1: NoMatch
vOTU 22: NoMatch
Same genome
20/48 (41.67%)
Indx
0
10
20
30
40
50
Position
0e+00
1e+06
2e+06
3e+06
4e+06

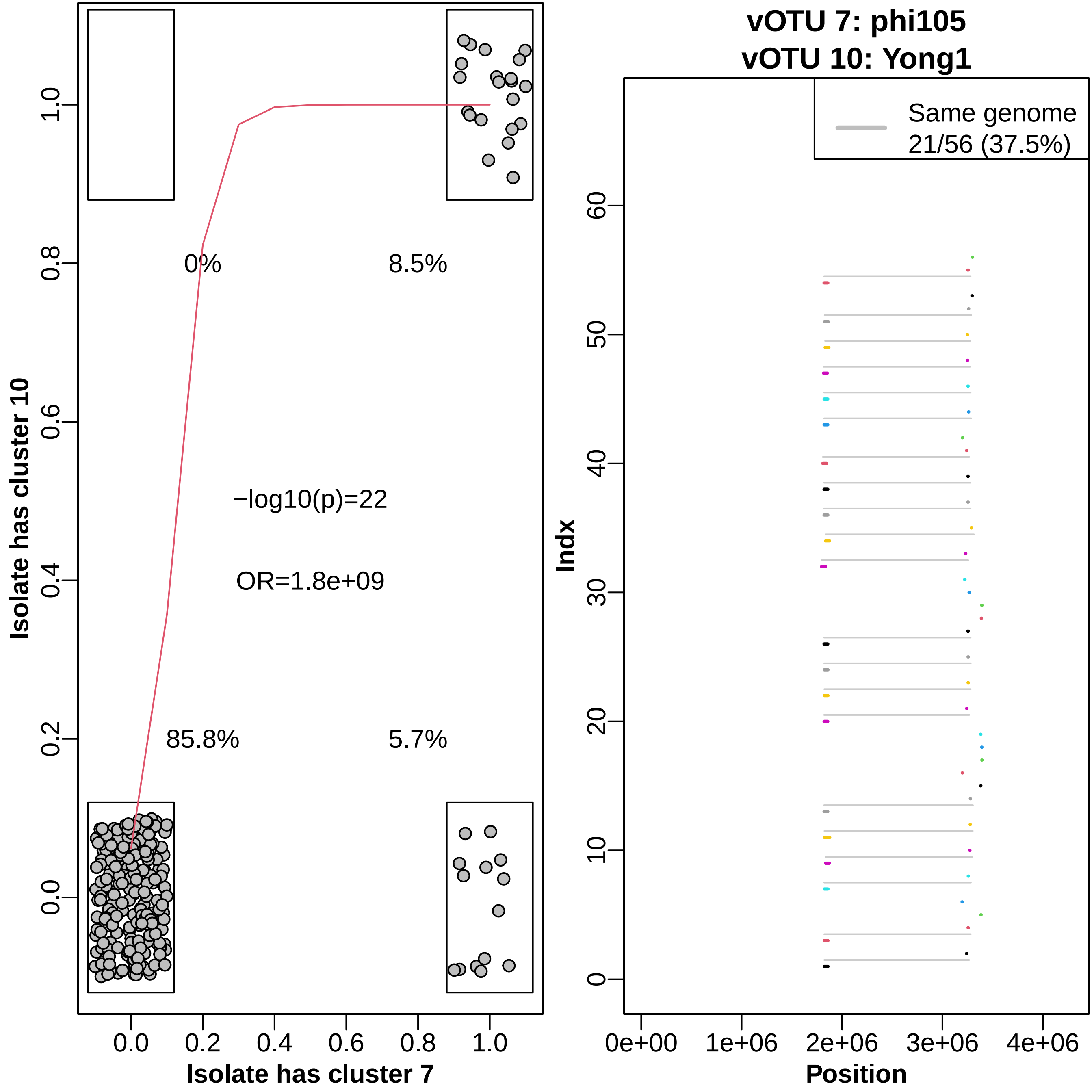

vOTU 7: phi105
vOTU 10: Yong1
Same genome
21/56 (37.5%)
0%
8.5%
85.8%
5.7%
−log10(p)=22
OR=1.8e+09
Isolate has cluster 10
Isolate has cluster 7
Indx
Position
0.0
0.2
0.4
0.6
0.8
1.0
0e+00
1e+06
2e+06
3e+06
4e+06
0
10
20
30
40
50
60

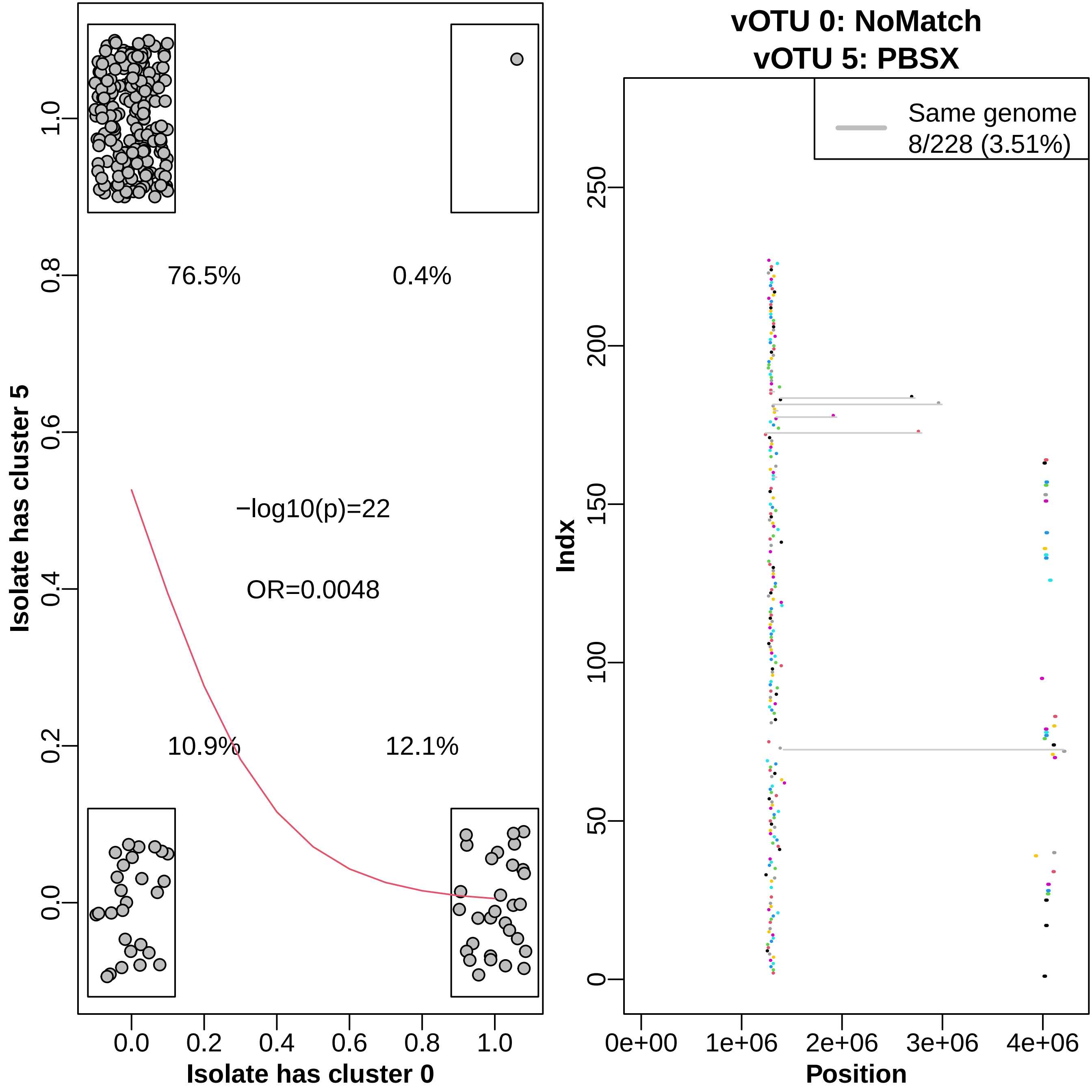

76.5%
0.4%
−log10(p)=22
OR=0.0048
10.9%
12.1%
Isolate has cluster 5
Isolate has cluster 0
0.0
0.2
0.4
0.6
0.8
1.0
vOTU 0: NoMatch
vOTU 5: PBSX
Same genome
8/228 (3.51%)
Indx
Position
0
50
100
150
200
250
0e+00
1e+06
2e+06
3e+06
4e+06

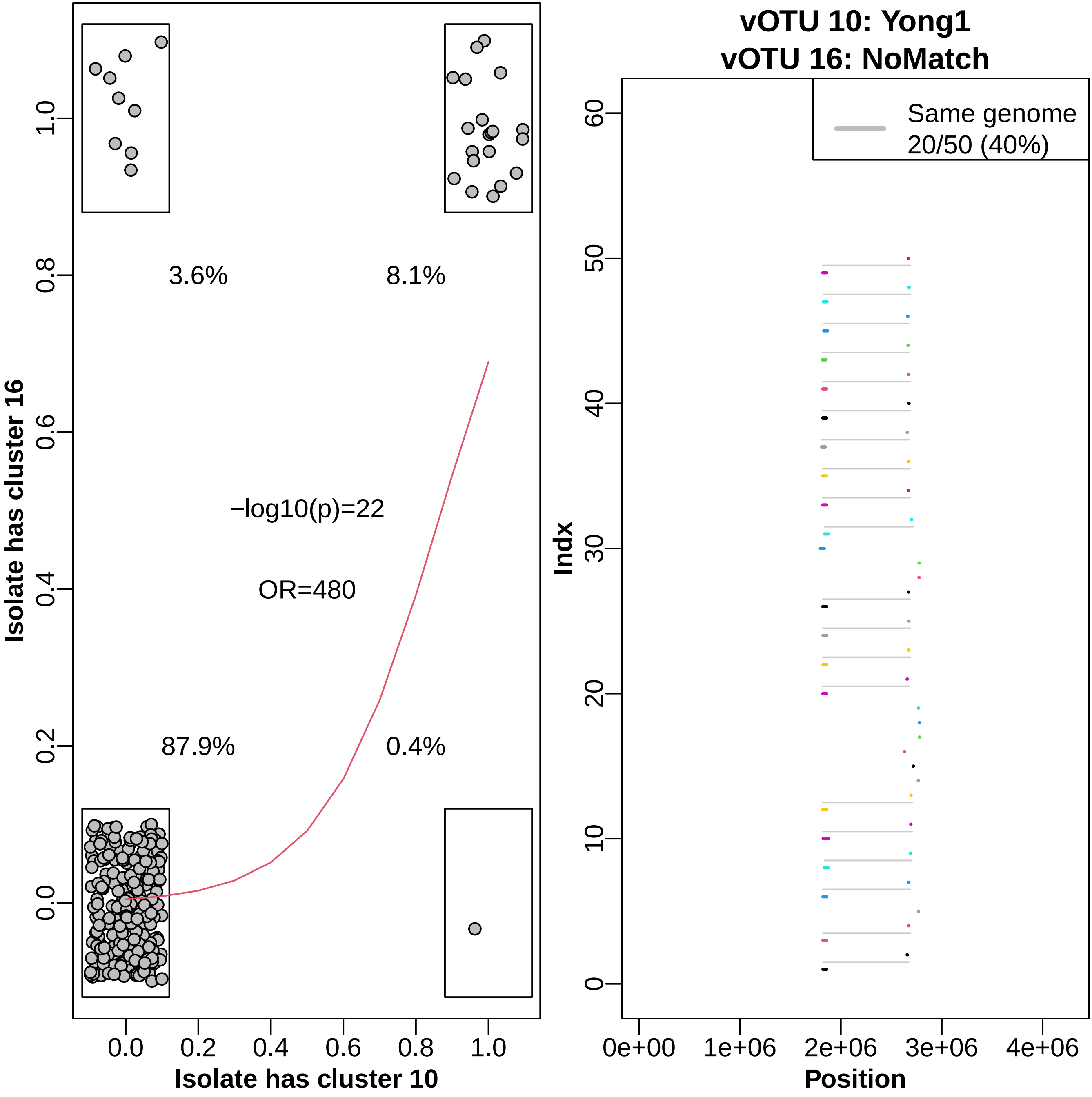

3.6%
8.1%
−log10(p)=22
OR=480
87.9%
0.4%
Isolate has cluster 16
Isolate has cluster 10
vOTU 10: Yong1
vOTU 16: NoMatch
Same genome
20/50 (40%)
Indx
Position

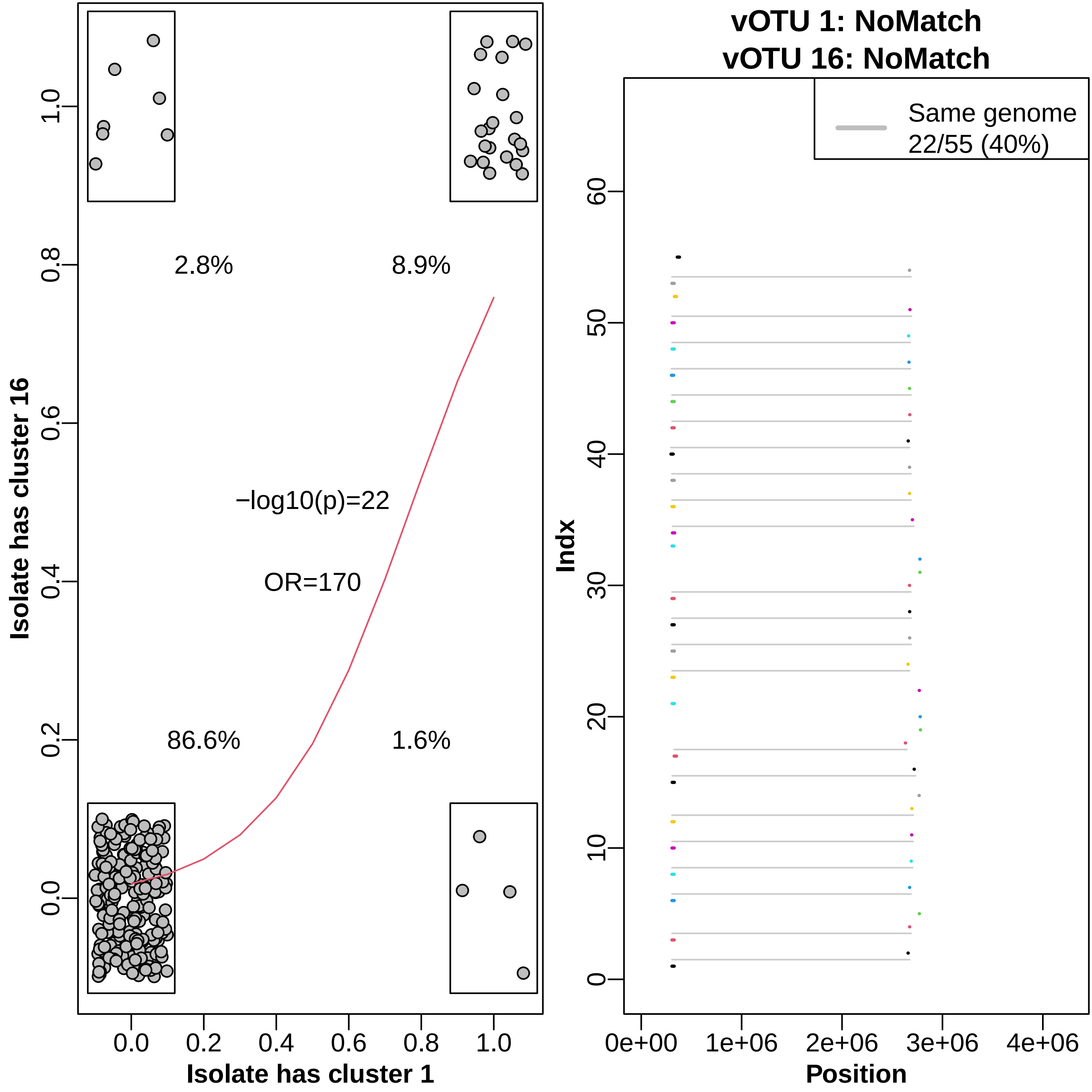
2.8%
8.9%
−log10(p)=22
OR=170
86.6%
1.6%
Isolate has cluster 16
Isolate has cluster 1
vOTU 1: NoMatch
vOTU 16: NoMatch
Same genome
22/55 (40%)
Indx
Position

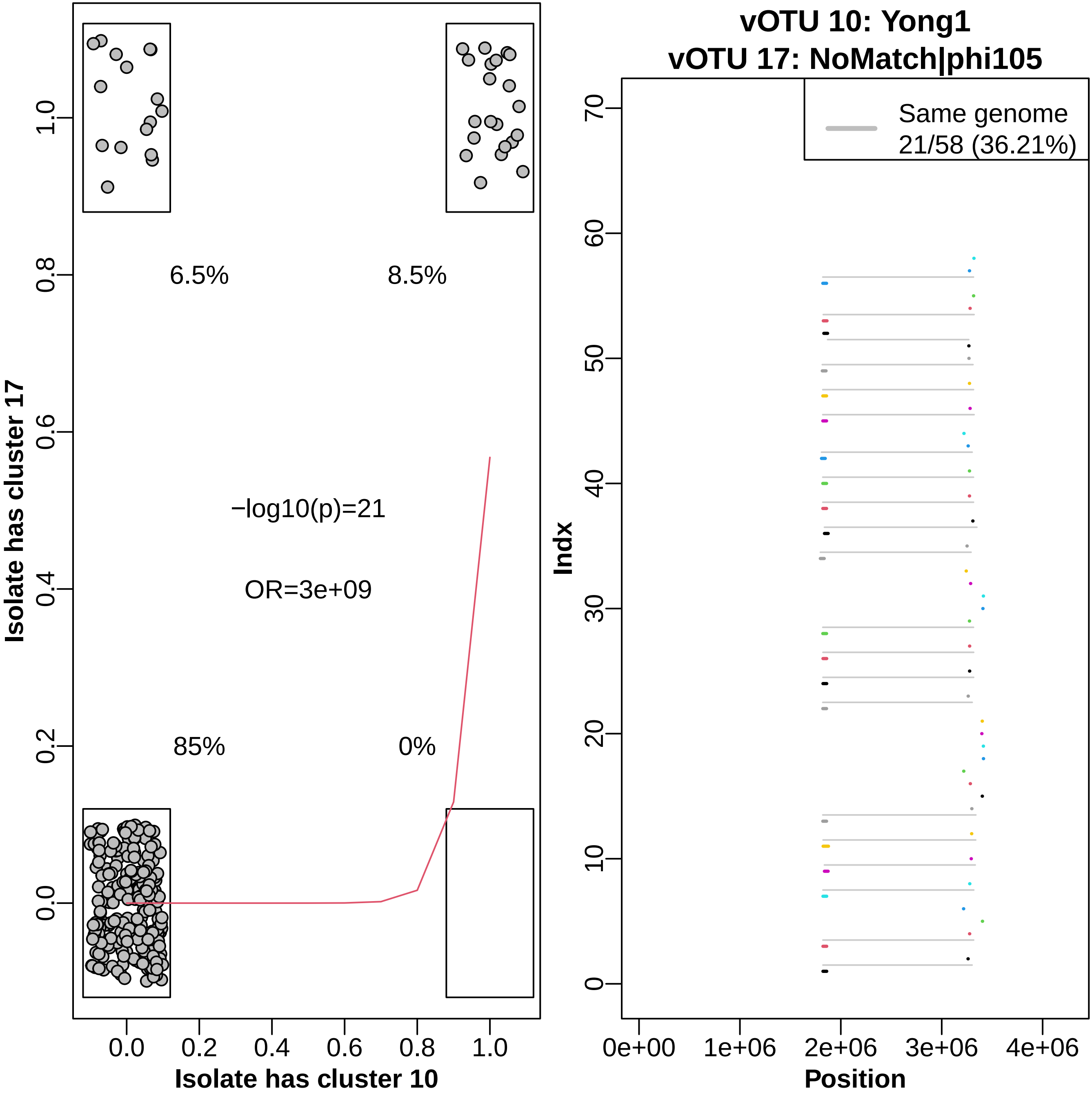

Isolate has cluster 17
Isolate has cluster 10
6.5%
8.5%
85%
0%
−log10(p)=21
OR=3e+09
0.0
0.2
0.4
0.6
0.8
1.0
vOTU 10: Yong1
vOTU 17: NoMatch|phi105
Same genome
21/58 (36.21%)
Indx
Position
0e+00
1e+06
2e+06
3e+06
4e+06
0
10
20
30
40
50
60
70

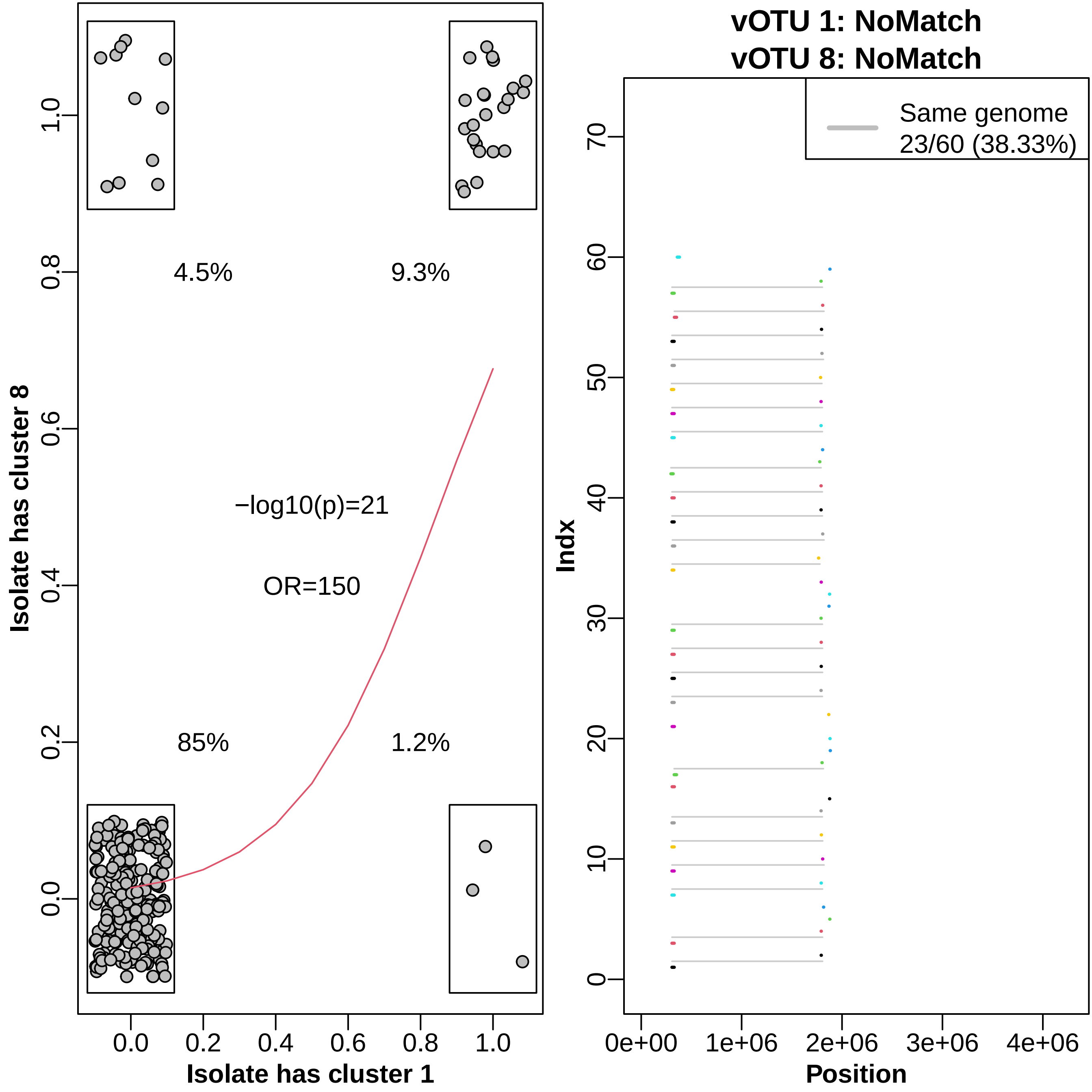

4.5%
9.3%
−log10(p)=21
OR=150
85%
1.2%
Isolate has cluster 8
Isolate has cluster 1
0.0
0.2
0.4
0.6
0.8
1.0
vOTU 1: NoMatch
vOTU 8: NoMatch
Same genome
23/60 (38.33%)
Indx
Position
0
10
20
30
40
50
60
70
0e+00
1e+06
2e+06
3e+06
4e+06

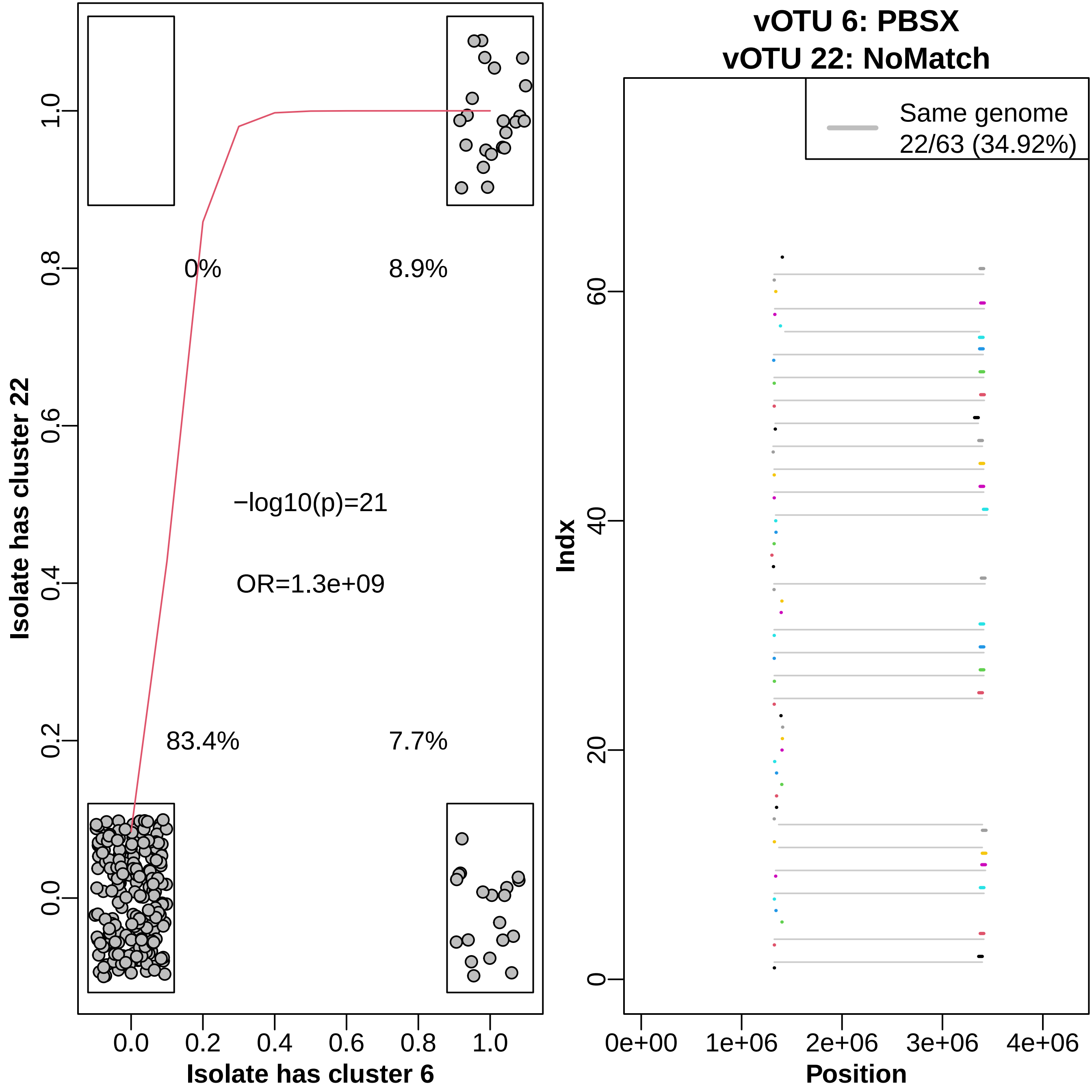

Isolate has cluster 22
Isolate has cluster 6
0%
8.9%
83.4%
7.7%
−log10(p)=21
OR=1.3e+09
vOTU 6: PBSX
vOTU 22: NoMatch
Same genome
22/63 (34.92%)
Indx
Position

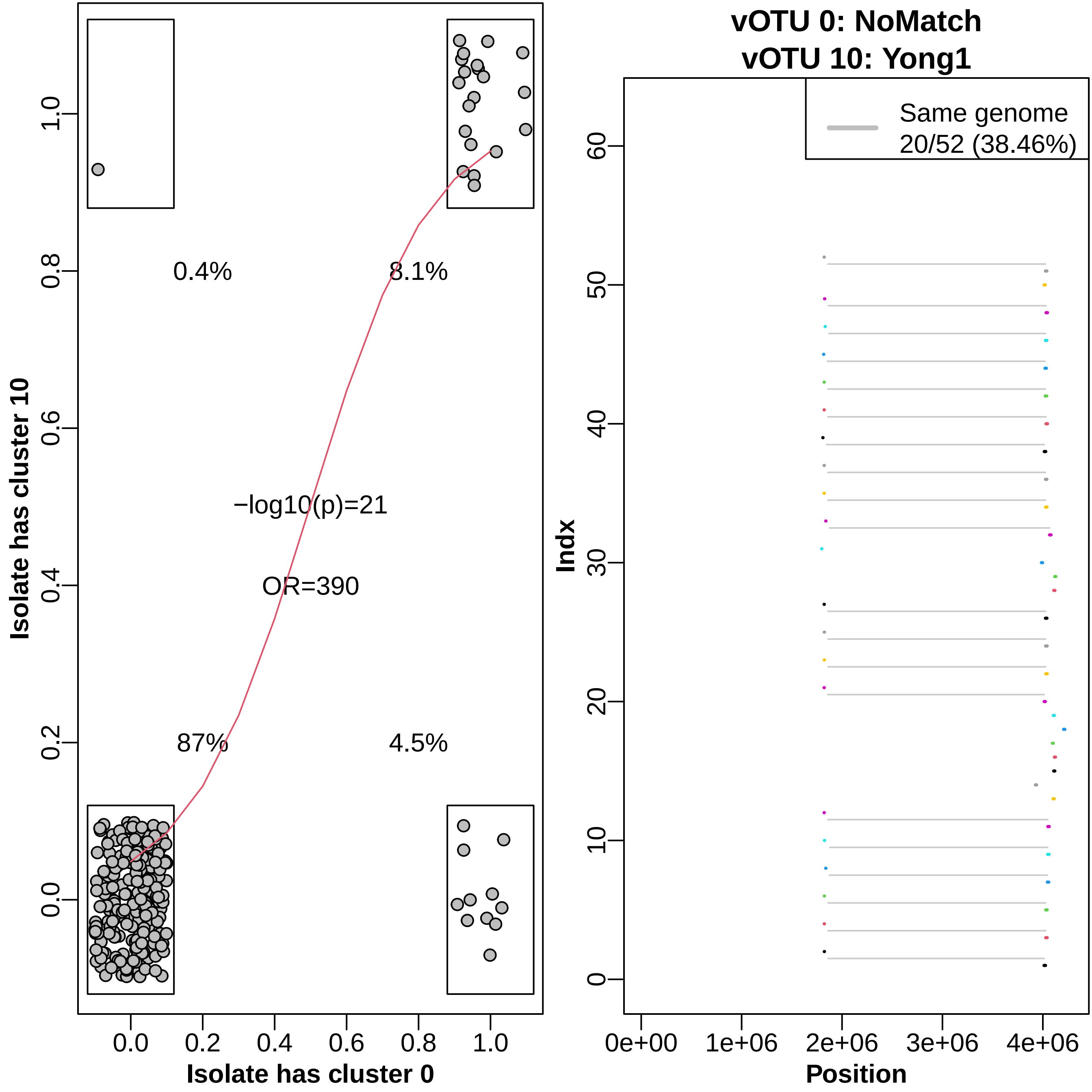

vOTU 0: NoMatch
vOTU 10: Yong1
Isolate has cluster 10
Isolate has cluster 0
0.4%
8.1%
−log10(p)=21
OR=390
87%
4.5%
0.0
0.2
0.4
0.6
0.8
1.0
Same genome
20/52 (38.46%)
Indx
Position
0e+00
1e+06
2e+06
3e+06
4e+06
0
10
20
30
40
50
60

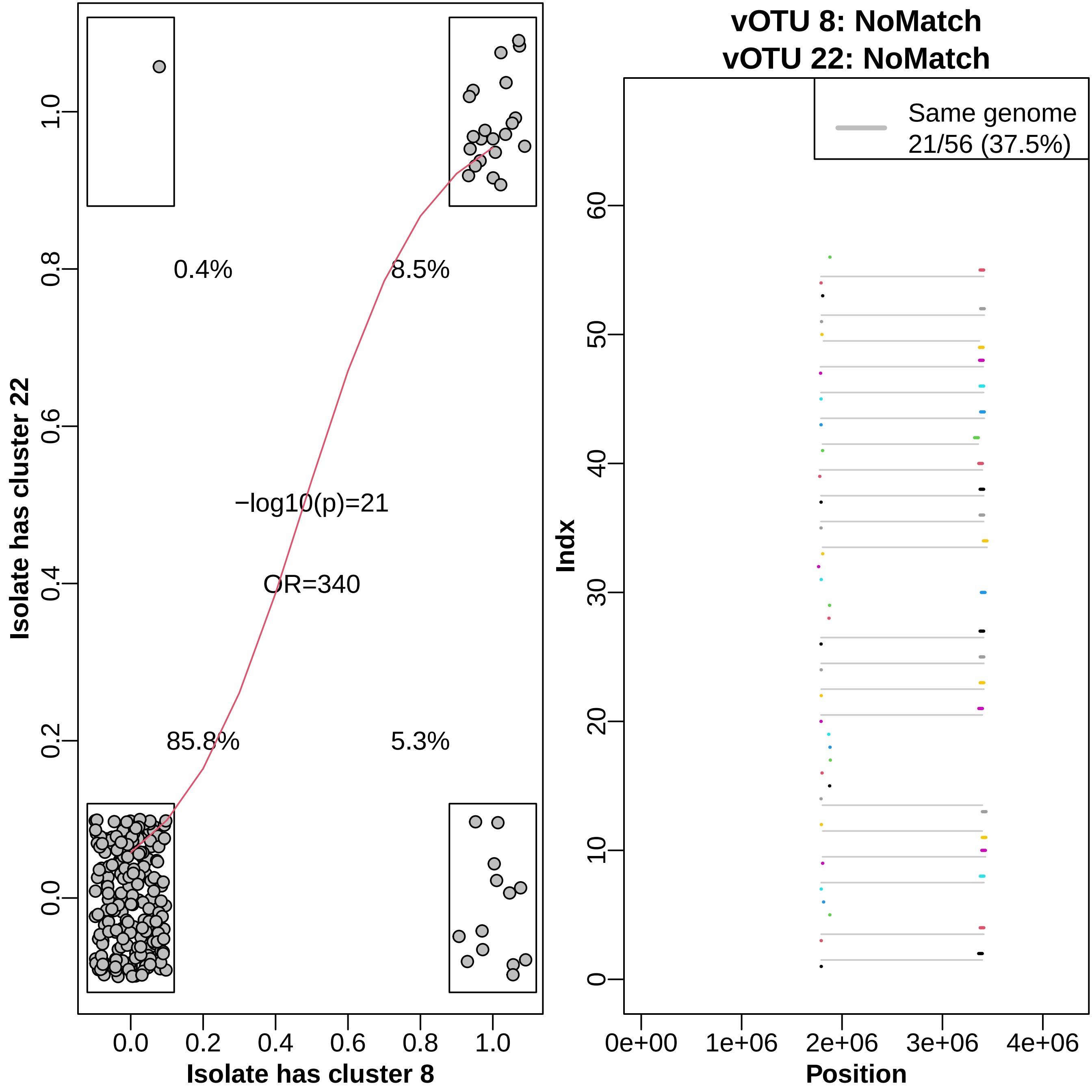

vOTU 8: NoMatch
vOTU 22: NoMatch
Same genome
21/56 (37.5%)
Isolate has cluster 22
Isolate has cluster 8
0.4%
8.5%
85.8%
5.3%
−log10(p)=21
OR=340
0.0
0.2
0.4
0.6
0.8
1.0
Indx
Position
0
10
20
30
40
50
60
0e+00
1e+06
2e+06
3e+06
4e+06

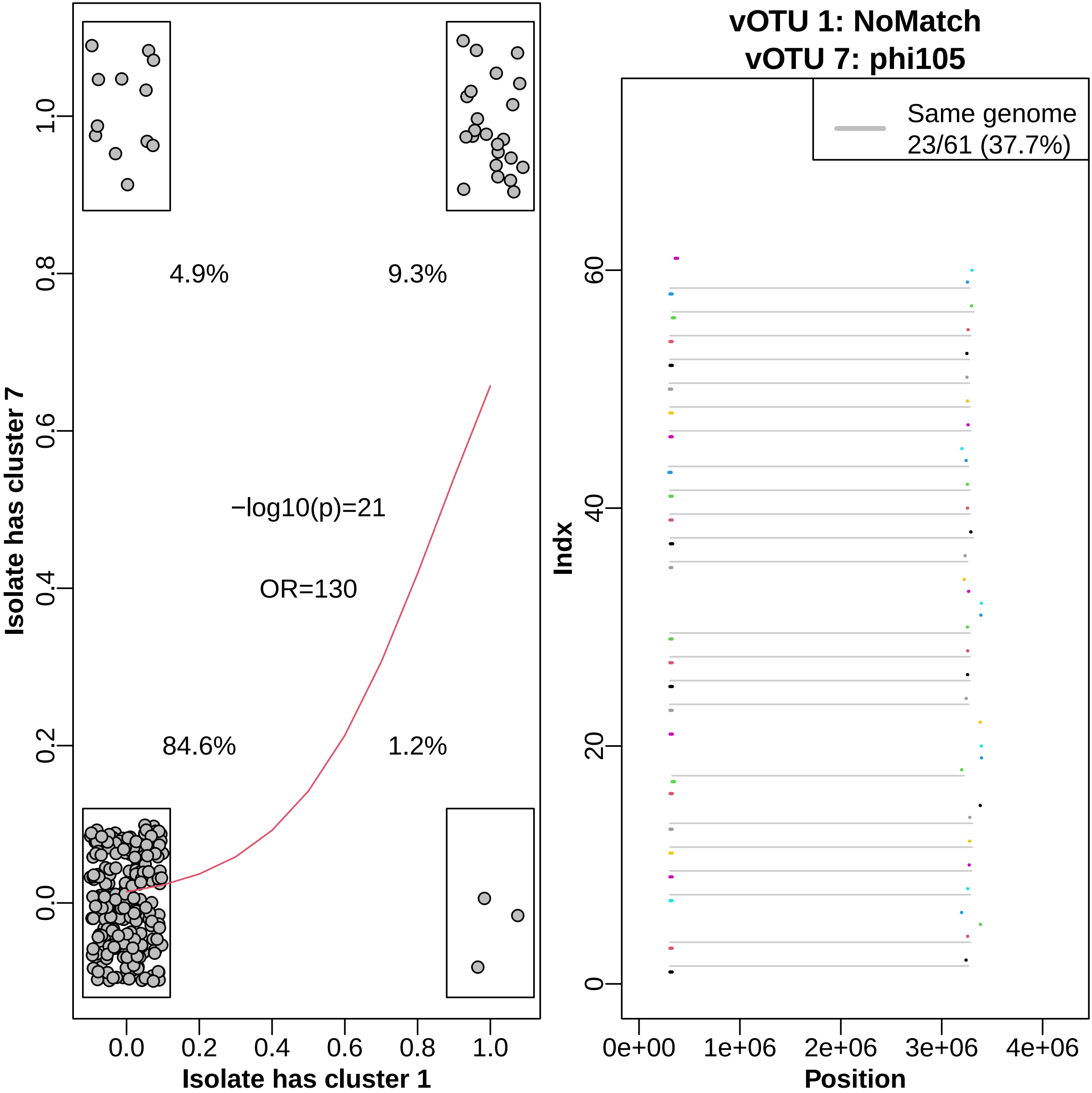

vOTU 1: NoMatch
vOTU 7: phi105
Isolate has cluster 7
Isolate has cluster 1
4.9%
9.3%
84.6%
1.2%
−log10(p)=21
OR=130
Same genome
23/61 (37.7%)
Indx
Position
0e+00
1e+06
2e+06
3e+06
4e+06

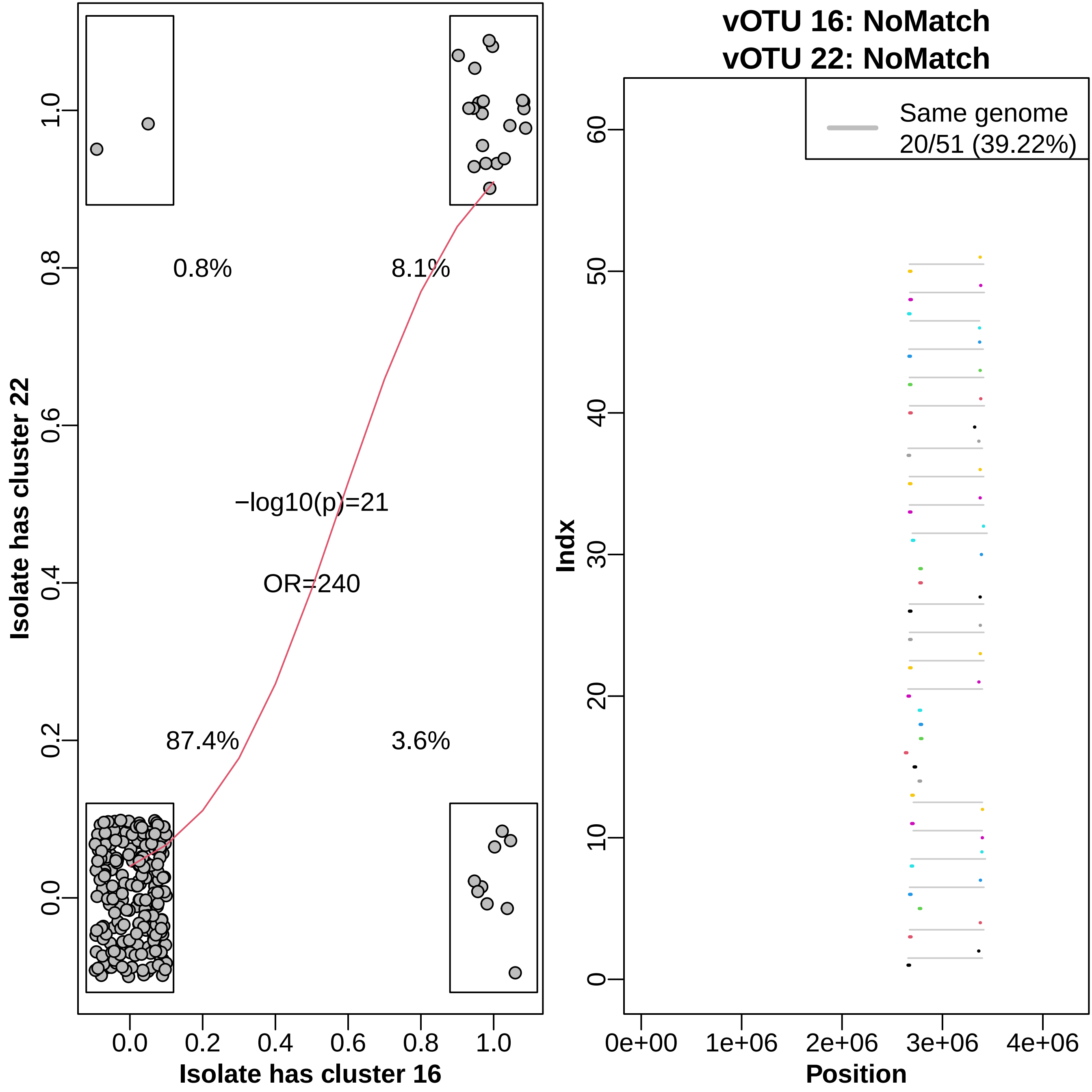

vOTU 16: NoMatch
vOTU 22: NoMatch
Isolate has cluster 22
Isolate has cluster 16
0.8%
8.1%
87.4%
3.6%
−log10(p)=21
OR=240
0.0
0.2
0.4
0.6
0.8
1.0
Same genome
20/51 (39.22%)
Indx
Position
0e+00
1e+06
2e+06
3e+06
4e+06
0
10
20
30
40
50
60

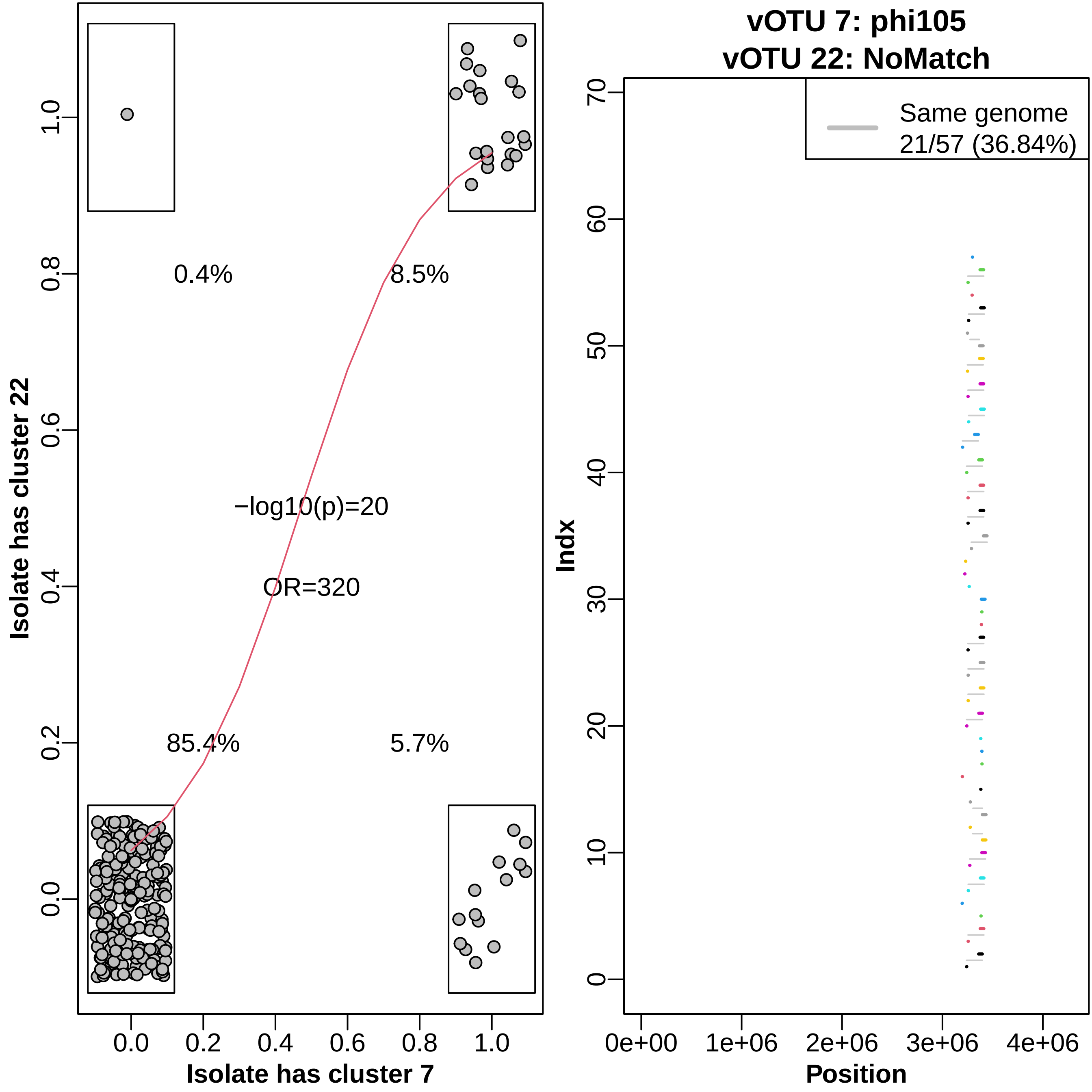

vOTU 7: phi105
vOTU 22: NoMatch
0.4%
8.5%
−log10(p)=20
OR=320
85.4%
5.7%
Isolate has cluster 22
Isolate has cluster 7
0.0
0.2
0.4
0.6
0.8
1.0
Same genome
21/57 (36.84%)
Indx
Position
0e+00
1e+06
2e+06
3e+06
4e+06
0
10
20
30
40
50
60
70

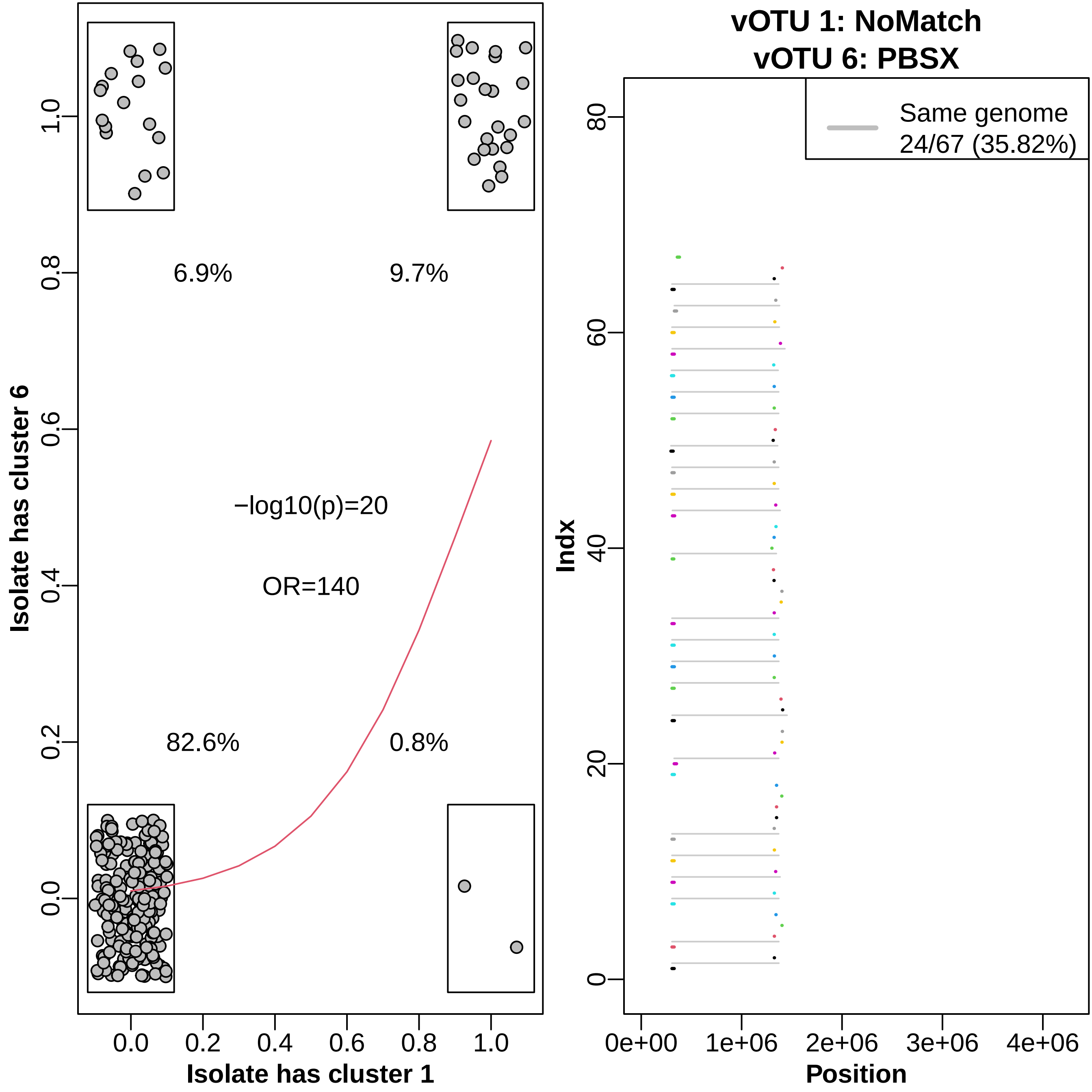

6.9%
9.7%
−log10(p)=20
OR=140
82.6%
0.8%
Isolate has cluster 6
Isolate has cluster 1
vOTU 1: NoMatch
vOTU 6: PBSX
Same genome
24/67 (35.82%)
Indx
Position

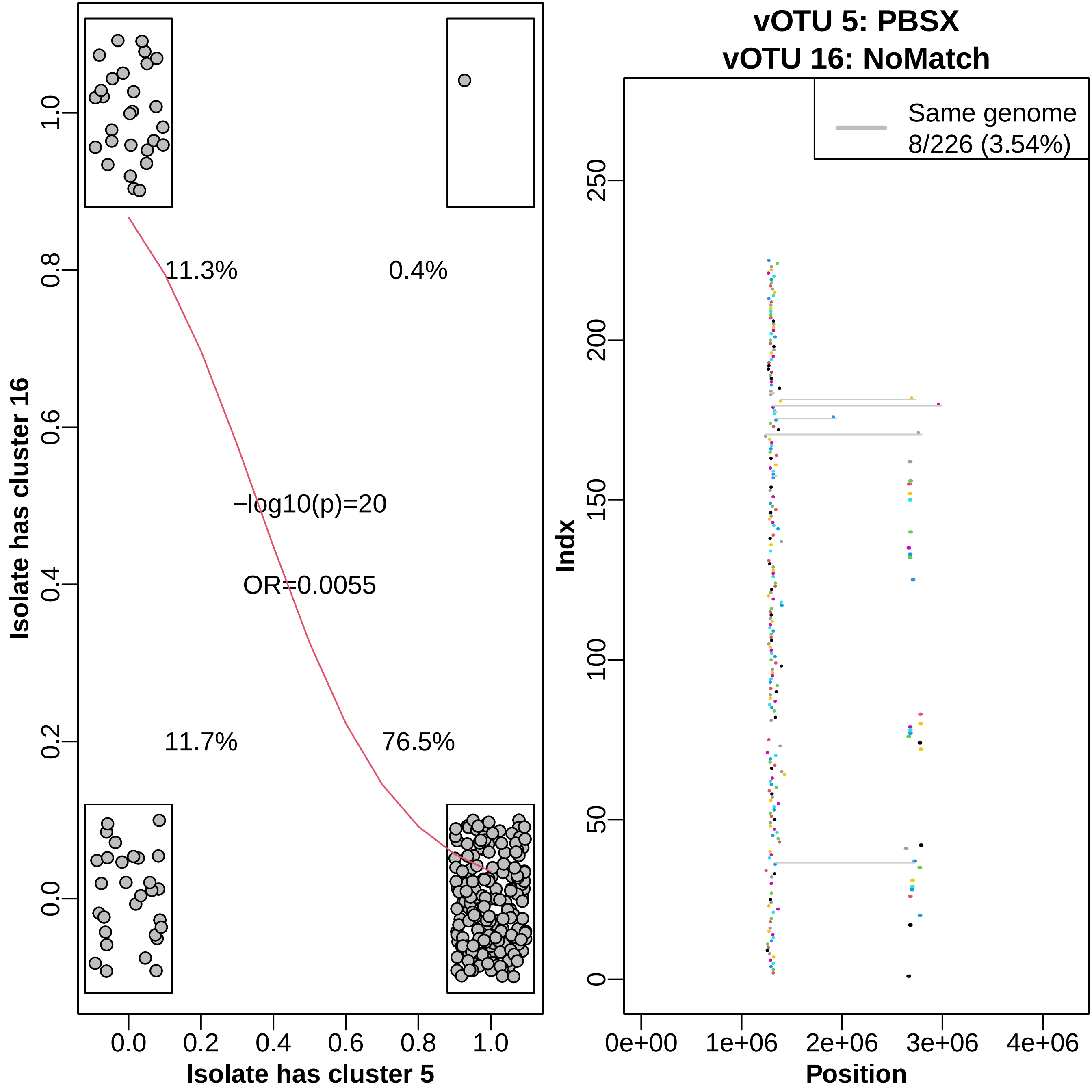

Isolate has cluster 16
Isolate has cluster 5
11.3%
0.4%
11.7%
76.5%
−log10(p)=20
OR=0.0055
0.0
0.2
0.4
0.6
0.8
1.0
vOTU 5: PBSX
vOTU 16: NoMatch
Same genome
8/226 (3.54%)
Indx
Position
0
50
100
150
200
250
0e+00
1e+06
2e+06
3e+06
4e+06

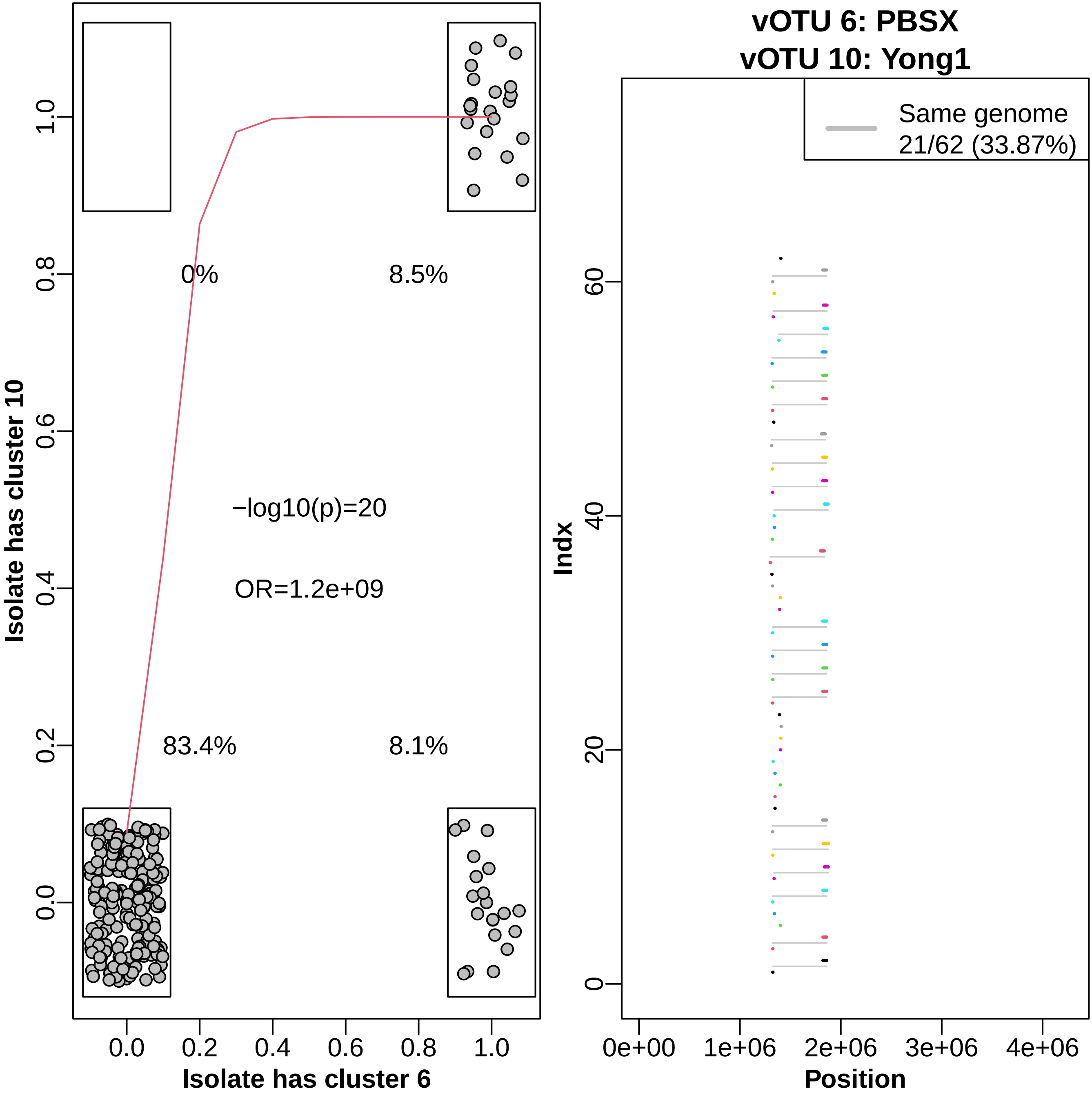

vOTU 6: PBSX
vOTU 10: Yong1
Isolate has cluster 10
Isolate has cluster 6
0%
8.5%
83.4%
8.1%
−log10(p)=20
OR=1.2e+09
0.0
0.2
0.4
0.6
0.8
1.0
Same genome
21/62 (33.87%)
Indx
Position
0
20
40
60
0e+00
1e+06
2e+06
3e+06
4e+06

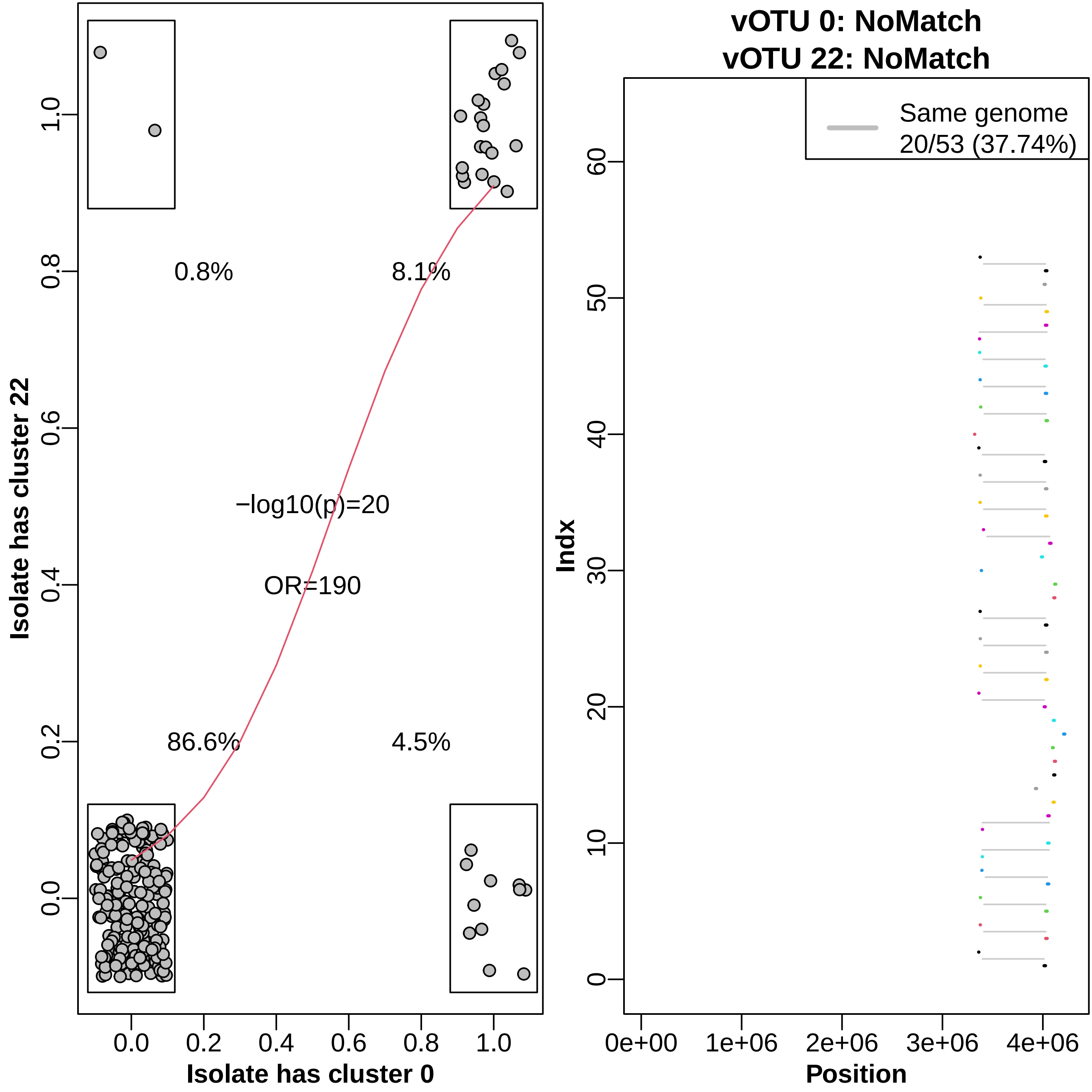

0.8%
8.1%
−log10(p)=20
OR=190
86.6%
4.5%
Isolate has cluster 22
Isolate has cluster 0
0.0
0.2
0.4
0.6
0.8
1.0
vOTU 0: NoMatch
vOTU 22: NoMatch
Same genome
20/53 (37.74%)
Indx
Position
0
10
20
30
40
50
60
0e+00
1e+06
2e+06
3e+06
4e+06

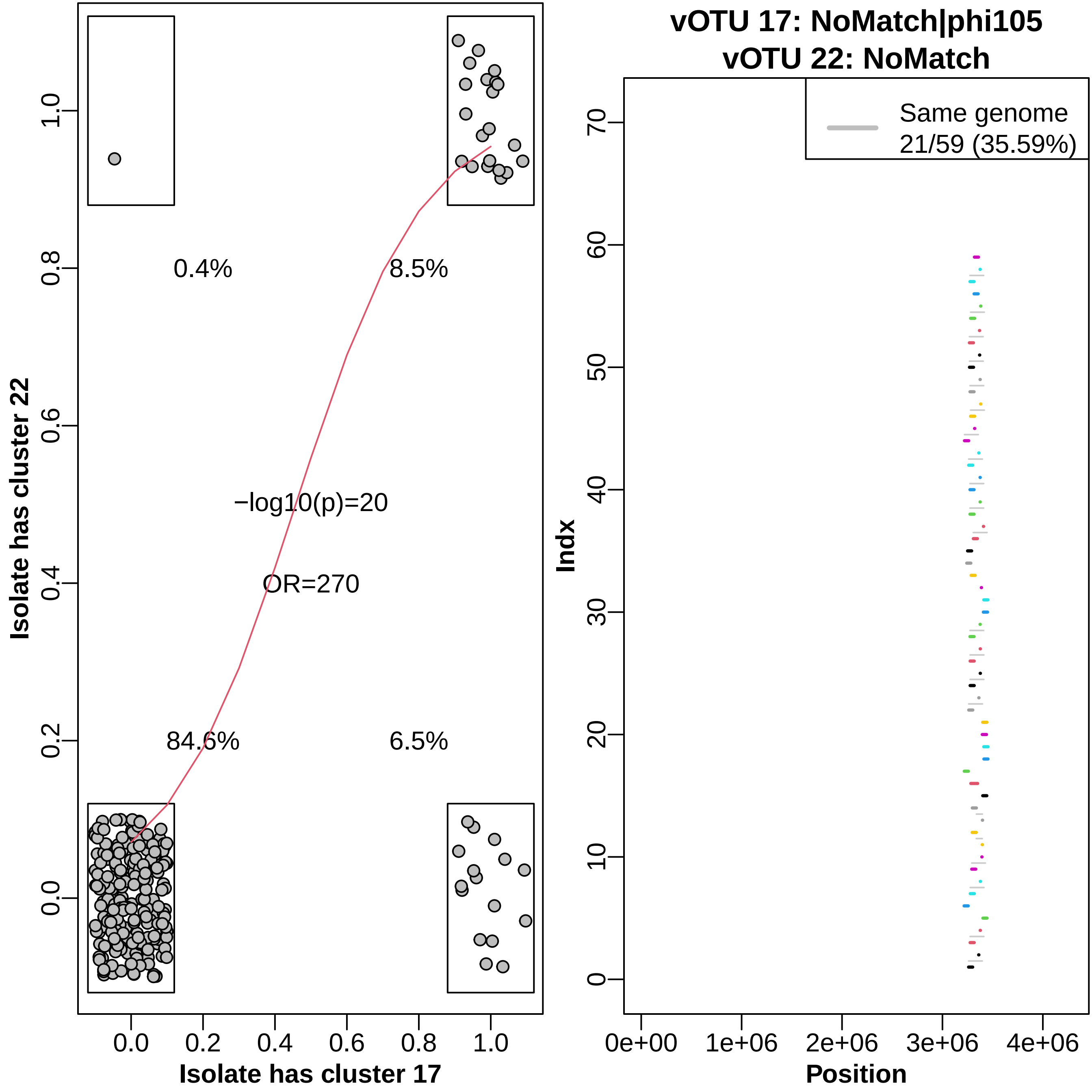

vOTU 17: NoMatch|phi105
vOTU 22: NoMatch
Isolate has cluster 22
Isolate has cluster 17
0.4%
8.5%
84.6%
6.5%
−log10(p)=20
OR=270
Same genome
21/59 (35.59%)
Indx
Position
0e+00
1e+06
2e+06
3e+06
4e+06

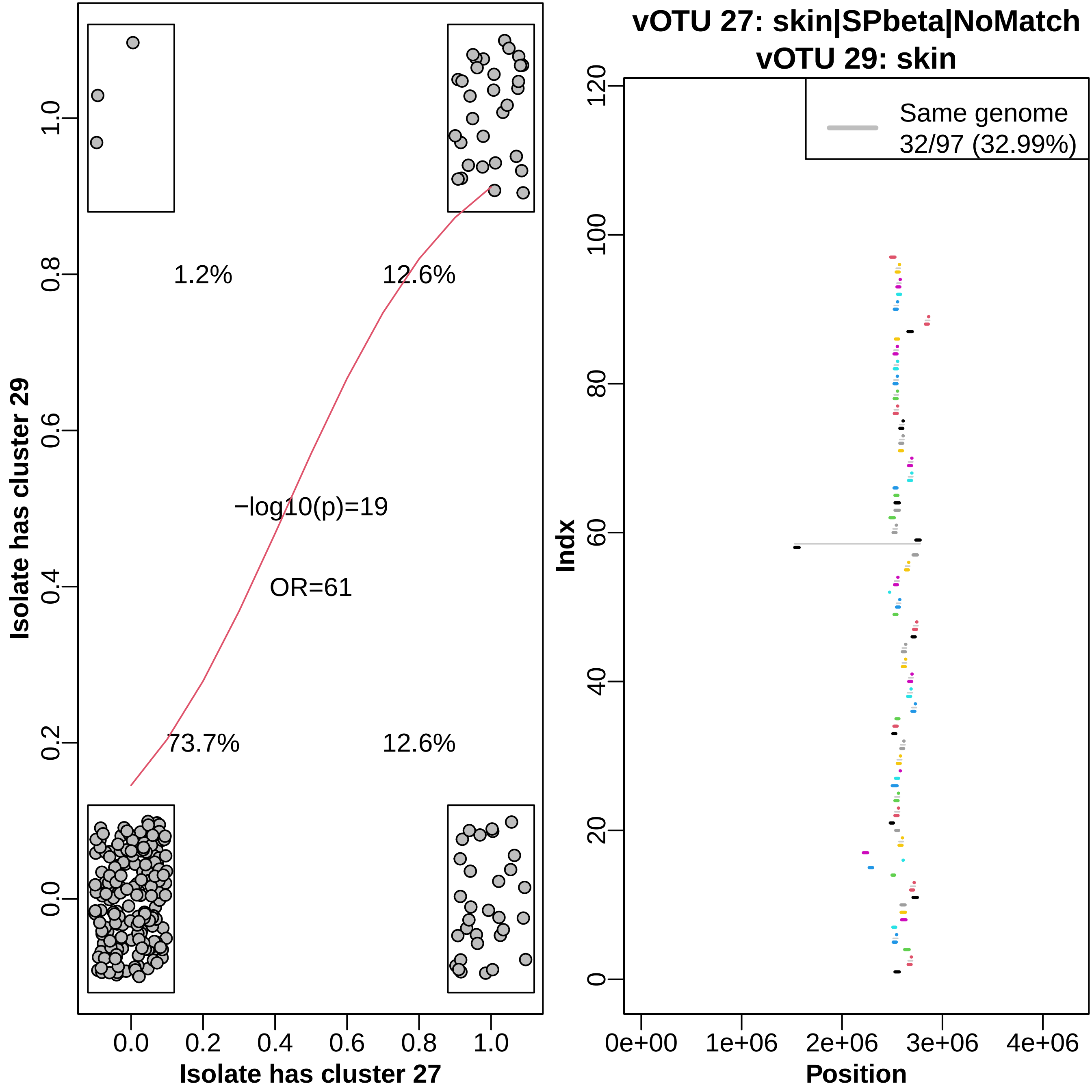

vOTU 27: skin|SPbeta|NoMatch
vOTU 29: skin
1.2%
12.6%
−log10(p)=19
OR=61
73.7%
12.6%
Isolate has cluster 29
Isolate has cluster 27
0.0
0.2
0.4
0.6
0.8
1.0
Same genome
32/97 (32.99%)
Indx
Position
0e+00
1e+06
2e+06
3e+06
4e+06
0
20
40
60
80
100
120

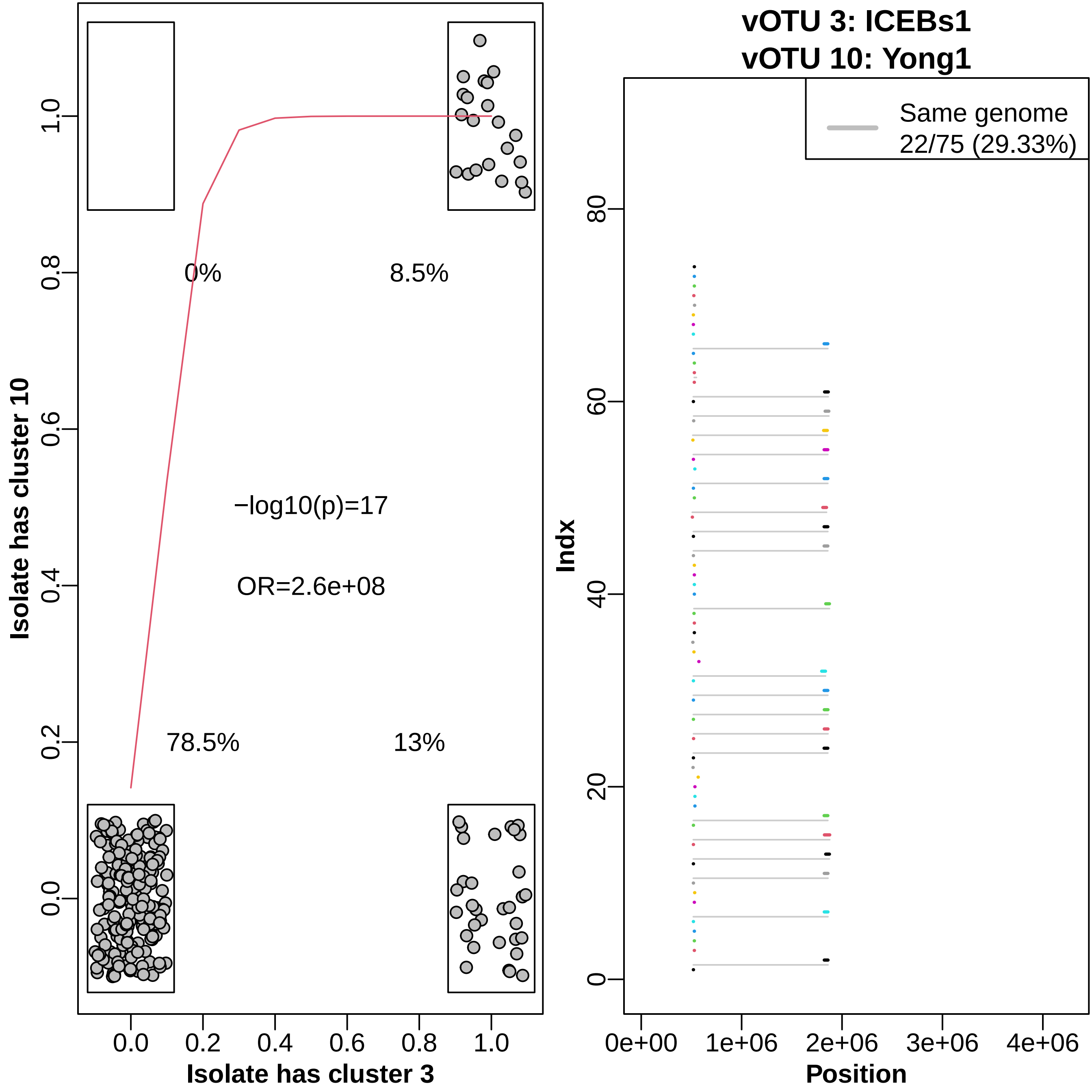

vOTU 3: ICEBs1
vOTU 10: Yong1
Isolate has cluster 10
Isolate has cluster 3
0%
8.5%
78.5%
13%
−log10(p)=17
OR=2.6e+08
0.0
0.2
0.4
0.6
0.8
1.0
Same genome
22/75 (29.33%)
Indx
Position
0e+00
1e+06
2e+06
3e+06
4e+06
0
20
40
60
80

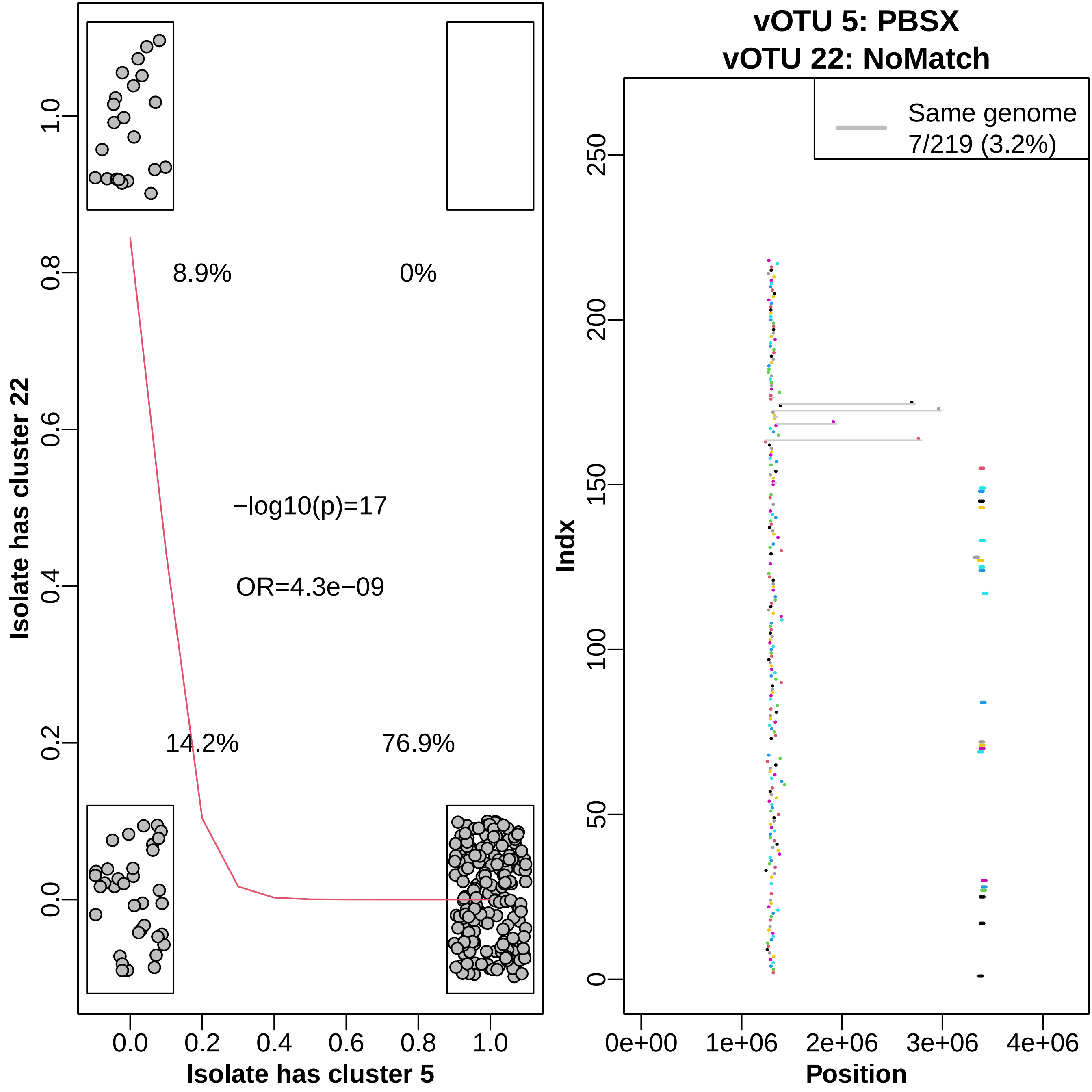

Isolate has cluster 22
Isolate has cluster 5
8.9%
0%
14.2%
76.9%
−log10(p)=17
OR=4.3e−09
0.0
0.2
0.4
0.6
0.8
1.0
vOTU 5: PBSX
vOTU 22: NoMatch
Same genome
7/219 (3.2%)
Indx
Position
0
50
100
150
200
250
0e+00
1e+06
2e+06
3e+06
4e+06

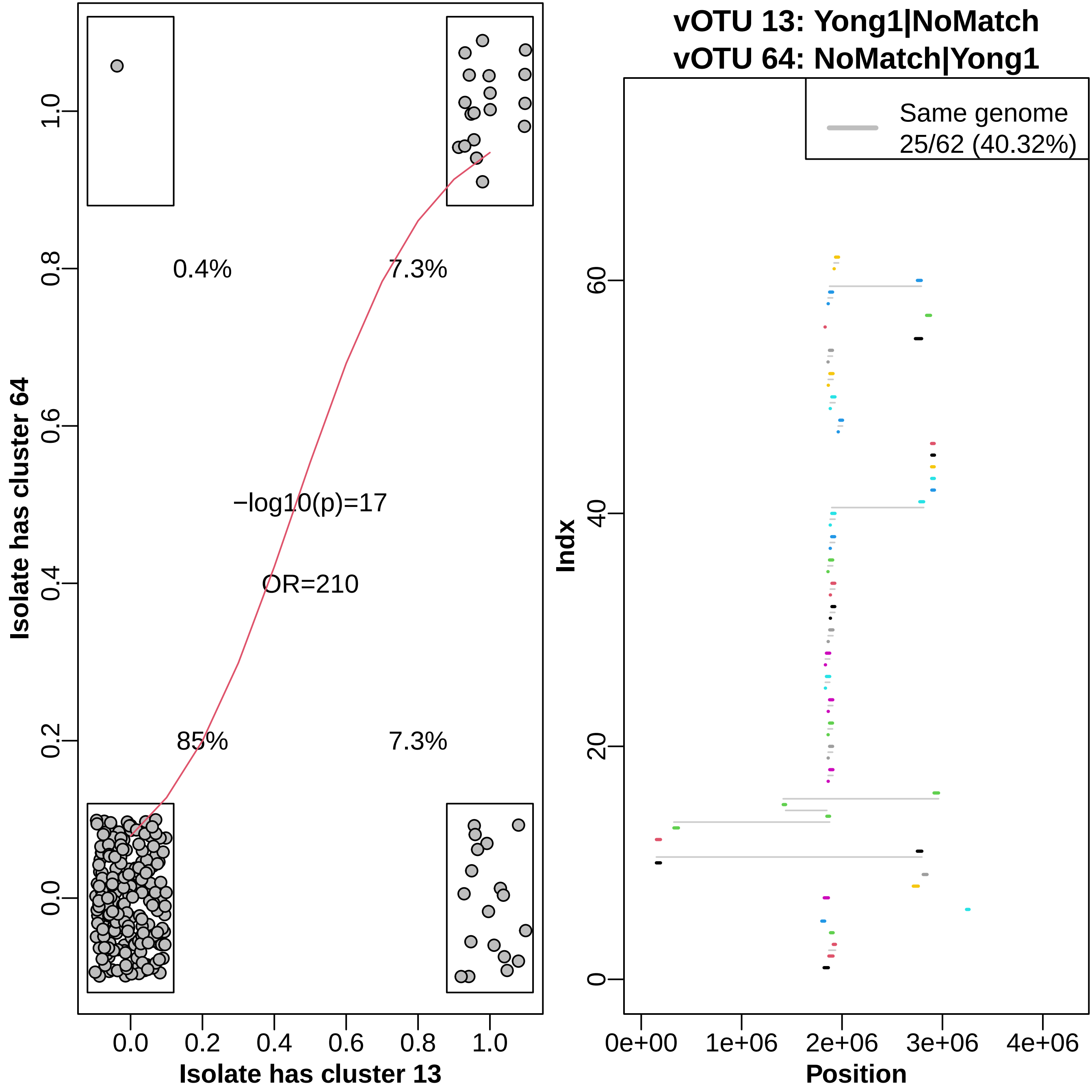

vOTU 13: Yong1|NoMatch
vOTU 64: NoMatch|Yong1
0.4%
7.3%
−log10(p)=17
OR=210
85%
7.3%
Isolate has cluster 64
Isolate has cluster 13
1.0
0.8
0.6
0.4
0.2
0.0
Same genome
25/62 (40.32%)
Indx
60
40
20
0
0e+00
1e+06
2e+06
3e+06
4e+06
Position

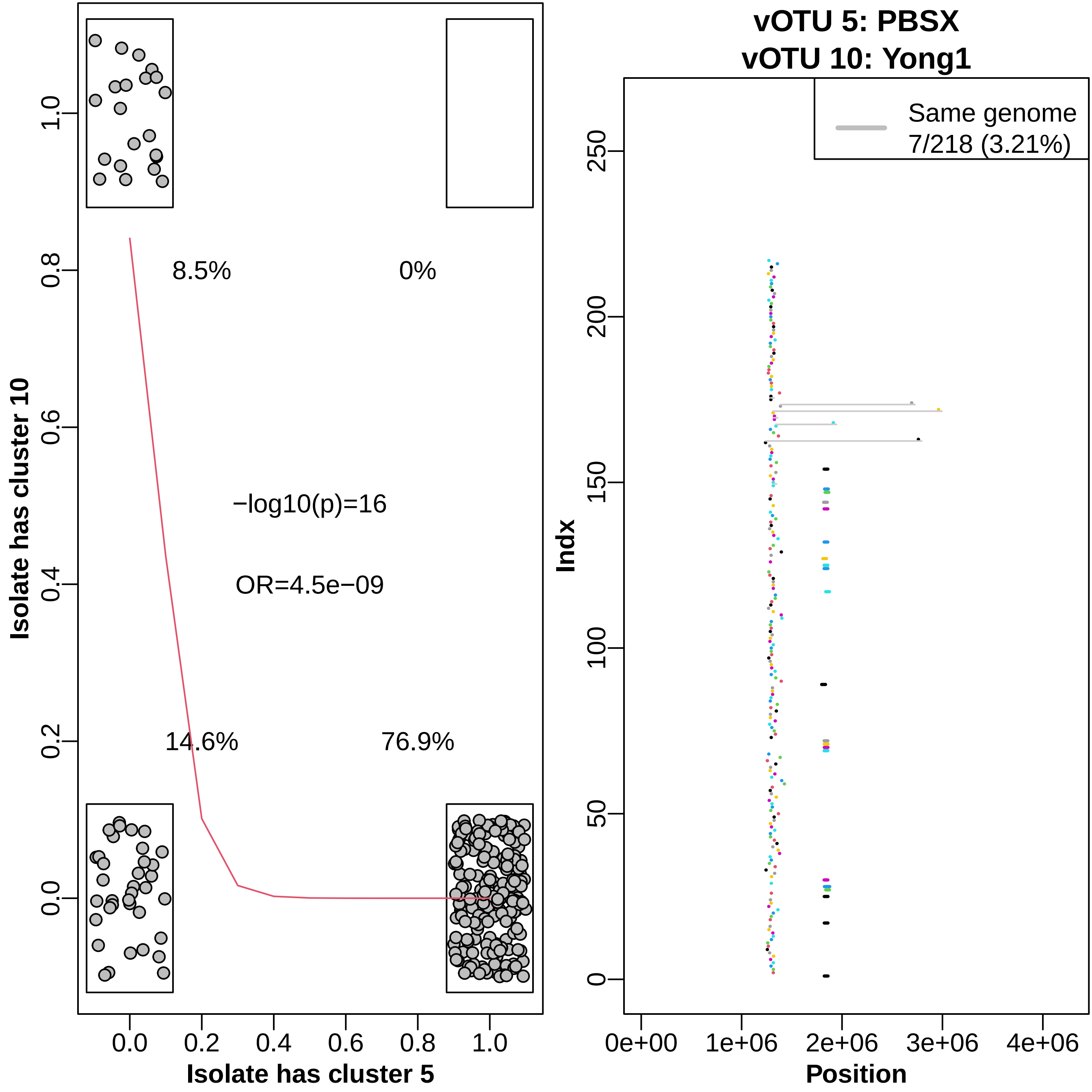

8.5%
0%
−log10(p)=16
OR=4.5e−09
14.6%
76.9%
Isolate has cluster 10
Isolate has cluster 5
vOTU 5: PBSX
vOTU 10: Yong1
Same genome
7/218 (3.21%)
Indx
Position

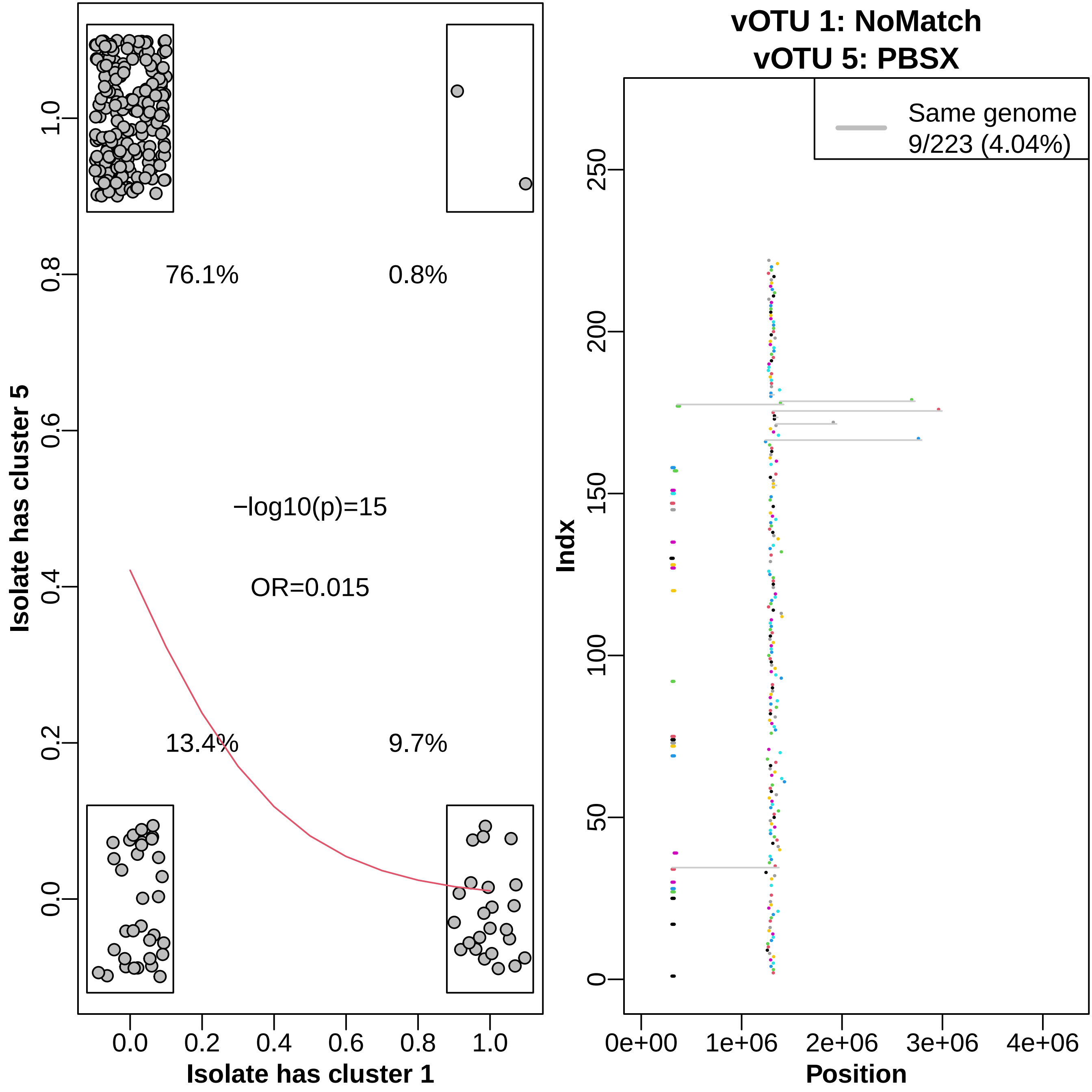

vOTU 1: NoMatch
vOTU 5: PBSX
Isolate has cluster 5
Isolate has cluster 1
76.1%
0.8%
13.4%
9.7%
−log10(p)=15
OR=0.015
Same genome
9/223 (4.04%)
Indx
Position

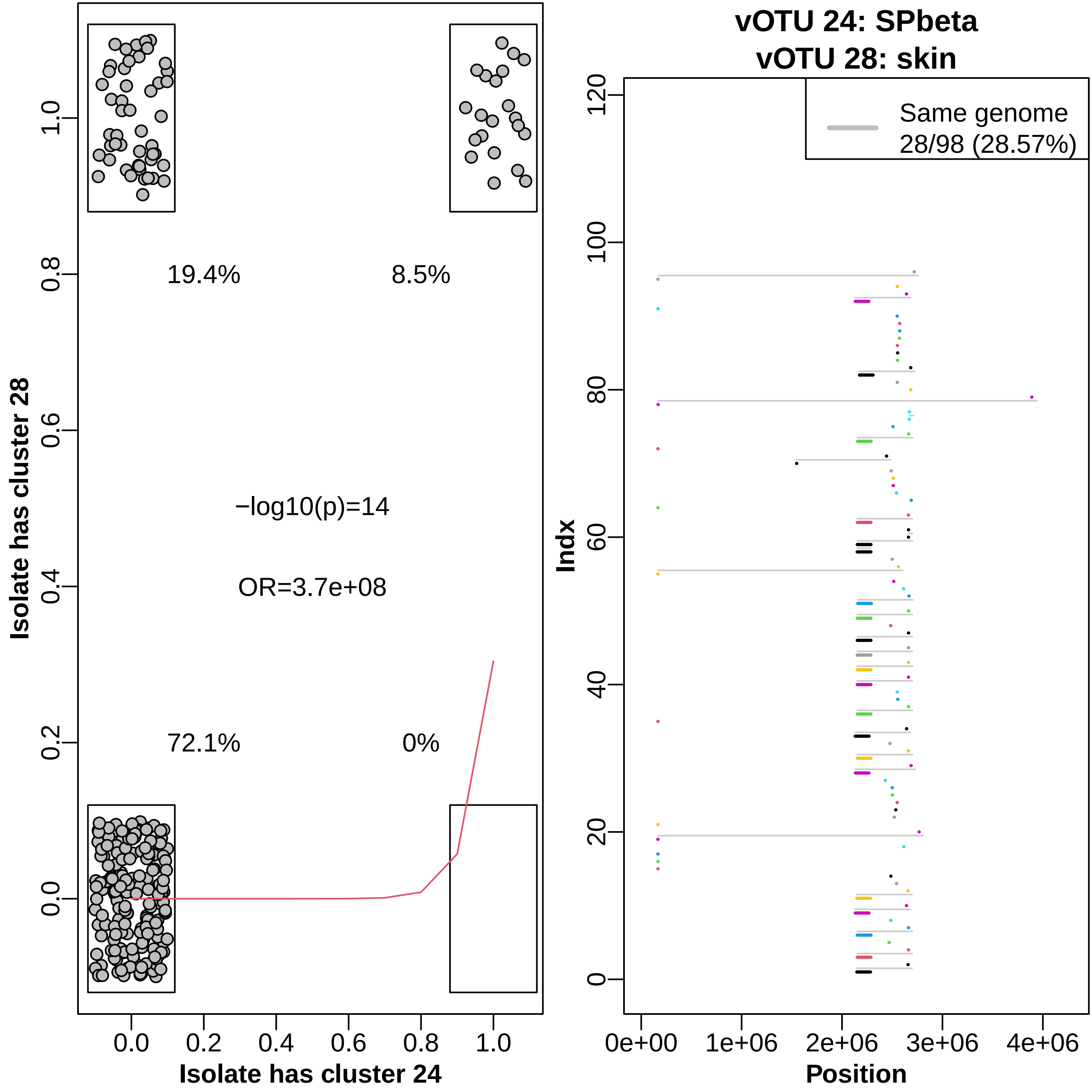

19.4%
8.5%
−log10(p)=14
OR=3.7e+08
72.1%
0%
Isolate has cluster 28
Isolate has cluster 24
vOTU 24: SPbeta
vOTU 28: skin
Same genome
28/98 (28.57%)
Indx
Position

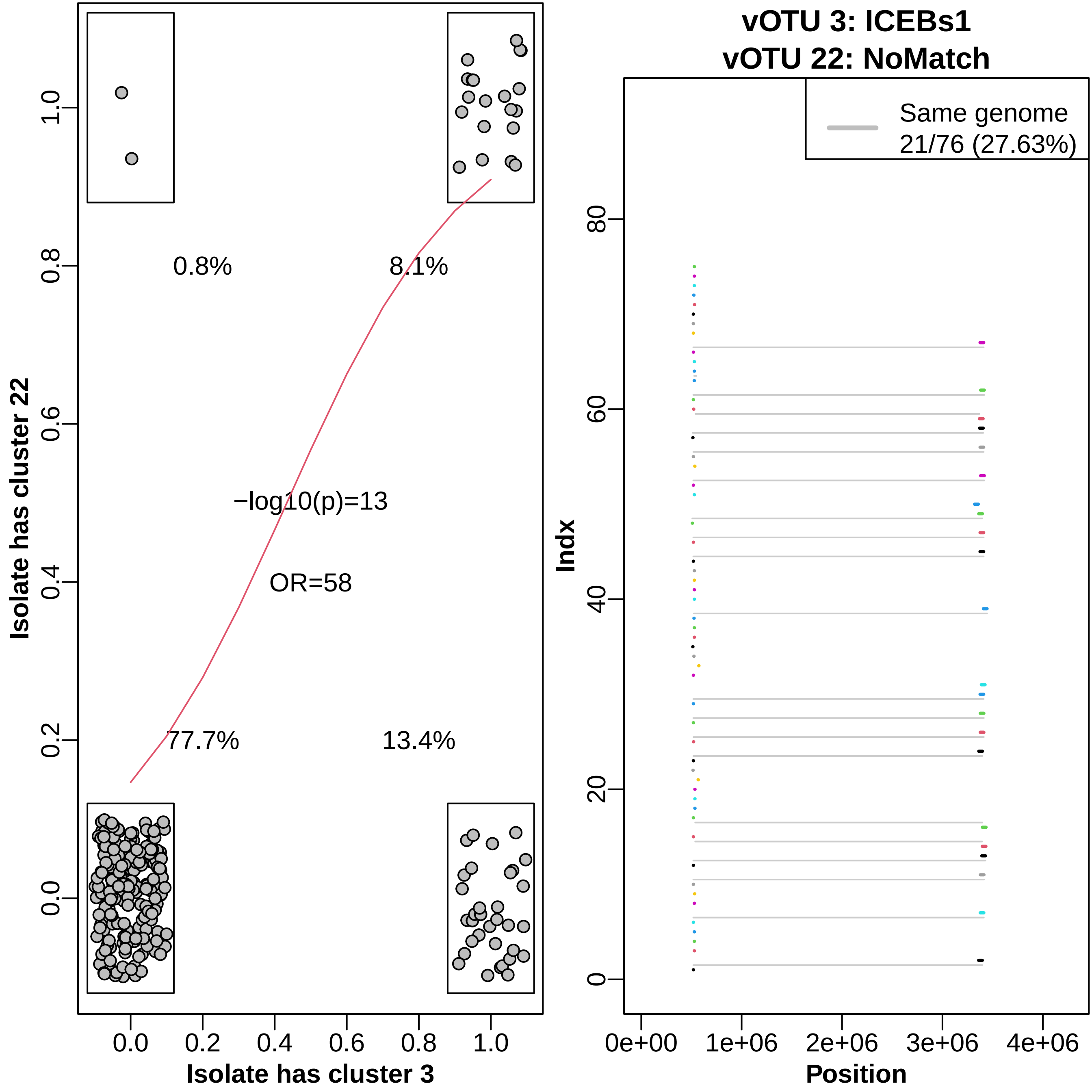

vOTU 3: ICEBs1
vOTU 22: NoMatch
0.8%
8.1%
77.7%
13.4%
−log10(p)=13
OR=58
Isolate has cluster 22
Isolate has cluster 3
0.0
0.2
0.4
0.6
0.8
1.0
Same genome
21/76 (27.63%)
Indx
Position
0e+00
1e+06
2e+06
3e+06
4e+06
0
20
40
60
80

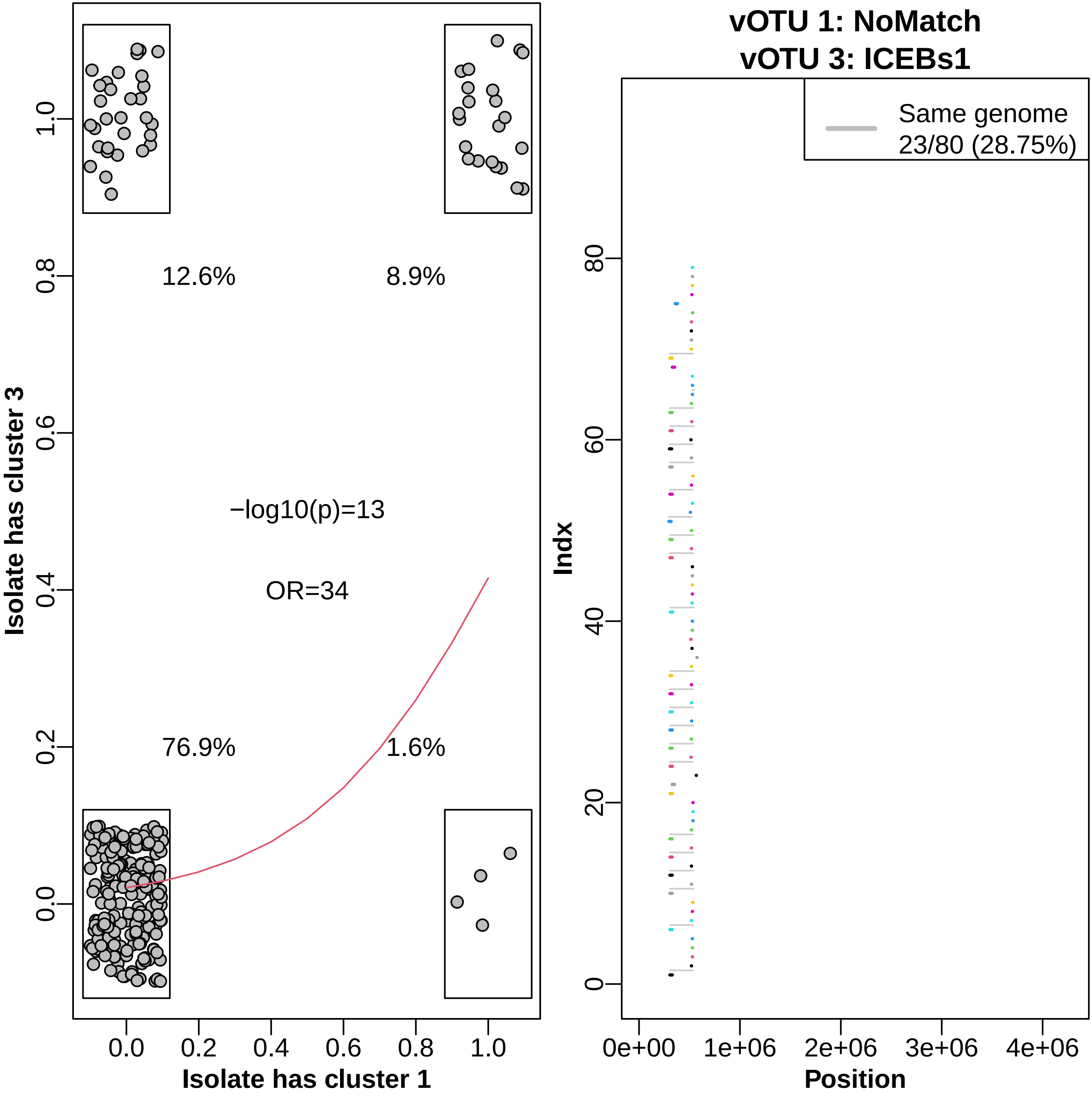

12.6%
8.9%
−log10(p)=13
OR=34
76.9%
1.6%
Isolate has cluster 3
Isolate has cluster 1
1.0
0.8
0.6
0.4
0.2
0.0
vOTU 1: NoMatch
vOTU 3: ICEBs1
Same genome
23/80 (28.75%)
Indx
80
60
40
20
0
0e+00
1e+06
2e+06
3e+06
4e+06
Position

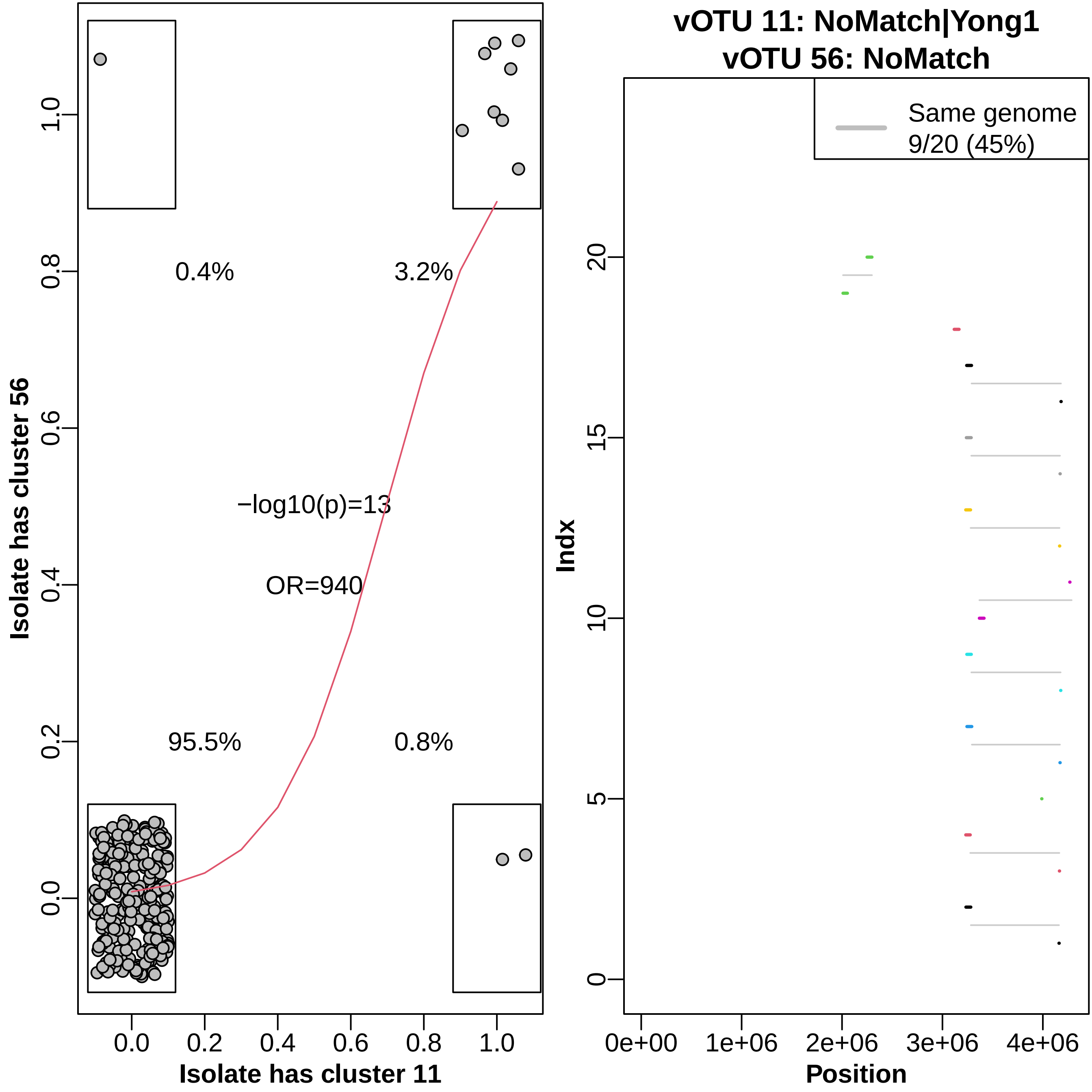

vOTU 11: NoMatch|Yong1
vOTU 56: NoMatch
Isolate has cluster 56
Isolate has cluster 11
0.4%
3.2%
95.5%
0.8%
−log10(p)=13
OR=940
0.0
0.2
0.4
0.6
0.8
1.0
Same genome
9/20 (45%)
Indx
Position
0e+00
1e+06
2e+06
3e+06
4e+06
0
5
10
15
20

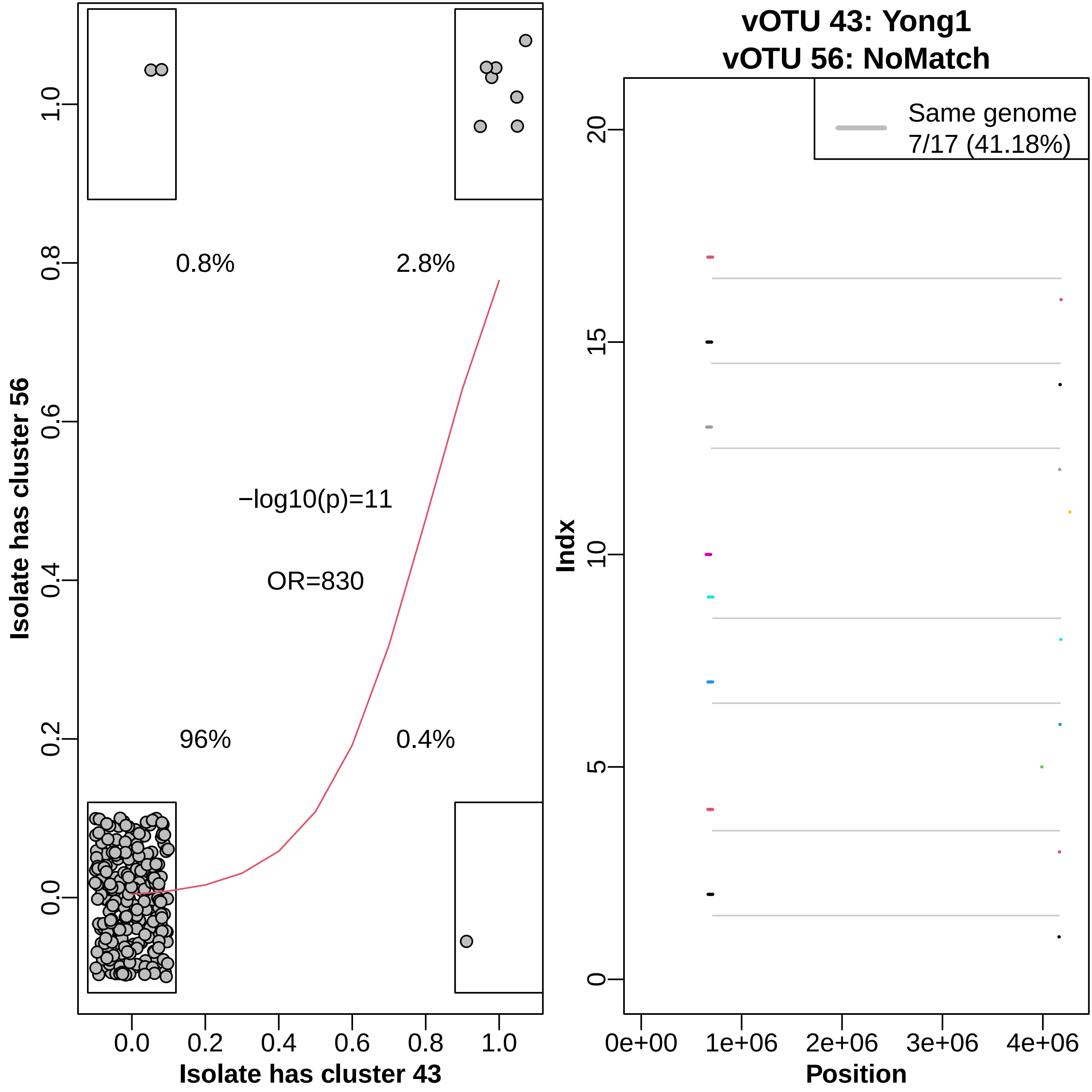

vOTU 43: Yong1
vOTU 56: NoMatch
0.8%
2.8%
−log10(p)=11
OR=830
96%
0.4%
Isolate has cluster 56
Isolate has cluster 43
0.0
0.2
0.4
0.6
0.8
1.0
Same genome
7/17 (41.18%)
Indx
Position
0e+00
1e+06
2e+06
3e+06
4e+06
0
5
10
15
20

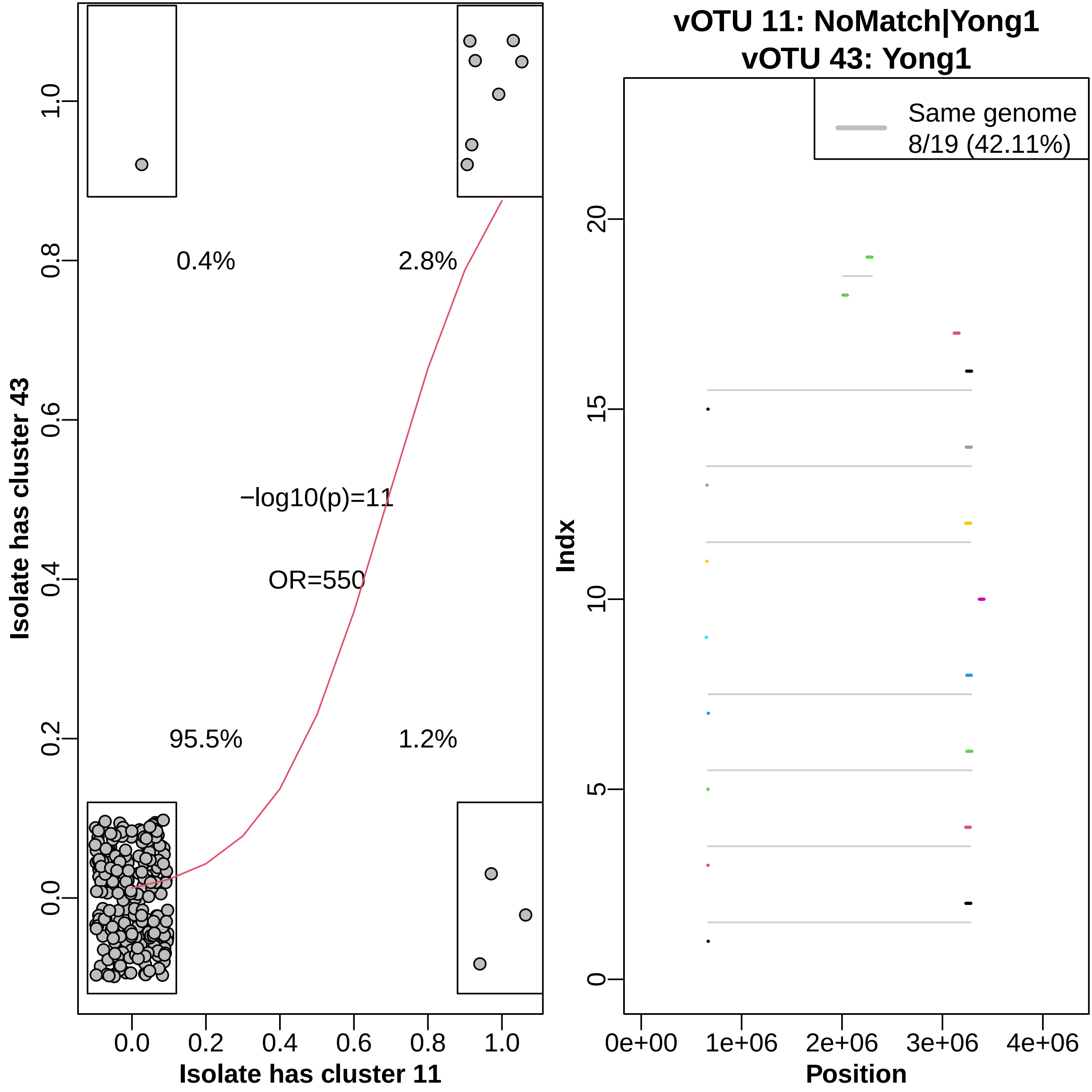

Isolate has cluster 43
Isolate has cluster 11
0.4%
2.8%
95.5%
1.2%
−log10(p)=11
OR=550
vOTU 11: NoMatch|Yong1
vOTU 43: Yong1
Same genome
8/19 (42.11%)
Indx
Position

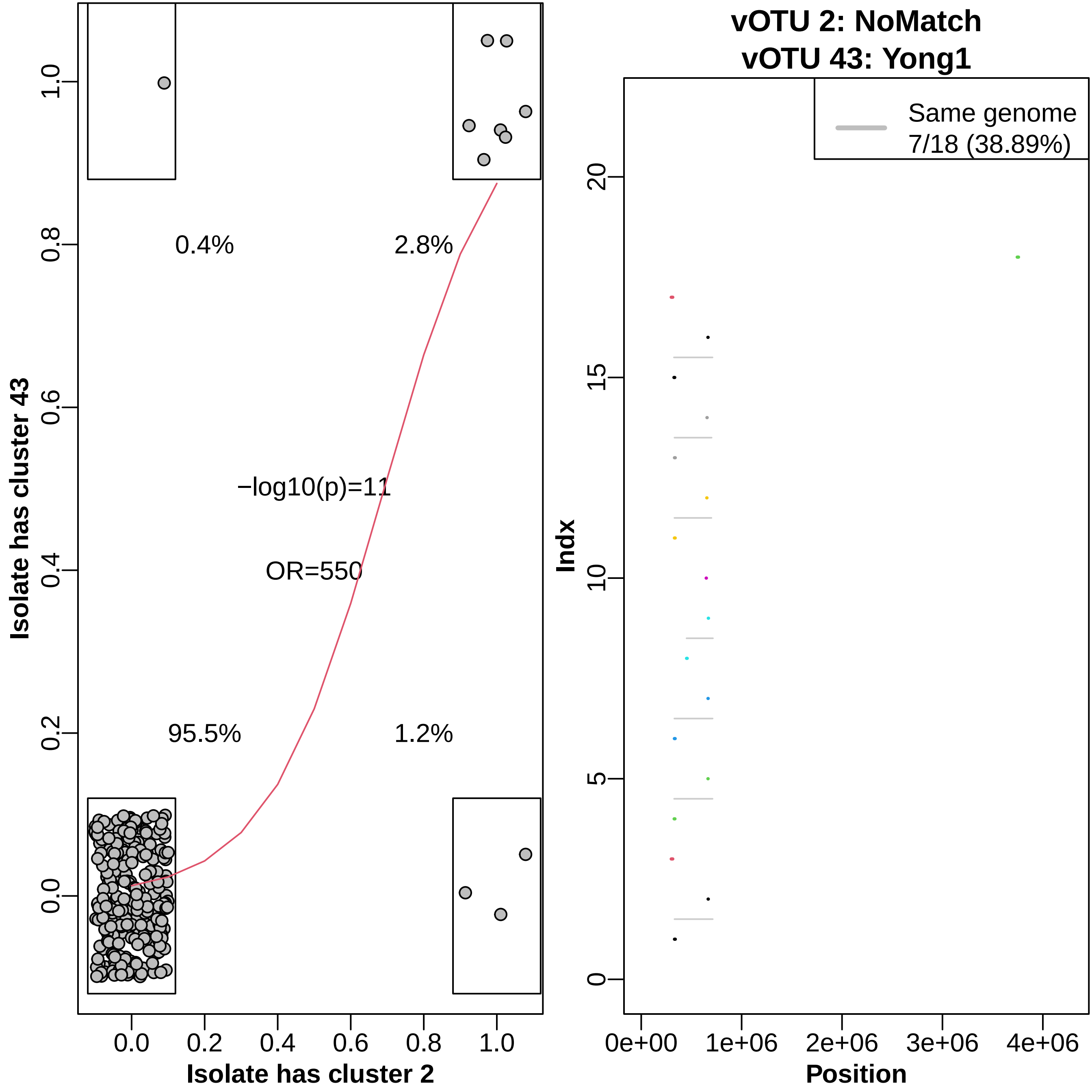

vOTU 2: NoMatch
vOTU 43: Yong1
Same genome
7/18 (38.89%)
0.4%
2.8%
−log10(p)=11
OR=550
95.5%
1.2%
Isolate has cluster 43
Isolate has cluster 2
Indx
Position
0.0
0.2
0.4
0.6
0.8
1.0
0
5
10
15
20
0e+00
1e+06
2e+06
3e+06
4e+06

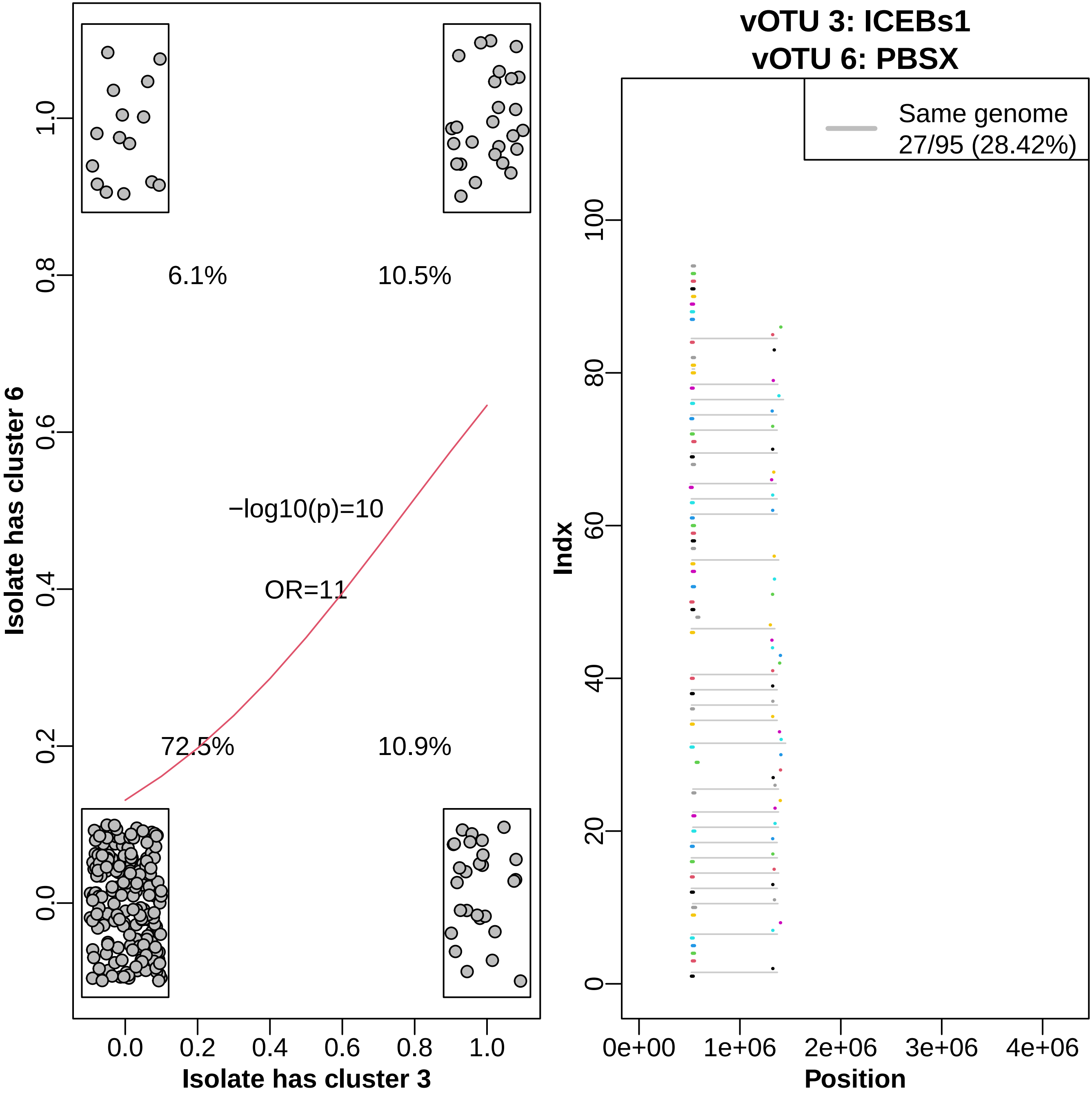

Isolate has cluster 6
Isolate has cluster 3
6.1%
10.5%
72.5%
10.9%
−log10(p)=10
OR=11
vOTU 3: ICEBs1
vOTU 6: PBSX
Same genome
27/95 (28.42%)
Indx
Position
0e+00
1e+06
2e+06
3e+06
4e+06

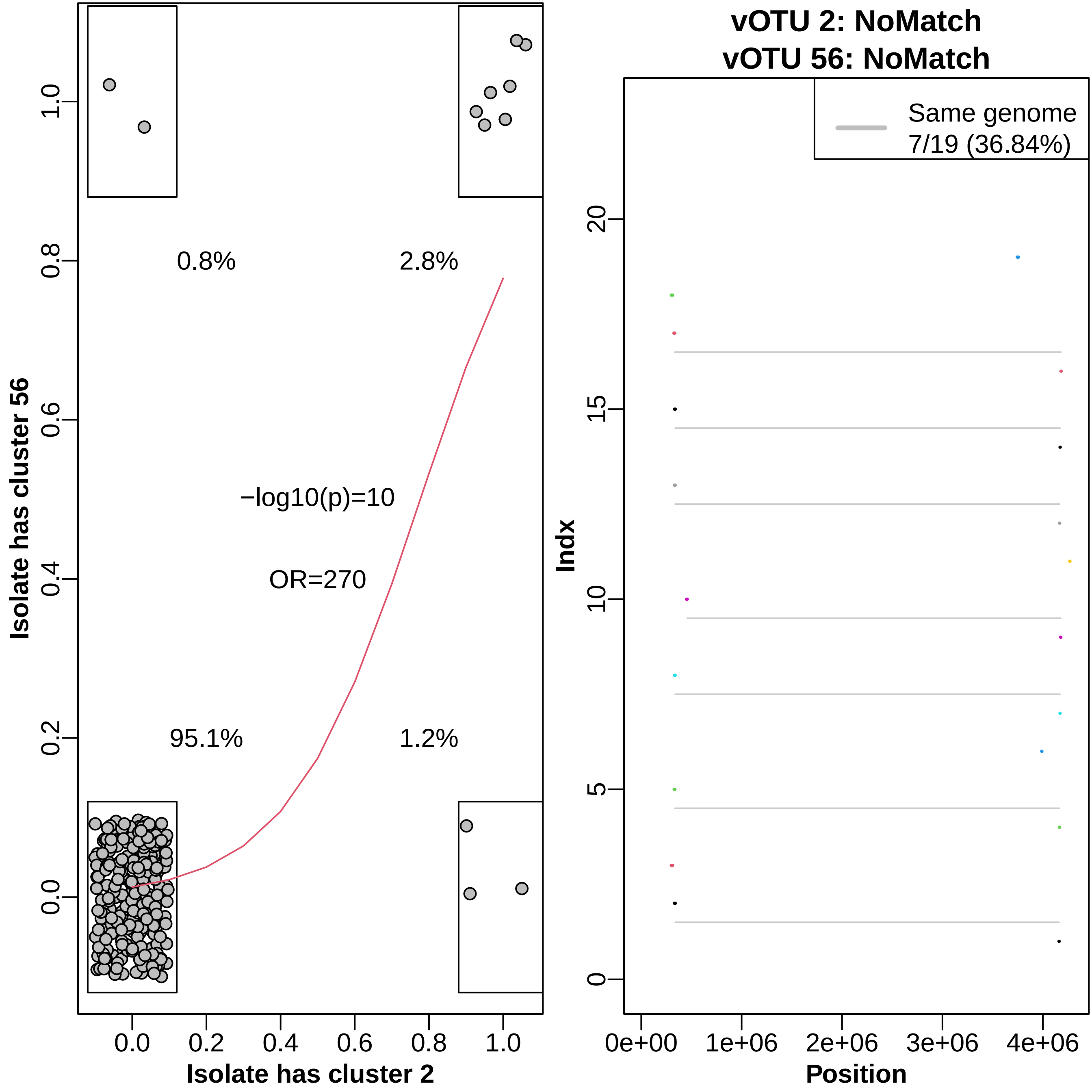

vOTU 2: NoMatch
vOTU 56: NoMatch
Same genome
7/19 (36.84%)
0.8%
2.8%
−log10(p)=10
OR=270
95.1%
1.2%
Isolate has cluster 56
Isolate has cluster 2
Indx
Position
0.0
0.2
0.4
0.6
0.8
1.0
0
5
10
15
20
0e+00
1e+06
2e+06
3e+06
4e+06
